# Molecular and Cellular Mechanisms of Teneurin Signaling in Synaptic Partner Matching

**DOI:** 10.1101/2024.02.23.581689

**Authors:** Chuanyun Xu, Zhuoran Li, Cheng Lyu, Yixin Hu, Colleen N. McLaughlin, Kenneth Kin Lam Wong, Qijing Xie, David J. Luginbuhl, Hongjie Li, Namrata D. Udeshi, Tanya Svinkina, D. R. Mani, Shuo Han, Tongchao Li, Yang Li, Ricardo Guajardo, Alice Y. Ting, Steven A. Carr, Jiefu Li, Liqun Luo

## Abstract

In developing brains, axons exhibit remarkable precision in selecting synaptic partners among many non-partner cells. Evolutionally conserved teneurins were the first identified transmembrane proteins that instruct synaptic partner matching. However, how intracellular signaling pathways execute teneurin’s functions is unclear. Here, we use *in situ* proximity labeling to obtain the intracellular interactome of teneurin (Ten-m) in the *Drosophila* brain. Genetic interaction studies using quantitative partner matching assays in both olfactory receptor neurons (ORNs) and projection neurons (PNs) reveal a common pathway: Ten-m binds to and negatively regulates a RhoGAP, thus activating the Rac1 small GTPases to promote synaptic partner matching. Developmental analyses with single-axon resolution identify the cellular mechanism of synaptic partner matching: Ten-m signaling promotes local F-actin levels and stabilizes ORN axon branches that contact partner PN dendrites. Combining spatial proteomics and high-resolution phenotypic analyses, this study advanced our understanding of both cellular and molecular mechanisms of synaptic partner matching.

**HIGHLIGHTS:**

- *In situ* spatial proteomics reveal the first intracellular interactome of teneurins
- Ten-m signals via a RhoGAP and Rac1 GTPase to regulate synaptic partner matching
- Single-axon analyses reveal a stabilization-upon-contact model for partner matching
- Ten-m signaling promotes F-actin in axon branches contacting partner dendrites

## INTRODUCTION

The precise assembly of neural circuits involves multiple developmental processes. Compared to axon guidance and dendrite morphogenesis, much less is known about cellular and molecular mechanisms that mediate synaptic partner matching^1–4^. Evolutionarily conserved teneurins are among the first identified transmembrane proteins that instruct synaptic partner matching^5,6^. Teneurins also regulate diverse other biological processes including cell polarity, neuronal migration, axon guidance and target selection, axon myelination, and synapse development^6–17^. All four human teneurins have been implicated in a variety of diseases including sensory dysfunctions, movement disorders, neurodevelopmental and psychiatric disorders, and cancers^18–26^.

Teneurins are type II transmembrane proteins that comprise a small intracellular amino terminus, a single transmembrane domain, and a large extracellular carboxyl terminus with evolutionarily conserved domains for protein-protein interactions^27^ (**Figure 1J**). Previous structural and functional studies of teneurins have largely focused on the cell-cell interactions mediated by their extracellular domains, which include EGF-like repeats essential for teneurin *cis*-dimerization, a beta-propeller region implicated in *trans*-homophilic binding, a tyrosine and glutamate rich YD domain for heterophilic interactions with latrophilins, members of adhesion G-protein-coupled receptor family^9,28–36^. For example, homophilic attractions between mouse teneurin-3 regulate topographic target selection of hippocampal axons^13^, whereas heterophilic interactions between teneurin-3 and latrophilin-2 mediate reciprocal repulsions between axons and target neurons that express them^14,37^. Heterophilic interactions between teneurins and latrophilins also regulate neuronal migration^9^ and synapse formation in specific subcellular compartments^15^. However, compared to our rich knowledge of the extracellular domains, little is known about how intracellular signaling works to execute the diverse functions triggered by extracellular interactions of teneurins. Indeed, it is unknown whether intracellular domains are required for any of tenerins’ functions.

**Figure 1.**
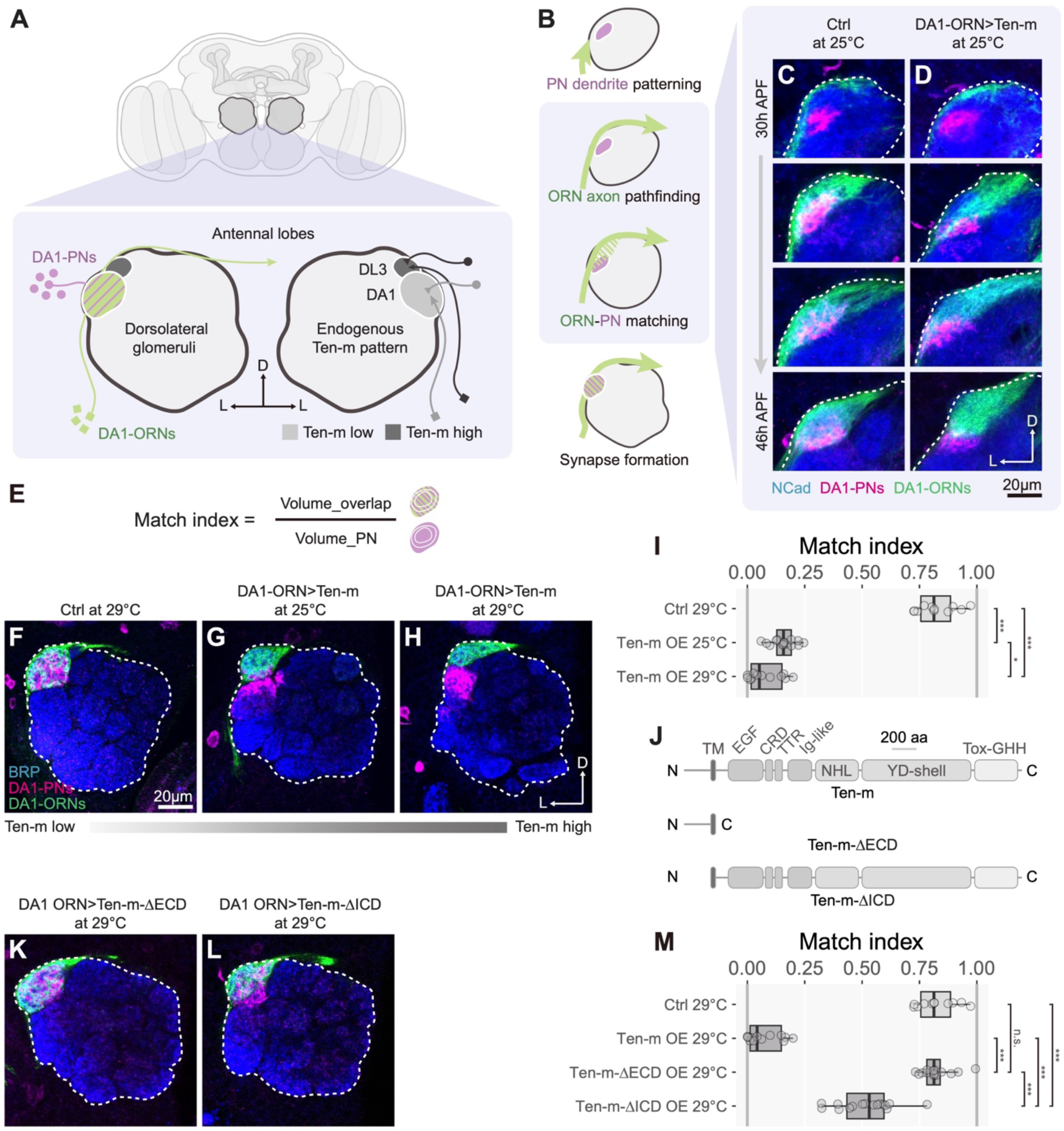
A quantitative gain-of-function assay for synaptic partner matching reveals the requirement of both the extracellular and intracellular domains of Ten-m for signaling. (A) Schematic of the adult *Drosophila* brain highlighting a pair of antennal lobes and locations of DA1 and DL3 glomeruli. Left, DA1-ORN axons (green) synapse with the corresponding DA1-PN dendrites (purple, contralateral projection omitted for simplicity). Right, endogenous Ten-m levels are low in DA1-ORNs and PNs, but high in DL3-ORNs and PNs. (B) Schematic of sequential developmental steps of DA1 ORN-PN pairing. (C) Time course of control DA1-ORN axons (green, labeled by a plasma membrane marker mouse CD8-GFP (mCD8-GFP) driven by a spilt GAL4 consisted of *R22E04-GAL4^DBD^* and *VT028327-p65^AD^*, hereafter DA1-ORN GAL4) innervating, elaborating, and coalescing with DA1-PN dendrites (magenta, labeled by a membrane-tagged tdTomato, driven by *Mz19-QF2^G4HACK^*, hereafter *Mz19-QF2).* APF, after puparium formation. (D) Time course of DA1-ORN axons with Ten-m overexpression elaborating at a more dorsomedial region, eventually resulting in only partial overlap between DA1-ORN axons and DA1-PN dendrites. (E) “Match index” is defined as the ratio of the overlapping volume between DA1-ORN axons and DA1-PN dendrites to the total volume of DA1-PN dendrites. (F–H) Confocal optical sections of adult antennal lobes showing DA1-ORN axons (green) of control (F), Ten-m overexpression at 25°C (G) and 29°C (H), as well as DA1-PN dendrites (magenta). (I) Match indices for experiments in panels F–H. (J) Domain organization of Ten-m, Ten-m-ΔECD, and Ten-m-ΔICD. TM, transmembrane domain; EGF, epidermal growth factor repeat; CRD, cysteine-rich domain; TTR, transthyretin-related domain; Ig-like, immunoglobulin-like domain; NHL, a domain named after homology between NCL-1, HT2A, and Lin-41, also called β-propeller domain; YD-shell, enriched for tyrosine and aspartate, also called β-barrel domain; Tox-GHH, toxin-like domain; aa, amino acids. (K, L) Confocal optical sections of adult antennal lobes showing DA1-ORN axons (green) of Ten-m-ΔECD overexpression (K) or Ten-m-ΔICD overexpression (L) at 29°C, as well as DA1-PN dendrites (magenta). (M) Match indices for experiments in K and L. D, dorsal; L, lateral. Dashed white circle, antennal lobe. NCad, N-cadherin, a general neuropil marker; BRP, Bruchpilot, an active zone marker used for general neuropil staining. The one-way ANOVA (with Tukey’s test) was used for multiple comparisons. In this and all subsequent figures, * p < 0.05; ** p < 0.01; *** p < 0.001; n.s., not significant.

In the *Drosophila* olfactory circuit, about 50 types of olfactory receptor neurons (ORNs) synapse with 50 types of second-order projection neurons (PNs) to form precise 1-to-1 matching at 50 discrete glomeruli (**Figure 1A**), providing an excellent model for investigating mechanisms of synaptic partner matching. We previously found that two *Drosophila* teneurins, Ten-m (tenascin-major) and Ten-a (tenascin-accessory), are each expressed in select matching ORN–PN pairs and instruct ORN–PN synaptic partner matching through homophilic attraction^5^. In this study, we combine spatial proteomics and *in vivo* genetic interaction assays to investigate the intracellular signaling mechanisms that mediate this attraction. We find that Ten-m signals through a RhoGAP and the Rac1 small GTPase to regulate the actin cytoskeleton. Developmental analyses with single-axon resolution further reveal that this signaling pathway acts to selectively stabilize ORN axon branches that contact partner PN dendrites.

## RESULTS

### A quantitative gain-of-function assay for Ten-m signaling *in vivo*

To investigate Ten-m signaling mechanisms, we first sought to establish a quantitative assay in which altering Ten-m activity would lead to a robust phenotype *in vivo*. We can then examine how perturbing Ten-m’s signaling partner(s) would enhance or suppress such a phenotype. We focused on DA1-ORNs that target their axons to the DA1 glomerulus where they synapse with the dendrites of DA1-PNs (**Figure 1A**, bottom left; **Figure S2A**). Both DA1-ORNs and DA1-PNs express Ten-m at low levels^5^ (**Figure 1A**, bottom right). By utilizing a selective driver to genetically access DA1-ORNs and an orthogonal driver for labeling DA1-PNs with a distinct fluorophore, we simultaneously tracked axons and their partner dendrites across development in the same control (**Figure 1C**) or Ten-m overexpressing (**Figure 1D**) animals.

During fly olfactory circuit assembly, PNs first pattern the antennal lobe by targeting their dendrites to the approximate regions of the antennal lobe corresponding to their eventual glomerular positions (**Figure 1B**)^38,39^. At 30 hours after puparium formation (h APF), DA1-ORN axons extended along the antennal lobe surface without forming extensive contact with DA1-PN dendrites in both control and Ten-m overexpression conditions. During the next 16 hours, control DA1-ORN axons initially elaborated over a larger region than DA1-PN dendrites, and then gradually coalesced their axons with DA1-PN dendrites (**Figure 1C**). However, DA1-ORN axons overexpressing Ten-m elaborated over a region more dorsomedial than the region occupied by DA1-PN dendrites, resulting in only partial overlap between DA1-ORN axons and DA1-PN dendrites even at later developmental stages (**Figure 1D**).

To quantify the Ten-m overexpression–induced mismatching phenotype in DA1-ORNs, we devised a “match index” as the volume in which DA1-ORN axons and DA1-PN dendrites overlap divided by the total volume of DA1-PN dendrites in the adult antennal lobe (**Figure 1E**). We found a substantial difference in the match index between the control and Ten-m overexpression conditions (**Figure 1F, H**; quantified in **Figure 1I**). To determine whether this mismatching phenotype depends on Ten-m overexpression levels in DA1-ORNs, we exploited the temperature-dependent increase in GAL4-driven transgene expression^40,41^ (**Figure S1A–C, S1F)**, and observed a more pronounced Ten-m overexpression-induced phenotype at 29°C than at 25°C (**Figure 1G, H**; quantified in **Figure 1I**). Thus, the match index provides an assay sensitive to Ten-m overexpression levels.

Using *trans-*synaptic labeling^42^, we found that mistargeted DA1-ORN axons likely matched with dendrites of DL3-PNs, based on the location of the labeled postsynaptic PN dendrites and axonal projection patterns (**Figure S2**). These results further underscore the role of like-to-like matching in teneurin levels between synaptic partners as DL3-PNs express high levels of both Ten-m and Ten-a^5^, paralleling Ten-m-overexpressing DA1-ORNs that normally express high levels of Ten-a^5^. In addition, co-overexpressing Ten-m in DA1-PNs could partially suppress the mismatching phenotypes caused by overexpressing Ten-m in DA1-ORNs (**Figure S1G, H**). Thus, the gain-of-function phenotypes we observed likely result from homophilic attraction between Ten-m-expressing ORNs and PNs.

### Both the extracellular and intracellular domains of Ten-m are required for signaling

Using our quantitative assay, we assessed the role of the extracellular and intracellular domains of Ten-m in mediating signaling by overexpressing *Ten-m* transgenes lacking the extracellular domain (ΔECD) or the intracellular domain (ΔICD) (**Figure 1J**). These truncated mutants were integrated into the same genomic locus as the full-length *Ten-m* transgene, expressed at a similar level as full-length Ten-m proteins *in vivo* (**Figure S1D–F**), and were trafficked to the cell surface (**Figure S1I, J**). Interestingly, overexpression of Ten-m-ΔECD did not cause any mismatching phenotype (**Figure 1K**, quantified in **Figure 1M**), while overexpression of Ten-m-ΔICD exhibited a partial mismatching phenotype (**Figure 1L**, quantified in **Figure 1M**). These experiments indicate that the extracellular domain of Ten-m is essential for mediating its gain-of-function effect. Signaling through the intracellular domain is also required for the full activity of Ten-m; the remaining mismatching phenotypes induced by overexpressing Ten-m-ΔICD could be caused by homophilic adhesion between DA1-ORNs and non-partner PNs without intracellular signaling, or by a potential co-receptor of Ten-m that can mediate some intracellular signaling. Regardless, the substantial difference between overexpressing wild-type Ten-m and Ten-m-ΔICD in the match index offers a quantitative assay for examining the signaling mechanism that depends on the intracellular domain of Ten-m.

### Proximity labeling to identify Ten-m-ICD interacting proteins *in situ*

We next investigated the molecular mechanisms by which the intracellular domain of Ten-m transduces signals. As cell-surface signaling often involves transient binding events and requires a physiological membrane environment for full biological activity, many conventional methods for proteomic profiling are not effective^43^. We thus used proximity labeling^43–45^ to identify proteins in physical proximity to Ten-m-ICD, including both stable and transient partners, in native tissues. Given the critical role of teneurin levels in synaptic matching, we used CRISPR knockin rather than transgene overexpression to maintain endogenous Ten-m levels. Specifically, we inserted the coding sequence of *APEX2-V5* N-terminal to the beginning of the *Ten-m* coding sequence (**Figure 2A**) such that APEX2 would catalyze the addition of biotin to proteins in physical proximity to Ten-m-ICD in the presence of biotin-phenol and H_2_O_2_ (**Figure 2B**). Flies homozygous for the insertion allele were viable, whereas flies homozygous for *Ten-m* mutant are embryonic lethal^11^, suggesting that APEX2-V5 insertion did not disrupt native Ten-m function. APEX2-V5-Ten-m expression patterns from the knockin allele recapitulated that of the endogenous Ten-m, as exemplified by high-level expression in the VA1d and VA1v glomeruli and low-level expression in the DA1 glomerulus (**Figure 2C, E, E’’**), consistent with our previous study^5^. Furthermore, in the presence of biotin-phenol and H_2_O_2_, APEX2-V5-Ten-m catalyzed biotinylation with a similar spatial pattern as V5 staining (**Figure 2C, C’, E, E’**); no biotinylation was observed when H_2_O_2_ was omitted (**Figure 2D, D’**).

**Figure 2.**
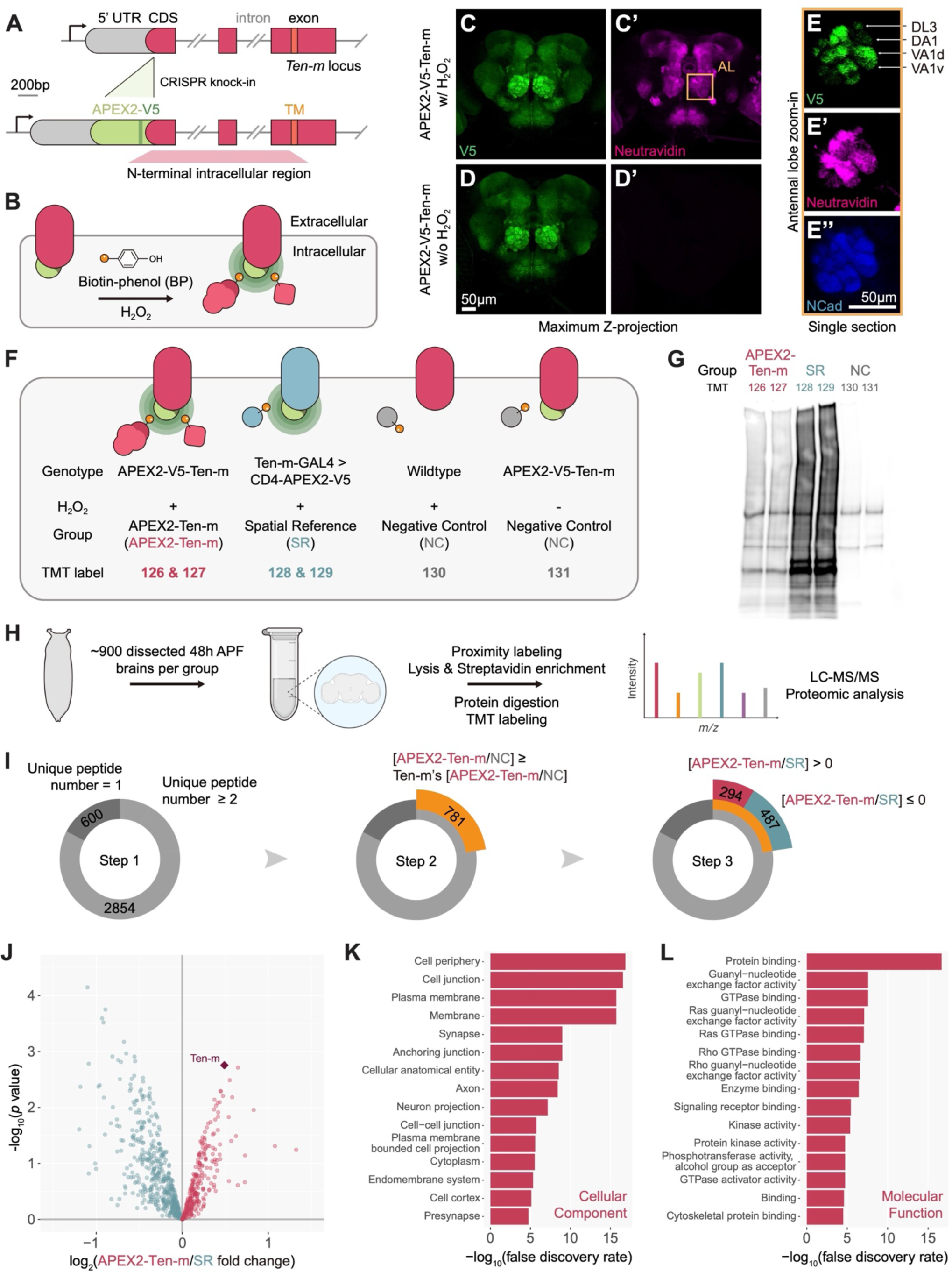
*In situ* spatial proteomics to identify proteins in physical proximity of the Ten-m intracellular domain. (A) CRISPR knockin at the *Ten-m* gene locus. APEX2-V5 is N-terminal to the coding sequence (CDS) of Ten-m. TM, transmembrane domain. (B) Schematic of APEX2-based *in situ* proximity labeling for profiling of the Ten-m intracellular interactome. Extracellular domain size is not in scale. (C and C’) V5 and Neutravidin staining of APEX2-V5-Ten-m fly brain after proximity labeling. (D and D’) V5 and Neutravidin staining of APEX2-V5-Ten-m fly brain without H_2_O_2_. (E–E’’) Single optical section of the antennal lobe showing that Ten-m expression and APEX2 activity are high in the DL3, VA1d, and VA1v glomeruli but low in the DA1 glomerulus. NCad, N-cadherin, a general neuropil marker. (F) Design of the 6-plex tandem mass tag (TMT6)-based quantitative proteomic experiment. Labels in the TMT row (e.g., 126) indicate the TMT tags used for each of the three groups: APEX2-V5-Ten-m labeling group (APEX2-Ten-m, red, two replicates; TMT 126 and 127); spatial reference group (SR, blue, two replicates; TMT 128 and 129); and negative control group (NC, gray, omitting APEX2 transgenes or H_2_O_2_; TMT 130 and 131). (G) Streptavidin blot of the post-enrichment bead elute. (H) Workflow of the Ten-m intracellular interactome profiling. (I) Numbers of proteins after each step of the ratiometric and cutoff analysis. (J) Volcano plot showing proteins passing the step 1 and 2 filters. Based on the TMT ratio of Ten-m labeling (APEX2-Ten-m) and spatial reference (SR) groups, spatially enriched Ten-m intracellular interactome (log_2_[APEX2-Ten-m/SR] > 0) is colored red while the other proteins (log_2_[APEX2-Ten-m/SR] ≤ 0) are colored blue. Diamond, Ten-m protein. (K) Top 15 cellular component Gene Ontology terms of the Ten-m intracellular interactome. (L) Top 15 molecular function Gene Ontology terms of the Ten-m intracellular interactome.

We next carried out large-scale proximity labeling experiments from pupal brains followed by quantitative mass spectrometry to identify proteins that interact with Ten-m-ICD during development. We devised a 6-plex tandem mass tag (TMT) design for ratiometric analysis, featuring an APEX2-V5-Ten-m labeling group (APEX2-Ten-m group, 2 replicates, to capture Ten-m-ICD interactors), a spatial reference group (SR group, 2 replicates, to identify the background from generic proteins close to the plasma membrane), and a negative control group (NC group with 2 samples, omitting either H_2_O_2_ or any APEX2 transgene, to identify the background from endogenously biotinylated and endogenous peroxidase-labeled proteins) (**Figure 2F**). For the SR group, CD4-APEX2-V5—a generic transmembrane protein (CD4) with APEX2 at its intracellular C-terminus—was expressed in Ten-m-expressing cells (**Figure 2F**). Biochemical characterization of the post-enrichment eluate via streptavidin blot analysis revealed that both the APEX2-Ten-m and spatial reference groups had much more biotinylated proteins, each with a distinct pattern, than the negative control group, indicating group-specific protein enrichment (**Figure 2G**).

We dissected ∼900 brains at 48h APF per TMT plex (∼5400 brains for this 6-plex experiment) and processed the samples using a protocol similar to previously described ones^46,47^. In brief, we pre-incubated freshly dissected brains of each sample with the biotin-phenol substrate before 1 min H_2_O_2_ incubation to catalyze proximity labeling. We then separately lysed each sample and enriched them using streptavidin beads. After on-bead trypsin digestion and 6-plex TMT labeling, we pooled all samples for liquid chromatography-tandem mass spectrometry (LC-MS/MS) analysis (**Figure 2H**). All proteomes exhibited strong correlations between biological replicates (**Figure S3A**), suggesting high sample quality.

To identify prospective interacting partners of Ten-m, we applied 3 steps in our proteomic analysis. (1) We filtered a total of 3454 detected proteins from 6 samples, focusing on those with two or more unique peptides, resulting in 2854 proteins (**Figure 2I**, Step 1; **Table S1**). (2) To remove potential contaminants such as endogenously biotinylated and endogenous peroxidase-labeled proteins (as revealed by biotinylated proteins in the negative control lanes in **Figure 2G**), we used [APEX2-Ten-m/NC] fold change of the Ten-m protein itself (**Figure 2F**) as a cutoff and obtained 781 proteins (**Figure 2I**, Step 2; **Table S1**). (3) To remove generic proteins close to the cell membrane, we applied a [APEX2-Ten-m/SR] fold change-based ratiometric strategy (**Figure 2F**) and acquired 294 proteins enriched by APEX2-Ten-m (**Figure 2I**, Step 3; **Figure 2J**, red-color coded; **Table S1**)—hereafter, the Ten-m intracellular interactome.

Gene Ontology analysis suggested features of the cellular components and molecular functions of the Ten-m intracellular interactome. Cellular component terms indicated that the Ten-m interactome consisted of proteins localized at the cell surface, synapse, cytoplasm, and endomembrane systems (**Figure 2K**). Furthermore, these proteins functionally relate to GTPase signaling pathways, kinase activity, signaling receptor binding, and cytoskeletal protein binding (**Figure 2L**, **Figure S3B–D**), reminiscent of previously identified axon guidance receptors such as Eph receptors^48,49^ and Plexins^50–52^.

### Ten-m binds to and genetically interacts with Syd1, a GAP for Rho GTPases

Among 37 proteins that were significantly enriched in the APEX2-Ten-m group relative to the spatial reference group (**Figure 3A**; **Table S2**) was RhoGAP100F (Syd1), the *Drosophila* homolog of *C. elegans* Syd-1, which functions in the assembly of presynaptic terminals in worms and flies^53,54^. Syd1 has a GTPase activating protein (GAP) domain for the Rho family of small GTPases (**Figure 3B**) and has been shown to exhibit GAP activity towards Rac1 and Cdc42^55^. Given the central role for Rho GTPases in transducing extracellular signals to the cytoskeleton^56^ (**Figure 3C**), we next investigated the interactions between Ten-m and Syd1.

**Figure 3.**
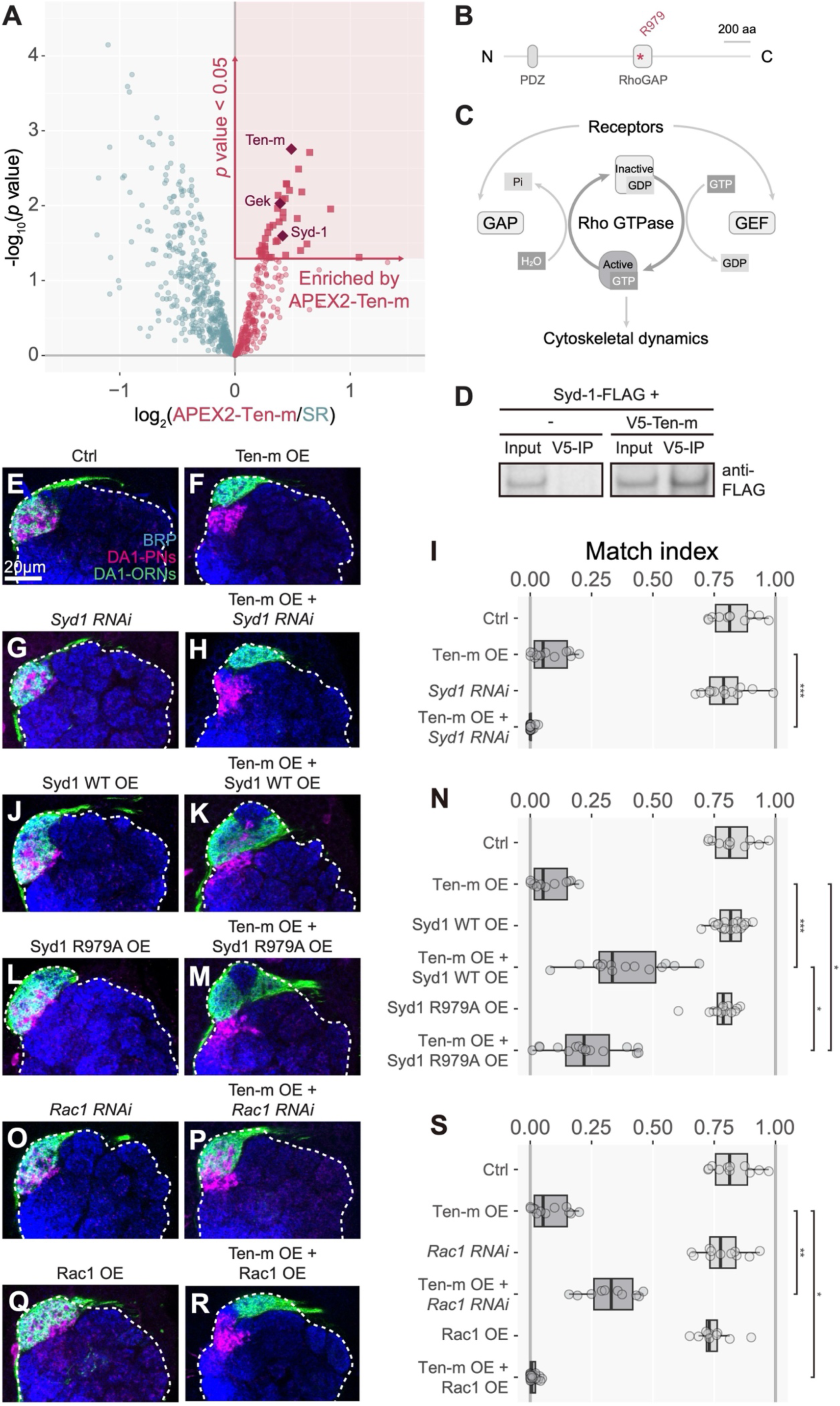
Ten-m interacts with the RhoGAP Syd1 and Rac1 GTPase in ORNs. (A) Selection criteria for the top candidate Ten-m-interacting proteins: log_2_[APEX2-Ten-m/SR] > 0 and *p* value < 0.05. Ten-m, Syd1, and Gek are highlighted with diamond. (B) Domain organization of Syd1. PDZ, PDZ domain (named after PSD95, Dlg1, and ZO-1); RhoGAP, Rho GTPase-activating protein domain; aa, amino acids. The asterisk (*) marks the active site in the RhoGAP domain. (C) Diagram illustrating the Rho GTPase cycle. Upstream signals (e.g., from receptors on the plasma membrane) control Rho GTPases through regulating guanine nucleotide exchange factors (GEFs) or GTPase-activating proteins (GAPs), which facilitate switching the Rho GTPase on or off, respectively. In the GTP-bound state, the Rho GTPase binds to and activates its effectors to regulate cytoskeletal dynamics. (D) Co-immunoprecipitation of V5-tagged Ten-m and FLAG-tagged Syd1 proteins from co-transfected S2 cells. (E–I) Representative confocal images of DA1-PN dendrites (magenta) and DA1-ORN axons (green) of control (E), Ten-m overexpression (F), *Syd1-RNAi* alone (G), and Ten-m overexpression with *Syd1-RNAi* (H). Match indices are quantified in (I). (J–N) Representative confocal images of DA1-PN dendrites (magenta) and DA1-ORN axons (green) of Syd1 overexpression (J), Syd1 and Ten-m co-overexpression (K), Syd1-R979A GAP-domain mutation overexpression (L), and Syd1-R979A and Ten-m co-overexpression (M). Match indices are quantified in (N). (O–S) Representative confocal images of DA1-PN dendrites (magenta) and DA1-ORN axons (green) of *Rac1-RNAi* (O), Ten-m overexpression with *Rac1-RNAi* (P), Rac1 overexpression (Q), and Ten-m and Rac1 co-overexpression (R). Match indices are quantified in (S). D, dorsal; L, lateral. Dashed white circle, antennal lobe. BRP, Bruchpilot, an active zone marker used for general neuropil staining. The Kruskal-Wallis test with Bonferroni post-hoc correction for multiple comparisons was used in (I), (N), and (S).

To test whether Ten-m physically interacts with Syd1, we transfected S2 cells with expression constructs for V5-tagged full-length Ten-m and FLAG-tagged full-length Syd1. Immunoprecipitation of S2 cell extracts with a V5 antibody co-precipitated Syd1-FLAG (**Figure 3D**), indicating that Ten-m and Syd1 directly interact or belong to a same protein complex.

To test whether Ten-m genetically interacts with Syd1, we utilized our Ten-m overexpression assay (**Figure 1**) and examined whether knocking down or overexpressing Syd1 in DA1-ORNs would modify the Ten-m overexpression phenotypes. Compared with Ten-m overexpression alone (**Figure 3F**), co-expressing *Syd1-RNAi* to knock down Syd1 in DA1-ORNs enhanced the mismatching phenotypes (**Figure 3H**) while knocking down Syd1 alone did not affect the match index (**Figure 3E, G**; quantified in **Figure 3I**). Conversely, co-expressing wild-type Syd1 in DA1-ORNs partially suppressed the mismatching phenotype of Ten-m overexpression alone (**Figure 3K**), while overexpressing Syd1 alone did not affect the match index (**Figure 3J**; quantified in **Figure 3N**). We note that overexpressing Syd1, either alone or co-expressed with Ten-m, expanded the volume occupied by DA1-ORN axons. This phenotype may result from Syd1’s role in promoting presynaptic terminal development^54,55^. We also tested the effect of overexpressing Syd1 with a point mutation (R979A) in the RhoGAP domain that abolishes its RhoGAP activity^55^. Overexpression of Syd1-R979A also caused the expansion of DA1-ORN axons (**Figure 3L, M**), suggesting that this activity does not depend on RhoGAP activity, consistent with a previous study^55^. However, the suppression of ORN–PN mismatch was significantly reduced compared to expressing wild-type Syd1 (**Figure 3M, N**), suggesting that the regulation of ORN–PN synaptic partner matching by Syd1 is partially dependent on its RhoGAP activity.

In summary, our data indicate that Syd1 physically and genetically interacts with Ten-m. The genetic experiments further suggest a negative interaction between Ten-m and Syd1 in target selection: increasing Syd1 levels decreases Ten-m signaling, whereas decreasing Syd1 levels increases Ten-m signaling.

### Ten-m genetically interacts with Rac1 GTPase

Given RhoGTPase-related pathways highlighted by the Ten-m intracellular interactome (**Figure S3C**) and the reported RhoGAP activity of Syd1 towards Cdc42 and Rac1^55^, we next examined genetic interactions between Ten-m and Rho1, Cdc42, and Rac1 using the same Ten-m overexpression assay as above. We found that knocking down Rho1 or Cdc42 did not significantly affect the Ten-m overexpression phenotype (**Figure S4A–E**). However, knocking down Rac1 in DA1-ORNs suppressed the Ten-m overexpression phenotype (**Figure 3P**; quantified in **Figure 3S**). Conversely, overexpressing Rac1 in DA1-ORNs enhanced the Ten-m overexpression phenotype (**Figure 3R, S**). Knocking down or overexpressing Rac1 alone did not cause significant changes to the match index (**Figure 3O, Q**, **S**).

Thus, Rac1 exhibited a positive genetic interaction with Ten-m. This is consistent with the negative genetic interaction between Syd1 and Ten-m, since as a RhoGAP, Syd1 should negatively regulate Rac1 activity. Given that RhoGAP and Rho GTPases are known to mediate signaling between cell-surface receptors and the cytoskeleton (**Figure 3C**), our data suggest a signaling pathway in which Ten-m negatively regulates Syd1, and in turn activates Rac1 GTPase, in ORN axons for synaptic partner matching (**Figure 4R**).

**Figure 4.**
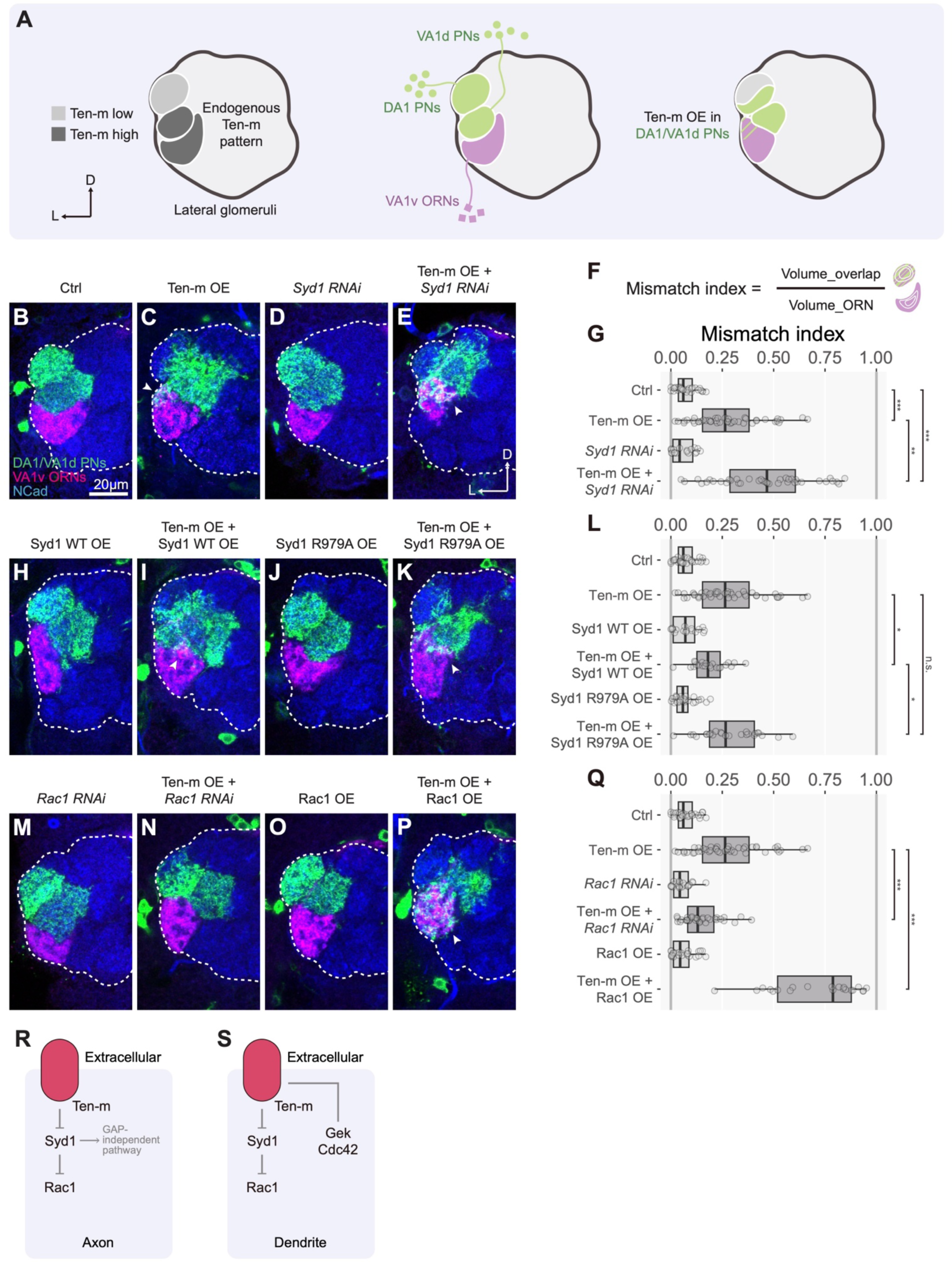
Ten-m interacts with Syd1 and Rac1 in PNs. (A) Schematic of the genetic interaction assay for Ten-m signaling in postsynaptic PN dendrites. Left, Ten-m is expressed at a low level in the DA1 glomerulus but a high level in the VA1d and VA1v glomeruli. Middle, *Mz19-GAL4* is expressed in VA1d-PNs and DA1-PNs (green), whose dendrites do not overlap with VA1v-ORN axons (purple) in control (confocal image in B). Right, Ten-m overexpression in Mz19-PNs causes a mismatch between Mz19-PN dendrites and VA1v-ORN axons (confocal image in C). (B–E) Representative confocal images of VA1v-ORN axons (magenta) and Mz19-PN dendrites (green) of control (B), Ten-m overexpression (C), *Syd1-RNAi* (D), and Ten-m overexpression with *Syd1-RNAi* (E). (F) “Mismatch index” is defined as the ratio of the overlapping volume between VA1v-ORN axons and Mz19-PNs to the volume of VA1v-ORN axons. (G) Mismatch indices for experiments in panels B–E. (H–L) Representative confocal images of VA1v-ORN axons (magenta) and Mz19-PN dendrites (green) of Syd1 overexpression (H), Syd1 and Ten-m co-overexpression (I), Syd1-R979A GAP-domain mutation overexpression (J), and Syd1-R979A and Ten-m co-overexpression (K). Mismatch indices are quantified in (L). (M–Q) Representative confocal images of VA1v-ORN axons (magenta) and Mz19-PN dendrites (green) of *Rac1-RNAi* (M), Ten-m overexpression with *Rac1-RNAi* (N), Rac1 overexpression (O), and Ten-m and Rac1 co-overexpression (P). Mismatching indices are quantified in (Q). (R, S) Summary and working models for Ten-m signaling in ORN axons (R, from data in Figure 3) and in PN dendrites (S, from data in Figure 4 and Figure S5). In both cases, Ten-m negatively regulates Syd1, and in turn activates Rac1 GTPase. Ten-m exhibits negative genetic interactions with Gek and Cdc42 only in PN dendrites. D, dorsal; L, lateral. Dashed white circle, antennal lobe. NCad, N-cadherin, a general neuropil marker. Arrowheads indicate overlap regions. The Kruskal-Wallis test with Bonferroni post-hoc correction for multiple comparisons was used in (G), (L), and (Q).

We note that manipulating Syd1 or Rac1 levels alone did not cause significant mismatching phenotypes. These data seem to contradict a key role for Syd1 and Rac1 in regulating synaptic partner matching. A likely possibility—using Rac1 as an example—is that RNAi knockdown did not reduce the Rac1 level sufficiently to interfere with its function in promoting signals from endogenous partner recognition, but interfered with a stronger signal caused by overexpressed Ten-m. It has been reported that the function of Rac1 can partially be compensated for by two other Rac GTPases in some developmental contexts^57,58^. Such dose-sensitive genetic interactions have been effectively used in identifying and analyzing intracellular signaling mechanisms, including those that involve Rac GTPases^59–61^.

### Variations of Ten-m signaling in PN dendrites for synaptic partner matching

Given the proposed homophilic attraction between Ten-m-expressing ORN axons and PN dendrites for synaptic partner matching, we next examined Ten-m signaling mechanisms in PN dendrites. We used an analogous strategy as in ORN axons by first establishing a Ten-m overexpression assay in PNs and then examining genetic interactions with candidate signaling partners. We overexpressed Ten-m using *Mz19-GAL4*, which drives transgene expression in DA1-PNs and VA1d-PNs that normally express low and high Ten-m, respectively, along with a marker to label their dendrites (**Figure 4A**). In the same flies, we also labeled VA1v ORN axons, which did not intermingle with DA1- and VA1d-PN dendrites in the control (**Figure 4B**). However, overexpressing Ten-m in Mz19-PNs caused a partial mismatching between Mz19-PN dendrites and VA1v-ORN axons (**Figure 4C**), likely due to DA1-PNs with an elevated Ten-m level now matching with VA1v ORNs, which also express high-level Ten-m^5^. This mismatching phenotype (quantified as mismatch index in **Figure 4F, G**) provided us with a quantitative assay for studying genetic interactions in PN dendrites.

We found that co-expression of a *Syd1-RNAi* transgene in Mz19*-*PNs enhanced the mismatching phenotype (**Figure 4C, E**, quantified in **Figure 4G**). Conversely, co-expression of wild-type Syd1 suppressed the mismatching phenotypes (**Figure 4C, I**, quantified in **Figure 4L**). Expression of *Syd1-RNAi* or wild-type Syd1 alone did not cause mismatching (**Figure 4D, G, H, L**). Expression of Syd1-R979A did not affect the Ten-m overexpression phenotype (**Figure 4J–L**), suggesting that the suppressive effect of expressing Syd1 is dependent on its RhoGAP activity. Furthermore, co-expression of *Rac1*-*RNAi* suppressed the Ten-m overexpression phenotype, whereas co-expression of wild-type Rac1 enhanced the Ten-m overexpression phenotype (**Figure 4M–Q**). Together, these experiments suggest a similar signaling pathway at work in PN dendrites as in ORN axons: Ten-m negatively regulates Syd1, which in turn activates Rac1 (**Figure 4S**, left; compared to **Figure 4R**). We also uncovered differences in Ten-m signaling in PN dendrites and ORN axons. Among the Ten-m intracellular interactome (**Figure 3A**; **Table S2**) was Genghis Khan (Gek), a serine/threonine kinase previously identified as an effector of the small GTPase Cdc42^62^ (**Figure S5A**). Co-immunoprecipitation experiments supported that Ten-m and Gek were in the same complex when overexpressed in S2 cells (**Figure S5B**). This prompted us to perform genetic interaction experiments in both ORN axons and PN dendrites. In ORN axons, we did not detect a significant genetic interaction between Ten-m and Gek or Gek-associated Cdc42 (**Figure S4**). However, in PN dendrites, co-expression of *Gek-RNAi* enhanced the Ten-m overexpression phenotype (**Figure S5C–E**), whereas co-expression of wild-type Gek suppressed the Ten-m overexpression phenotype (**Figure S5F, G, J**). Overexpression of a kinase-dead Gek mutant (K129A)^62,63^ did not suppress the Ten-m overexpression phenotype (**Figure S5H–J**), suggesting that the kinase activity is required for Gek’s function in counteracting Ten-m. Finally, knocking down Cdc42 also enhanced, whereas overexpressing Cdc42 suppressed, the Ten-m overexpression phenotype in dendrites (**Figure S5K–O**). Thus, Gek and its upstream activator Cdc42 appears to negatively interact with Ten-m signaling in PN dendrites but not in ORN axons (**Figure 4S**).

### Syd1 and Rac1 levels also modify *Ten-m* loss-of-function phenotypes

So far, all our *in vivo* genetic interaction experiments were performed in the context of Ten-m overexpression. We next examined whether genetic interactions also occurred in the context of loss of endogenous Ten-m function. To achieve this, we identified a split-GAL4 combination that specifically drove expression with an early onset in VA1d-ORNs, which express high Ten-m^5^ (**Figure 5A**). We found that expressing *Ten-m-RNAi* in VA1d-ORNs caused a fraction of VA1d-ORN axons to innervate the neighboring DA1 glomerulus, which expresses a low level of Ten-m (compare **Figure 5B, D**, quantified in **Figure 5G**; mistarget index defined in **Figure 5F**). Elevating the level of *Ten-m-RNAi* expression at 29°C compared to at 25°C resulted in a stronger mistargeting phenotype (**Figure 5C, D**, **G**), suggesting that loss of Ten-m in VA1d-ORNs caused a level-dependent mistargeting of axons to the DA1 glomerulus.

**Figure 5.**
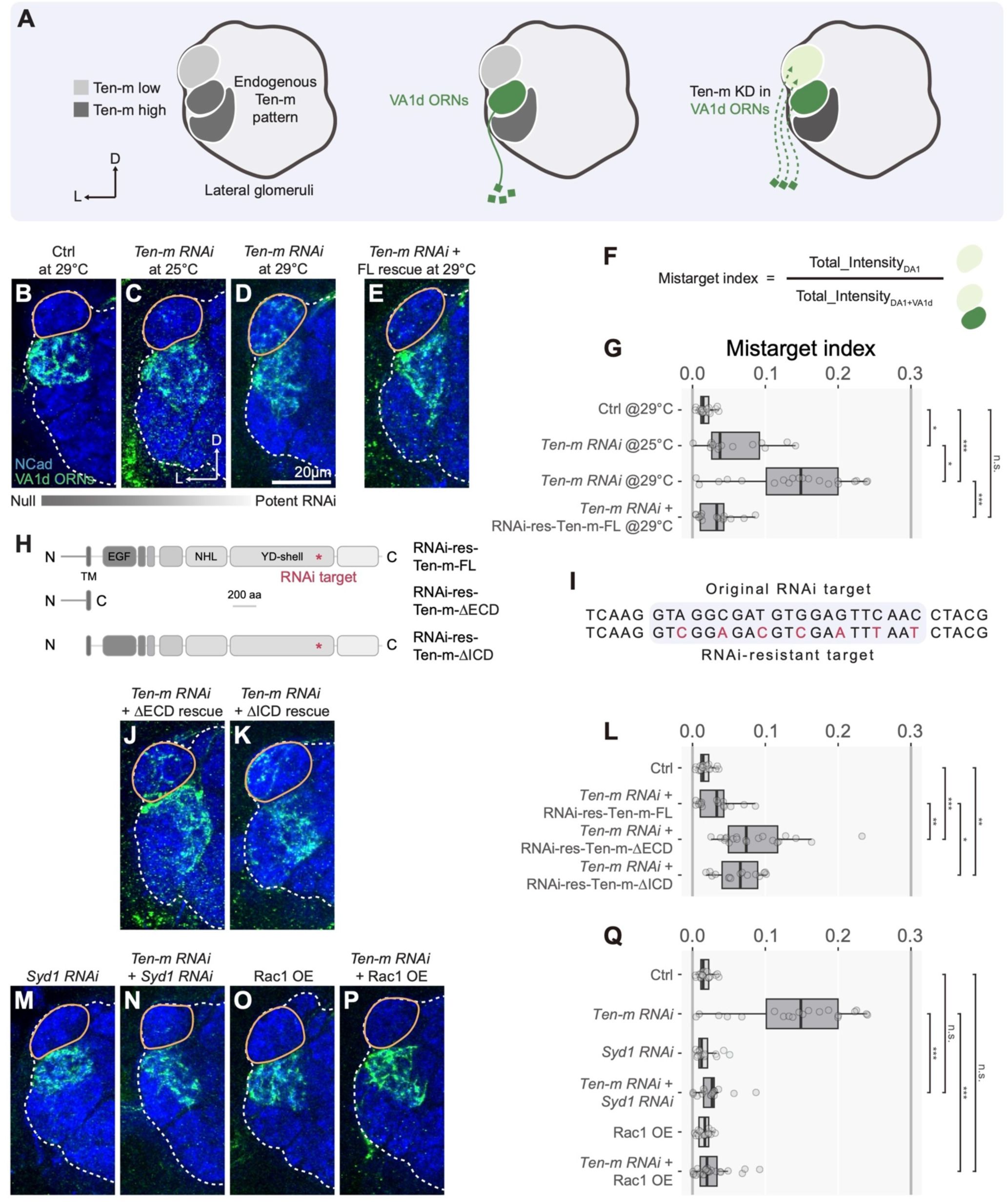
Syd1 and Rac1 modify *Ten-m* loss-of-function phenotypes. (A) Schematic of investigating the Ten-m signaling using loss-of-function assays. Left, Ten-m is expressed at a low level in the DA1 glomerulus but a high level in the VA1d and VA1v glomeruli. Middle, a split GAL4 pair specifically labels VA1d-ORNs (confocal image in B). Right, knocking down Ten-m in VA1d-ORNs causes partial mistargeting of VA1d-ORNs to the DA1 glomerulus and mismatching with DA1-PN dendrites (confocal images in C). (B–E) Representative confocal images of VA1d-ORN axons of control (B), *Ten-m-RNAi* at 25°C (C), *Ten-m-RNAi* at 29°C (D), and *Ten-m-RNAi* with RNAi-resistant full-length (FL) *Ten-m* rescue at 29°C (E). (F) “Mistarget index” is defined as the ratio of total GFP fluorescence intensity of VA1d-ORN axons in the DA1 glomerulus to that in the DA1 and VA1d glomeruli. (G) Mistarget indices of experiments in panels B–E. (H) Location of the RNAi target in *Ten-m*. (I) Comparison of the original and RNAi-resistant *Ten-m* transgene sequences at the RNAi target site. (J–L) Representative confocal images of VA1d-ORN axons of *Ten-m-RNAi* with RNAi-resistant Ten-m-ΔECD rescue (J) or RNAi-resistant Ten-m-ΔICD rescue (K) at 29°C. Mistarget indices are quantified in (L). (M–Q) Representative confocal images of VA1d-ORN axons of *Syd1-RNAi* (M), *Ten-m-RNAi* and *Syd1-RNAi* (N), Rac1 overexpression (O), and *Ten-m-RNAi* with Rac1 overexpression (P). Mistarget indices are quantified in (Q). D, dorsal; L, lateral. Dashed white circle, antennal lobe. Yellow circle, DA1 glomerulus. Ncad, N-cadherin, a general neuropil marker. The Kruskal-Wallis test with Bonferroni post-hoc correction for multiple comparisons was used in (G), (L), and (Q).

To validate if the mistargeting was due to knocking down Ten-m, we designed an RNAi-resistant transgene that encodes the full-length Ten-m protein (**Figure 5H, I**). Co-expression of this transgene rescued the mistargeting phenotype due to *Ten-m-RNAi* expression (**Figure 5E**, **G**). However, co-expression of an RNAi-resistant Ten-m without the extracellular or intracellular domains was not as effective in rescuing (**Figure 5J, K**, quantified in **Figure 5L**), suggesting that both the extracellular and intracellular domains contribute to preventing VA1d-ORN axons from mismatching with DA1-PN dendrites.

Next, we used this loss-of-function assays to test for genetic interactions between Ten-m and Syd1 or Rac1. While expressing *Syd1-RNAi* or wild-type Rac1 alone did not cause significant mismatching (**Figure 5M, O**, quantified in **Figure 5Q**), co-expression of *Syd1-RNAi* or wild-type Rac1 with *Ten-m-RNAi* suppressed the mismatching between VA1d-ORN axons and DA1-PN dendrites (**Figure 5N, P**, **Q**). These experiments support the signaling pathway deduced from our gain-of-function genetic assay: that Ten-m negatively regulates Syd1, and in turn activates the Rac1 GTPase (**Figure 4R**).

### Single-axon analyses reveal that synaptic partner matching results from selective stabilization of ORN axon branches that contact partner PN dendrites

To examine in detail how Ten-m signaling affects the behavior of ORN axons during each step of wiring specificity establishment, we next developed a sparse driver system to limit the labeling and genetic manipulation to a fraction of neurons of a particular type (**Figure 6A, S6A, S6B**). We inserted a transcriptional stop flanked by two mutant *FRT* (*FRT10*) sites with a reduced FLP-mediated recombination rate^64^ in between the start codon and coding sequence of a driver transgene. The probability that the sparse driver is expressed in a particular cell type can be controlled by FLP expression level or duration. Since our DA1-ORN driver utilized the intersection of two transgenes expressing a DNA-binding domain (DBD) and a transcription activation domain (AD), respectively, we inserted the *FRT10-stop-FRT10* cassette in the transgene that expressed the activation domain. This sparse strategy can be used to co-express multiple transgenes simultaneously for visualizing and genetically manipulating a subset of neurons of a given cell type (**Figure 6A**). By varying the heat shock duration, we could label a large subset, an intermediate subset, or a single DA1-ORN axon (**Figure 6B**, **S6B–D**).

**Figure 6.**
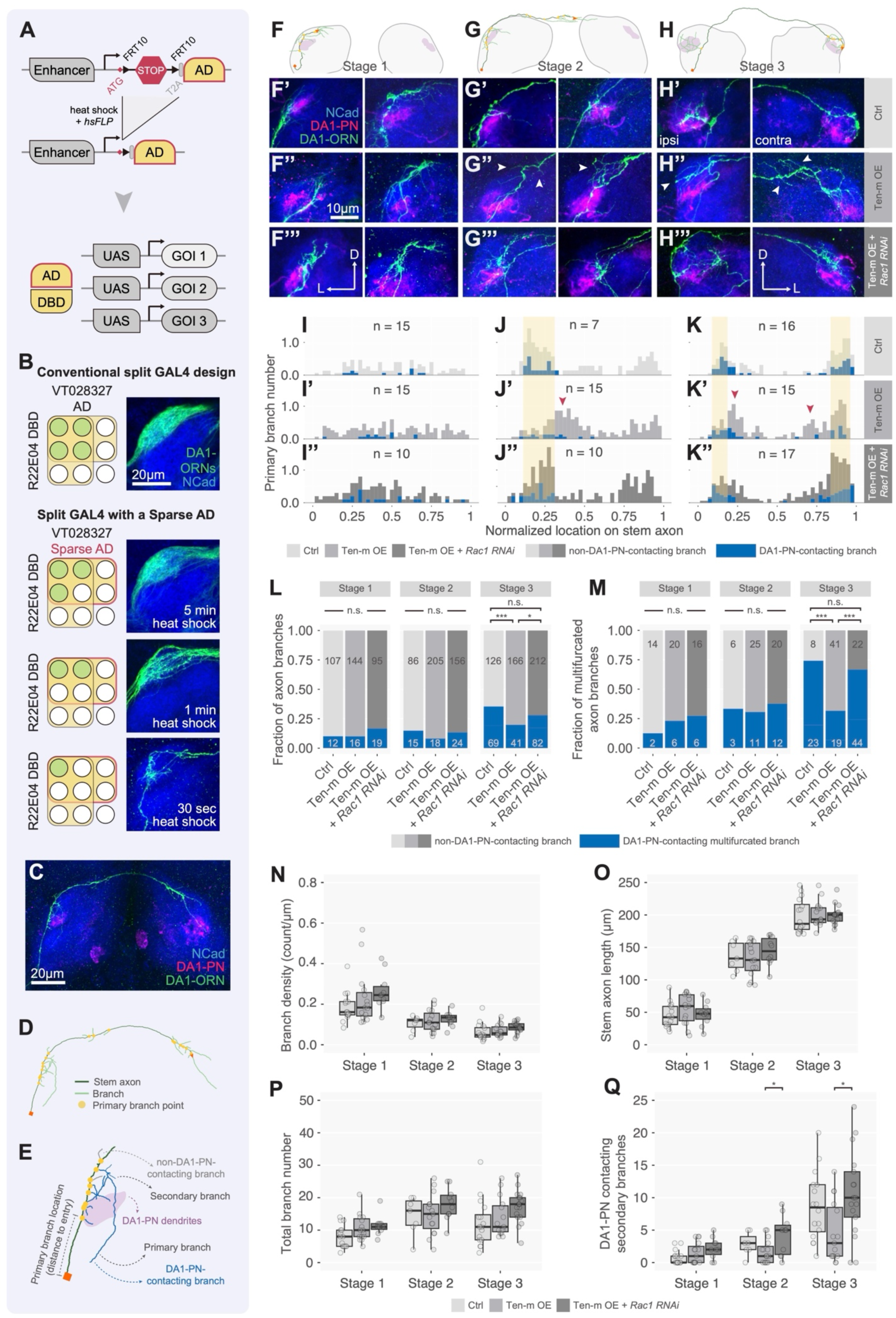
Analysis of Ten-m signaling with single-axon resolution. (A) Schematic of the “sparse driver” strategy. In a split GAL4 pair, the transcription activation domain (AD, from the transcription factor p65) is under the control of an enhancer separated by *FRT10-STOP-FRT10.* FLP-induced recombination between *FRT10* sites occurs at ∼10% efficiency compared to wild-type *FRT* sites. STOP designates a transcription termination sequence. Heat shock-induced FLP expression from the *hsFLP* transgene causes flip-out of the STOP and thus expression of AD in a fraction of cells, which together with the GAL4 DNA-binding domain (DBD) expressed from a separate transgene would reconstitute functional GAL4, driving co-expression of multiple genes of interest (GOI) in these cells. (B) Compared to the conventional split GAL4 design, sparse driver enables different sparsity of transgene expression tuned by heat shock time. (C) Example of a single developing DA1-ORN axon crossing the commissure, enabled by the sparse driver. NCad, N-cadherin, a general neuropil marker. (D) Z-projection of the 3D trace of the example DA1-ORN axon shown in (C) illustrating quantitative parameters extracted from the trace. Length of the stem axon (dark green) is measured from the antennal lobe entry point (orange square) to the end point (orange triangle). A primary branch point (yellow dot) is where a collateral branch (light green) intersects with the stem axon. (E) Zoom-in to the antennal lobe entry segment of the example DA1-ORN axon. Primary branch location is defined as the distance between the antennal lobe entry point (orange square) and the primary branch point (yellow dot). Some primary and secondary DA1-ORN branches are in contact with DA1-PN dendrites (purple shade). (F–H) Schematic of the three stages of a developing DA1-ORN axon. (F) Stage 1; stem axon length below 100 µm, usually before the axon crosses the commissure. (G) Stage 2; stem axon length from 100 to 170 µm; most axons have crossed the commissure but have not reached the contralateral PN dendrites. (H) Stage 3; stem axon length >170 µm, when the axon reaches with the contralateral PN dendrites. Purple shade, DA1-PN dendrites. (F’-H’’’) Representative maximum Z-projection images of sparse DA1-ORN axons in control (F’–H’), Ten-m overexpression (F’’–H’’), and Ten-m overexpression with *Rac1-RNAi* (F’’’–H’’’) at each developmental stage Two examples per genotype are shown for Stages 1 and 2. For Stage 3, a single example in both ipsilateral (left) and contralateral (right) antennal lobes is shown. NCad, N-cadherin, a general neuropil marker. Arrowheads indicate dorsomedial shifted branches. (I–K’’) Histograms of primary branch location distribution of DA1-ORN axons in control (I–K, top row), Ten-m overexpression (I’–K’, middle row), and Ten-m overexpression with *Rac1-RNAi* (I’’–K’’, bottom row) at each developmental stage (Stage 1, left column, I–I’’; Stage 2, middle column, J–J’’; Stage 3, right column, K–K’’). The location (x-axis) is normalized to the stem axon length. On the x-axis, 0 represents the antennal lobe entry point and 1 represents the end point of the stem axon. As axons align along the x-axis, right shifts of ipsilateral branches and left shifts of contralateral branches indicate dorsomedial shifting. Blue portions of the histogram indicate DA1-ORN axon branches that are in contact with DA1-PN dendrites. Yellow shade indicates peaks of DA1-PN contacting branches of the control group. Red arrowheads indicate shifted histogram peaks revealing ectopically targeted axons. (L) Fractions of DA1-ORN axon branches in contact with DA1-PN dendrites at each developmental stage in each genotype: control, Ten-m overexpression, and Ten-m overexpression with *Rac1-RNAi*. Blue and gray represent DA1-PN-contacting and non-DA1-PN-contacting branches, respectively. (M) Fractions of multifurcated DA1-ORN axon branches in contact with DA1-PN dendrites at each developmental stage in each genotype: control, Ten-m overexpression, and Ten-m overexpression with *Rac1-RNAi*. Blue and gray represent DA1-PN-contacting and non-DA1-PN-contacting branches, respectively. A primary axon branch with at least one secondary branch is categorized as multifurcated. (N–Q) Quantification of branch densities (total branch number normalized to the stem axon length) (N), stem axon lengths (O), total branch number (P) and DA1-PN-contacting secondary branch number (Q) at each developmental stage for the listed genotypes. The chi-squared tests (L and M) and the one-way ANOVA (with Tukey’s test) (N–Q) were used for multiple comparisons.

Our recent live-imaging experiments in the antenna-brain explant suggested that an individual ORN axon extends multiple ipsilateral branches along the main axon trunk (which we refer to as the stem axon hereafter), and a subset of branches are subsequently stabilized^64^. These observations were made from ORNs whose glomerular types were determined *post hoc*, and thus we had too few examples of any genetically defined ORN type to provide quantitative assessments. Furthermore, we did not label the postsynaptic target of the observed ORN type to assess what subset of branches were selectively stabilized. The sparse driver strategy for DA1-ORNs, concomitant with the labeling of DA1-PN dendrites in a different color, allowed us to systematically characterize the behavior of individual ORN axons during targeting selection by focusing on brains in which a single DA1-ORN axon was labeled (**Figure 6C–E**; **Figure S6**).

We sorted our samples into three stages of development based on the length of the stem axon. Stage 1 covered samples in which DA1-ORN axons were extending within the ipsilateral antennal lobe (**Figure 6F**). Stage 2 included samples in which DA1-ORN axons extended near the midline, across the midline, and in the contralateral antennal lobe (**Figure 6G**). By Stage 3, stem axons had reached the contralateral DA1 targets and elaborated branches in contact with contralateral DA1-PN dendrites (**Figure 6H**).

We first analyzed the distribution of primary branch points along the stem axon in control animals, normalized to the total length of the stem axon. At Stage 1, branching points were widely distributed along the entire ipsilateral stem axon (**Figure 6F’, 6I**); only a small fraction of these branches contacted DA1-PN dendrites (**Figure 6F’**; blue in **Figure 6I**, **L**). At Stage 2, while the total primary branch density decreased compared to Stage 1 (**Figure 6N**), more branches were in contact with DA1-PN dendrites (**Figure 6G’, J, L**). At Stage 3, the primary branches continued to cluster near the DA1-PN dendrites (**Figure 6H’, K**), the primary branch density in the ipsilateral antennal lobe further decreased (**Figure 6H’**, left; **Figure 6N**), and the fraction of DA1-ORN branches in contact with DA1-PN dendrites further increased (**Figure 6K, L**). Moreover, DA1-ORN axons produced many branches in the contralateral antennal lobe, some of which contacted the contralateral DA1-PN dendrites (**Figure 6H’**, right; **Figure 6K**).

We also quantified the total number of multifurcated branches (primary branches that branch further) as well as multifurcated branches in contact with DA1-PN dendrites. We found that the number of multifurcated branches and particularly those contacting DA1-PN dendrites increased substantially at Stage 3 (**Figure 6M**, left columns).

In summary, quantitative single-axon analyses revealed that, in wild type, DA1-ORN axons send many primary branches as the stem axon extends along the surface of the antennal lobe towards the midline. Branch points occur along almost the entire stem axon. As development proceeds, branch density decreases, branch points become more concentrated near DA1-PN dendrites, more branches contact PN dendrites, and more high-order branches emerge from DA1-PN dendrite contacting primary branches. These observations support a model in which stabilization of ORN axon branches by target PN dendrites is a key mechanism by which target selection is achieved (**Figure 7G, H**).

**Figure 7.**
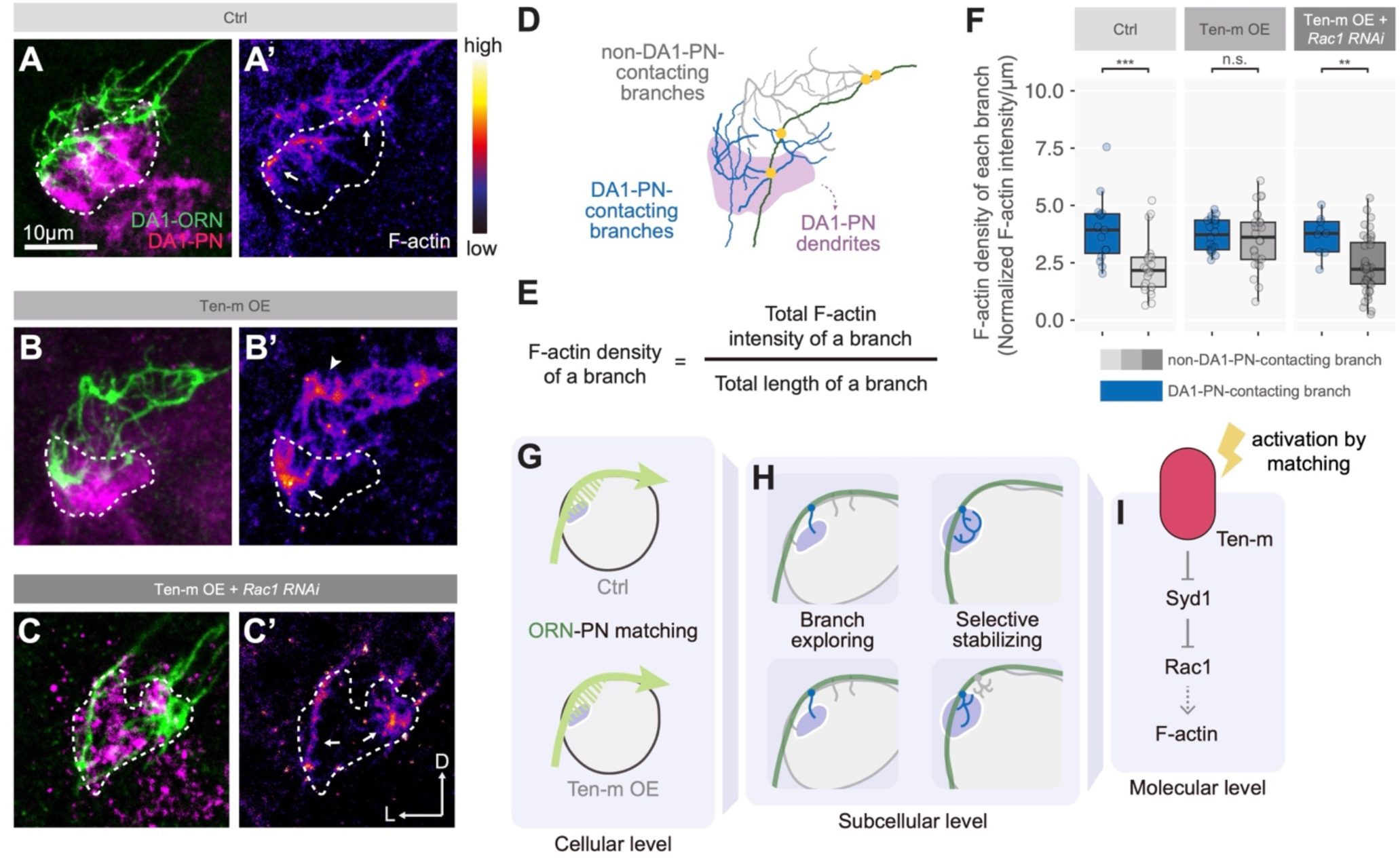
F-actin distribution analysis and summary. (A–C’) Representative confocal images of DA1-PN dendrites (A–C, magenta), DA1-ORN axons (A–C, green), and F-actin distribution in the same DA1-ORN axons (A’–C’, heatmap gradient based on Halo-Moesin staining) of control (A and A’), Ten-m overexpression (B and B’), and Ten-m overexpression with Rac1 knockdown (C and C’). Arrows indicate F-actin hot spots in DA1-contacting branches; arrowheads indicate F-actin hot spots in non-DA1-contacting branches. D, dorsal; L, lateral. Dashed white circle, DA1-PN dendrites. (D) Classification of DA1-ORN axon branches for quantifying axonal F-actin distribution: blue, axon branches contacting DA1-PN dendrites; gray, axon branches that do not contact DA1-PN dendrites; dark green, the stem axon; and purple shade, DA1-PN dendrites. (E) F-actin density of an axon branch is defined as the ratio of the total F-actin (Halo-Moesin staining) intensity of the branch to the branch length. (F) F-actin density of each axon branch of control (left), Ten-m overexpression (middle), and Ten-m overexpression with *Rac1-RNAi* (right). Each dot represents one DA1-ORN axon branch that contacts (blue) or does not contact (gray) DA1-PN dendrites. 15 DA1-PN-contacting and 22 non-DA1-PN-contacting branches from 2 axons for control; 20 DA1-PN-contacting and 26 non-DA1-PN-contacting branches from 4 axons for the Ten-m-overexpression group; 9 DA1-PN-contacting and 39 non-DA1-PN-contacting branches from 3 axons for the Ten-m-overexpression and *Rac1-RNAi* group. The Mann-Whitney *U* test was used for comparisons. (G–I) Summary of the Ten-m signaling in synaptic partner matching. Ten-m level directs ORN–PN synaptic partner matching (G; see also Figure 1 and our previous study^5^). Temporally resolved single-axon analysis revealed that Ten-m specifically acts at the step of stabilizing axon branches but not general axon growth or branch exploration (H; see also Figure 6). *In situ* proximity labeling proteomics (Figure 2) and *in vivo* genetic perturbations (Figures 3–7) delineated the signaling axis: Ten-m negatively regulates the RhoGAP Syd1 and, in turn, activates the Rac1 GTPase to tune F-actin distribution (I).

### Ten-m signaling promotes stabilization of ORN axon branches that contact partner PN dendrites

We next probed the cellular mechanism by which perturbing Ten-m signaling affects synaptic partner matching using single-axon analysis of DA1-ORNs. We focused on two genotypes in comparison with the control described above: (1) Ten-m overexpression in DA1-ORNs, which caused mismatching between DA1-ORNs and DA1-PNs when assayed in bulk (**Figure 1F–I**), and (2) Ten-m overexpression together with RNAi against *Rac1* in DA1-ORNs, which ameliorated the mismatching phenotype caused by Ten-m overexpression (**Figure 3P**, **S**).

Compared to controls, neither of the experimental conditions significantly affected branch density (**Figure 6N**), stem axon length (**Figure 6O**), or total branch number (**Figure 6P**) at all three stages. Perturbing Ten-m signaling also did not affect the distribution of axon branches along the stem axon (**Figure 6F’’**, **6I’**) or fractions of axon branches contacting DA1-PN dendrites (**Figure 6F’’**, **6L, 6M**) at early developmental stages. However, beginning at Stage 2 (**Figure 6G’’**, **6J’**) and continuing at Stage 3 (**Figure 6H’’**, **6K’**), axon branches of Ten-m-overexpressing DA1-ORNs were further from the origin of stem axons compared to the control, consistent with the mistargeting of axons in the bulk ORN assay (**Figure 1F–H**, **Figure S2F–I**). Reducing Rac1 levels in Ten-m-overexpressing DA1-ORN neurons shifted the branch distribution back to the control pattern (**Figure 6G’’’**, **6H’’’**, **6J’’**, **6K’’**). As a result, Rac1 knockdown suppressed the reduction of DA1-PN-contacting ORN axon branches when Ten-m was overexpressed (**Figure 6L**), most strikingly for the multifurcated axons at Stage 3 (**Figure 6M**, **6Q**).

These data indicate that perturbing Ten-m signaling alters neither general axon growth and branching, nor initial stages of branch exploration. Rather, Ten-m signaling promotes stabilization of ORN axon branches that contact dendrites of their postsynaptic partner PNs, particularly for higher-order branches. That almost all phenotypes caused by Ten-m overexpression at single-axon resolution could be suppressed by reducing Rac1 level reinforces the notion that Rac1 is a key mediator of Ten-m signaling in synaptic partner selection (**Figure 7H, 7I**).

### Partner recognition promotes actin polymerization in axon branches

Given that a key function of Rac1 signaling is cytoskeletal regulation^56,65–68^, we next examined the distribution of microtubules and polymerized (filamentous) actin (F-actin) using transgenic markers expressed from sparsely labeled ORNs. We found that microtubule markers—a tagged tubulin subunit^69^ or EB1 (which labels growing microtubule plus ends^70^)—were distributed along the entire length of DA1-ORN axons (**Figure S7A–B’**). By contrast, Halo-Moesin, which binds preferentially to F-actin^71^, was preferentially localized to subcellular regions close to the DA1 glomerulus (**Figure S7C, C’**), suggesting a role for F-actin in synaptic partner matching.

Focusing on F-actin distribution, we next examined samples from control, Ten-m overexpression, and Ten-m overexpression with Rac1 knockdown groups in which only 1–3 axons were labeled, such that we could resolve individual branches (**Figure 7A–C**). We also co-labeled DA1-PN dendrites, such that we could determine whether an individual ORN axon branch contacted DA1-PN dendrites (**Figure 7D**). In the control, DA1-ORN axon branches contacting DA1-PN dendrites had significantly higher F-actin density than those not contacting DA1-PN dendrites (**Figure 7E**; **Figure 7F**, left). This suggests that ORN axons receive a signal from the cognate PN dendrites, which promotes actin polymerization in ORN axons that might support the initiation of synaptic connections. In Ten-m-overexpressing DA1-ORNs, this difference disappeared (**Figure 7F**, middle), likely because ORN branches that did not contact DA1-PN dendrites could receive a partner matching signal from non-cognate PN partners such as DL3 (**Figure S2**), which activated Ten-m and Rac1 and thus promoted actin polymerization. In DA1-ORNs with Ten-m overexpression and Rac1 knockdown, non-DA1-PN-contacting branches had a significant reduction in F-actin compared to DA1-PN-contacting branches again (**Figure 7F**, right), consistent with the observation that a reduced level of Rac1 diminished the effect of Ten-m signaling and further suggesting that Rac1 is the key mediator transforming Ten-m signaling into actin network regulation (**Figure 7I**).

## DISCUSSION

By manipulating the levels of Ten-m, a synaptic partner matching regulator^5,6^ and by co-labeling single axons of a defined neuron type together with dendrites of their postsynaptic partners in the fly olfactory system across multiple developmental stages, we showed that synaptic partner matching is primarily mediated by selective stabilization of axon branches that contact dendrites of postsynaptic partners (**Figure 1**, **Figure 6**). Combining *in situ* proximity labeling, proteomic analysis, and *in vivo* genetic interaction studies, we further elucidated molecular pathways by which Ten-m signals to the actin cytoskeleton to mediate its function in synaptic partner matching (**Figures 2–7**).

### Cellular mechanisms of synaptic partner matching

An essential final step of establishing wiring specificity is to select synaptic partners among many non-partner cells. Studies of the neuromuscular junctions from insects to mammals, where motor axons and their distinct muscle targets can be readily resolved by light microscopy, suggest that axons of specific motor neuron types (or motor pools in vertebrates) navigate precisely to their specific muscle targets^72–75^. By contrast, connectivity between individual motor neurons *within the same motor pool* and specific muscle fibers *within the same muscle* appears more stochastic, involving the formation of exuberant connections followed by extensive synapse elimination in an activity-dependent manner^73,76–78^. Cellular mechanisms by which synaptic partner selection is achieved in the central nervous system is more difficult to discern because this involves visualizing pre- and postsynaptic partners with synaptic resolution or performing electrophysical recordings. Two of the best studied systems, the climbing fiber–Purkinje cell connections and eye-specific connections between retinal ganglion cells and lateral genicular nucleus target neurons, both involve initially forming exuberant connections followed by activity-dependent synapse elimination^79^.

The glomerular organization of the olfactory systems from insects to mammals provides an ideal model to investigate mechanisms of synaptic partner matching in the central nervous system. The convergence of axons of the same ORN type and dendrites of their cognate postsynaptic partner PNs (equivalent to mitral/tufted cells in vertebrates) to discrete glomeruli means that synaptic partner matching can be examined by light microscopy, since glomerular targeting is equivalent to synaptic partner matching. Indeed, in *Drosophila*, serial electron microscopic studies indicate that all ORN axons that target to a specific glomerulus form synaptic connections with all partner PNs^80–83^. By developing genetic drivers that allowed us to track individual types of ORNs and their partner PNs, and by sparsely labeling ORN axons of a specific type, we could chart developmental time course of synaptic partner matching with single-axon resolution. Our data indicate that after an ORN axon chooses a specific trajectory^84,85^, it produces exuberant branches followed by stabilization of those that contact dendrites of the postsynaptic partner (**Figure 6**). Misexpressing Ten-m, an instructive synaptic partner matching molecule and thereby partially respecifying its synaptic partners, causes corresponding stabilized axonal branches at a new target (**Figure 6**). Collectively, these data suggest that synaptic partner matching is largely achieved by selective axon branch stabilization resulting from molecular signaling between synaptic partners.

Our finding superficially resembles the formation of exuberant connections followed by synapse elimination in the vertebrate neuromuscular junction, climbing fiber–Purkinje cell connections, and retinal ganglion cell–lateral geniculate connections discussed above, as well as activity-dependent refinement of ORN axons and mitral cell dendrites in glomeruli of the mammalian olfactory bulb^86,87^. A fundamental difference is the time scale. While the exuberant connections in the vertebrate systems last on the order of days and involve synapse formation and elimination, the exuberant ORN axon branches we observed lasted on the order of hours to minutes (**Figure 6**; see also ^64^). Furthermore, the developmental timing of ORN axon target selection precedes synaptogenesis in the *Drosophila* brain^88^ or onset of odorant receptor expression^89^, making it unlikely to be dependent on synaptic or sensory activity. Thus, we propose that the exuberant ORN axon branches serve the purpose of expanding the search space for molecular interactions between ORN axons and their synaptic partners (resulting in stabilization) or non-partners (resulting in pruning). It will be interesting to see whether a similar mechanism operates in synaptic partner matching in other circuits in the fly and vertebrate nervous systems.

### Intracellular signaling mechanisms of Ten-m in synaptic partner matching

Most well-studied receptors for intercellular signaling are type-I single-pass transmembrane proteins or G-protein-coupled receptors that span the membrane 7 times^37,90–94^. Little is known how intracellular signaling works for type-II single-pass transmembrane proteins like teneurins. Intracellular domains of teneurins do not have motifs suggestive of engaging specific signaling pathways. Thus, to understand teneurin intracellular signaling, we took an unbiased approach of identifying potential interaction partners using proximity labeling followed by quantitative mass-spectrometry analysis, which captures both stable and transient molecular partners *in situ* in developing fly brains with a proteome-wide coverage^43^. Ten-m interactome included broad classes of proteins localized at the cell surface, synapse, cytoplasm, and endomembrane systems (**Figure 2, Figure S3**). Future investigation into proteins with different cellular component classifications could deepen our understanding of type II membrane proteins and answer whether their inverted topology (N-terminal intracellular domain) engages distinct pathways for protein trafficking, post-translational modification, quality control, and proteolysis.

By establishing quantitative phenotypic assays for genetic interactions *in vivo*, we identified a key signaling pathway that links Ten-m to the actin cytoskeleton in synaptic partner matching, through regulation of a RhoGAP, Syd1, and the Rac1 small GTPase (**Figure 7I**). This pathway is supported by genetic interaction data for both RhoGAP and Rac1, using both overexpression and knockdown manipulations of RhoGAP and Rac1, in Ten-m gain-of-function and loss-of-function contexts, and in bulk as well as in single-axon assays. Rho GTPases are key regulators of the actin cytoskeleton and have been implicated as mediators of growth cone signaling downstream of multiple classic guidance receptors, vast majority of which are type-I transmembrane proteins^49,56,67,68,90,95,96^. That Rac1 is also a key mediator of signaling downstream of a type-II transmembrane protein, Ten-m, highlights the importance of Rho GTPases as a signaling hub.

### Variations of teneurin signaling in diverse biological processes

Given that teneurins play diverse roles in many biological processes (see **Introduction**), to what extent does the signaling pathway we identified in synaptic partner matching apply to other processes? While this is to our knowledge the first study on intracellular signaling of teneurins and we therefore lack comparison data, the comparison of synaptic partner matching in presynaptic ORN axons and postsynaptic PN dendrites (**Figure 4R, 4S**) is instructive. Even though Ten-m mediates homophilic attraction between cognate ORN axons and PN dendrites, chemical synapses are inherently asymmetrical^97^. Furthermore, in the wiring of the olfactory circuit, PN dendrites initiate patterning followed by ORN axon targeting, thus the two processes have distinct spatiotemporal constraints. Despite these differences, signaling in PNs and ORNs share the same core motif: Ten-m negatively regulates Syd1, which in turn activates Rac1. However, the Cdc42/Gek pathway, which negatively regulates Ten-m signaling, is preferentially used in PNs, suggesting that cell-type-or cellular-compartment-specific variations of signaling do occur.

Since many of the teneurin-mediated biological processes, including cell polarity, neuronal migration, axon guidance, and synaptic organization, involve extracellular interaction–modulated cell shape changes, we speculate that the pathway we identified involving Rho GTPase signaling to the actin cytoskeleton, or some variations on the same theme, will likely be utilized. In addition to Syd1 and Gek, other members of the intracellular interactomes that we have not explored functionally (**Figure 3A**; **Table S2**) could provide entry points for investigating additional signaling pathways.

### High-resolution methods for developmental analysis *in vivo*

In neural circuit wiring and other developmental processes, molecular signaling directs cellular behaviors. However, *in vivo* genetic analysis to interrogate functions of specific molecules and in-depth cell biological studies are often detached from each other due to separate experimental paradigms. Our study attempts to break this barrier by developing or utilizing a variety of high-resolution methods, from spatial proteomics to sparse driver–based single-axon analysis. Specifically, *in situ* proximity labeling with high spatiotemporal resolution and quantitative mass spectrometry, such as what we employed here (**Figure 2**), could enable one to identify proteome-wide interacting partners of key proteins involved in any biological process at the desired developmental time and physiological subcellular location. This can then inform high-resolution phenotypic analysis and genetic interaction studies to validate the *in vivo* relevance of interacting partners.

In the context of neural circuit development, sparse neuronal labeling and genetic manipulation enable the morphology and connectivity analysis of individual neurons within dense networks of the central nervous system and the study of cell-autonomous gene function^98–103^. However, with regard to genetic manipulation, methods relying on probabilistic gating of transgene expression often fail to co-express all desired genes of interest in the same sparsely labeled neurons because different effector or reporter transgenes may be stochastically expressed in independent subsets of neurons^64,104,105^. Probabilistic expression of a driver transgene, which controls the expression of multiple effector or reporter transgenes, should in principle overcome this caveat. The MARCM system^106^ is such an example, but because it relies on the loss of a repressor after mitotic recombination, perdurance of the repressor from parental cells may limit the effectiveness of analyzing developmental events shortly after mitotic recombination^107^.

Our sparse driver strategy (**Figure 6A**) achieved this goal by using a FLPout strategy that combines mutant *FRT* sites with reduced recombination efficiency and tunable FLP recombinase levels. Sparse expression of a split activation domain further enabled the combinatorial use with a variety of existing transgenes expressing the DNA-binding domains of transcription factors such as GAL4, QF2, LexA^108–110^ in specific cell types, enabling the timely co-expression of multiple transgenes in cell-type-specific sparse neurons. The sparse driver strategy allowed us to perform multi-parameter quantification of developing single axons while genetically manipulating Ten-m and Rac1 (**Figure 6E–Q**). We envision that the combination of the above strategies could be used in dissecting cellular and molecular mechanisms of other developmental processes, with the goal of integrating *in vivo* cell biology with their underlying molecular signaling cascades.

## STAR METHODS

## KEY RESOURCES TABLE

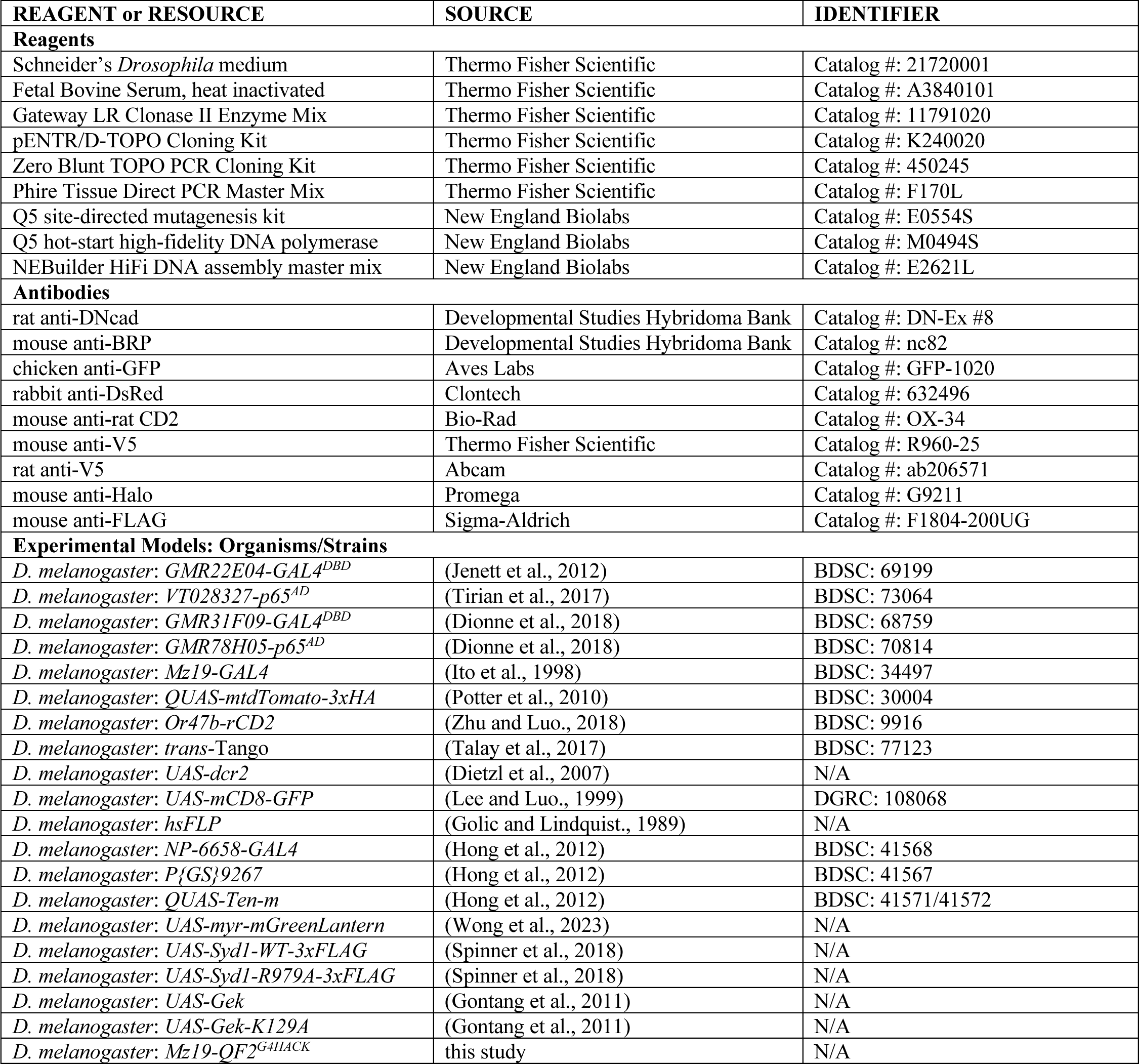
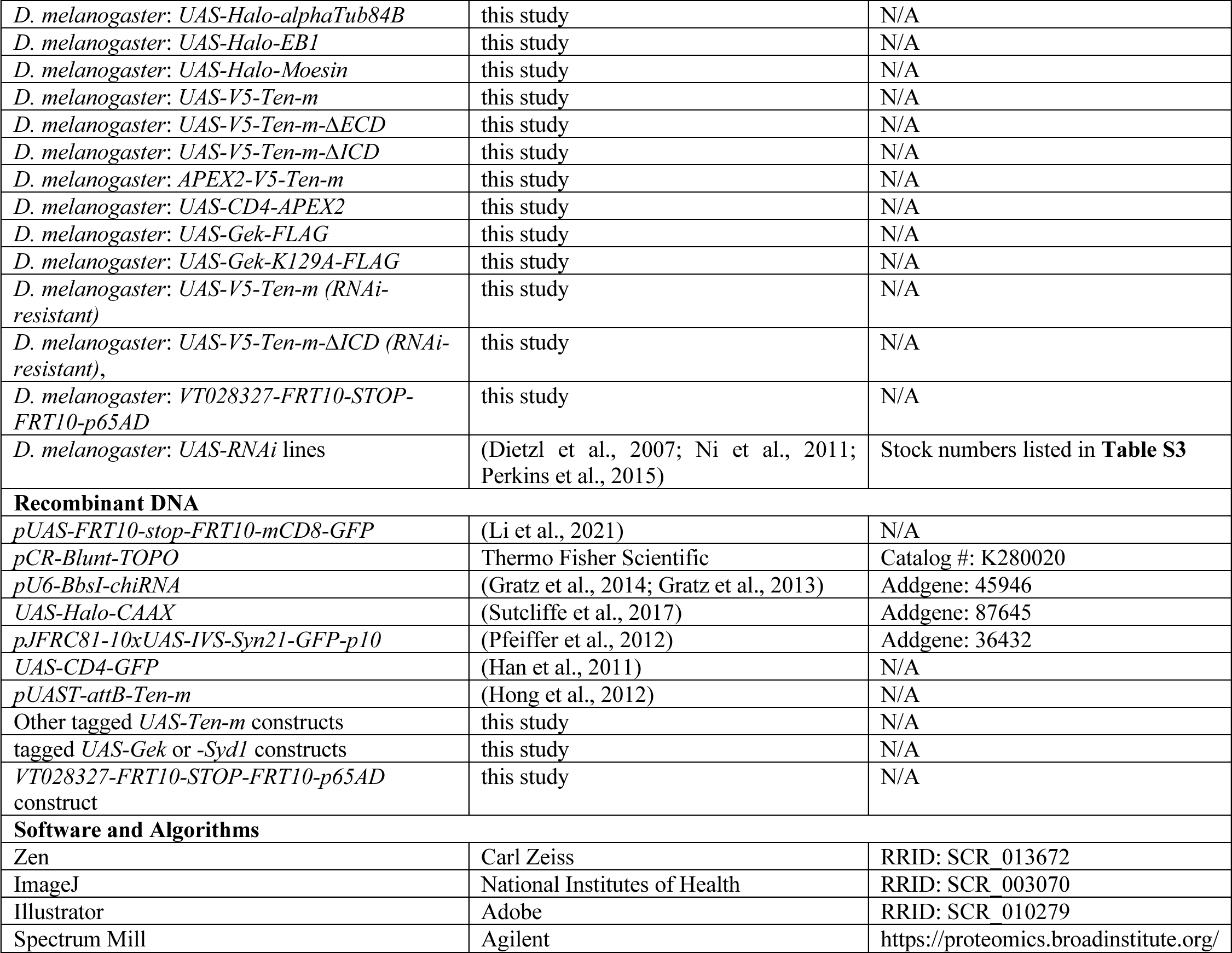

| REAGENT or RESOURCE | SOURCE | IDENTIFIER |
| --- | --- | --- |
| **Reagents** |  |  |
| Schneider's *Drosophila* medium | Thermo Fisher Scientific | Catalog #: 21720001 |
| Fetal Bovine Serum, heat inactivated | Thermo Fisher Scientific | Catalog #: A3840101 |
| Gateway LR Clonase II Enzyme Mix | Thermo Fisher Scientific | Catalog #: 11791020 |
| pENTR/D-TOPO Cloning Kit | Thermo Fisher Scientific | Catalog #: K240020 |
| Zero Blunt TOPO PCR Cloning Kit | Thermo Fisher Scientific | Catalog #: 450245 |
| Phire Tissue Direct PCR Master Mix | Thermo Fisher Scientific | Catalog #: F170L |
| Q5 site-directed mutagenesis kit | New England Biolabs | Catalog #: E0554S |
| Q5 hot-start high-fidelity DNA polymerase | New England Biolabs | Catalog #: M0494S |
| NEBuilder HiFi DNA assembly master mix | New England Biolabs | Catalog #: E2621L |
| **Antibodies** |  |  |
| rat anti-DNcad | Developmental Studies Hybridoma Bank | Catalog #: DN-Ex #8 |
| mouse anti-BRP | Developmental Studies Hybridoma Bank | Catalog #: nc82 |
| chicken anti-GFP | Aves Labs | Catalog #: GFP-1020 |
| rabbit anti-DsRed | Clontech | Catalog #: 632496 |
| mouse anti-rat CD2 | Bio-Rad | Catalog #: OX-34 |
| mouse anti-V5 | Thermo Fisher Scientific | Catalog #: R960-25 |
| rat anti-V5 | Abcam | Catalog #: ab206571 |
| mouse anti-Halo | Promega | Catalog #: G9211 |
| mouse anti-FLAG | Sigma-Aldrich | Catalog #: F1804-200UG |
| **Experimental Models: Organisms/Strains** |  |  |
| *D. melanogaster*: GMR22E04-GAL4<sup>DBD</sup> | (Jenett et al., 2012) | BDSC: 69199 |
| *D. melanogaster*: VT028327-p65<sup>AD</sup> | (Tirian et al., 2017) | BDSC: 73064 |
| *D. melanogaster*: GMR31F09-GAL4<sup>DBD</sup> | (Dionne et al., 2018) | BDSC: 68759 |
| *D. melanogaster*: GMR78H05-p65<sup>AD</sup> | (Dionne et al., 2018) | BDSC: 70814 |
| *D. melanogaster*: Mz19-GAL4 | (Ito et al., 1998) | BDSC: 34497 |
| *D. melanogaster*: QUAS-mtdTomato-3xHA | (Potter et al., 2010) | BDSC: 30004 |
| *D. melanogaster*: Or47b-rCD2 | (Zhu and Luo., 2018) | BDSC: 9916 |
| *D. melanogaster*: trans-Tango | (Talay et al., 2017) | BDSC: 77123 |
| *D. melanogaster*: UAS-dcr2 | (Dietzl et al., 2007) | N/A |
| *D. melanogaster*: UAS-mCD8-GFP | (Lee and Luo., 1999) | DGRC: 108068 |
| *D. melanogaster*: hsFLP | (Golic and Lindquist., 1989) | N/A |
| *D. melanogaster*: NP-6658-GAL4 | (Hong et al., 2012) | BDSC: 41568 |
| *D. melanogaster*: P{GS}9267 | (Hong et al., 2012) | BDSC: 41567 |
| *D. melanogaster*: QUAS-Ten-m | (Hong et al., 2012) | BDSC: 41571/41572 |
| *D. melanogaster*: UAS-myr-mGreenLantern | (Wong et al., 2023) | N/A |
| *D. melanogaster*: UAS-Syd1-WT-3xFLAG | (Spinner et al., 2018) | N/A |
| *D. melanogaster*: UAS-Syd1-R979A-3xFLAG | (Spinner et al., 2018) | N/A |
| *D. melanogaster*: UAS-Gek | (Gontang et al., 2011) | N/A |
| *D. melanogaster*: UAS-Gek-K129A | (Gontang et al., 2011) | N/A |
| *D. melanogaster*: Mz19-QF2<sup>G4HACK</sup> | this study | N/A |
| *D. melanogaster: UAS-Halo-alphaTub84B* | this study | N/A |
| *D. melanogaster: UAS-Halo-EB1* | this study | N/A |
| *D. melanogaster: UAS-Halo-Moesin* | this study | N/A |
| *D. melanogaster: UAS-V5-Ten-m* | this study | N/A |
| *D. melanogaster: UAS-V5-Ten-m-ΔECD* | this study | N/A |
| *D. melanogaster: UAS-V5-Ten-m-ΔICD* | this study | N/A |
| *D. melanogaster: APEX2-V5-Ten-m* | this study | N/A |
| *D. melanogaster: UAS-CD4-APEX2* | this study | N/A |
| *D. melanogaster: UAS-Gek-FLAG* | this study | N/A |
| *D. melanogaster: UAS-Gek-K129A-FLAG* | this study | N/A |
| *D. melanogaster: UAS-V5-Ten-m (RNAi-resistant)* | this study | N/A |
| *D. melanogaster: UAS-V5-Ten-m-ΔICD (RNAi-resistant),* | this study | N/A |
| *D. melanogaster: VT028327-FRT10-STOP-FRT10-p65AD* | this study | N/A |
| *D. melanogaster: UAS-RNAi lines* | (Dietzl et al., 2007; Ni et al., 2011; Perkins et al., 2015) | Stock numbers listed in **Table S3** |
| **Recombinant DNA** |  |  |
| *pUAS-FRT10-stop-FRT10-mCD8-GFP* | (Li et al., 2021) | N/A |
| *pCR-Blunt-TOPO* | Thermo Fisher Scientific | Catalog #: K280020 |
| *pU6-BbsI-chiRNA* | (Gratz et al., 2014; Gratz et al., 2013) | Addgene: 45946 |
| *UAS-Halo-CAAX* | (Sutcliffe et al., 2017) | Addgene: 87645 |
| *pJFRC81-10xUAS-IVS-Syn21-GFP-p10* | (Pfeiffer et al., 2012) | Addgene: 36432 |
| *UAS-CD4-GFP* | (Han et al., 2011) | N/A |
| *pUAST-attB-Ten-m* | (Hong et al., 2012) | N/A |
| Other tagged *UAS-Ten-m* constructs | this study | N/A |
| tagged *UAS-Gek* or *-Syd1* constructs | this study | N/A |
| *VT028327-FRT10-STOP-FRT10-p65AD* construct | this study | N/A |
| **Software and Algorithms** |  |  |
| Zen | Carl Zeiss | RRID: SCR_013672 |
| ImageJ | National Institutes of Health | RRID: SCR_003070 |
| Illustrator | Adobe | RRID: SCR_010279 |
| Spectrum Mill | Agilent | https://proteomics.broadinstitute.org/ |

## RESOURCE AVAILABILITY

### Lead contact

Further information and requests for resources and reagents should be directed to the lead contact, Liqun Luo.

### Materials availability

All unique reagents generated in this study are available from the lead contact.

### Data and code availability

The original mass spectra and the protein sequence database used for searches have been deposited in the public proteomics repository MassIVE (http://massive.ucsd.edu) and are accessible at ftp://MSV000094010@massive.ucsd.edu when providing the dataset password: teneurin. If requested, also provide the username: MSV000094010. These datasets will be made public upon acceptance of the manuscript. Processed proteomic data is provided in **Table S1**.

## EXPERIMENTAL MODEL AND SUBJECT DETAILS

### *Drosophila* stocks and genotypes

Flies were raised on standard cornmeal medium in a 12h/12h light cycle at 25°C. To increase transgene expression, 29°C was used for some experiments as specified in the figure legend. Complete genotypes of flies in each experiment are described in **Table S3**. The following lines were used: *GMR22E04-GAL4^DBD^* (the DBD of the DA1-ORN split GAL4)^111^, *VT028327-p65^AD^* (the AD of the DA1-ORN split GAL4)^127^, *GMR31F09-GAL4^DBD^* and *GMR78H05-p65^AD^* (the DBD and the AD of the VA1d-ORN split GAL4)^113^, *Mz19-GAL4*^115^, *QUAS-mtdTomato-3xHA*^117^, *Or47b-rCD2*^119^, *trans-*Tango^42^, *UAS-dcr2*^112^, *UAS-mCD8-GFP*^106^, *hsFLP* (*heat shock protein* promoter-driven *FLP*)^122^, *NP-6658-GAL4* (*Ten-m-GAL4*)^5^*, P{GS}9267 (UAS*-gated *Ten-m* overexpression), and *QUAS-Ten-m*^5^, and *UAS-myr-mGreenLantern*^39^. *UAS-Syd1-WT-3xFLAG* and *UAS-Syd1-R979A-3xFLAG*^55^ were kindly provided by the Herman lab (University of Oregon), *UAS-Gek and UAS-Gek-K129A*^63^ used for early experiments were kindly provided by the Clandinin lab (Stanford University). The DA1-ORN lines, one with Ten-m overexpression and one without (**Figure 1, S1, 3, and S4**), were generated in this study. The Mz19-PN line with Ten-m overexpression (**Figure 4**) was generated based on the previously built Mz19-PN genetic screen line^5^. The VA1d-ORN line (**Figure 5**) is an unpublished reagent generously provided by Cheng Lyu. *Mz19-QF2^G4HACK^, UAS-Halo-alphaTub84B, UAS-Halo-EB1*, *UAS-Halo-Moesin*, *UAS-V5-Ten-m*, *UAS-V5-Ten-m-ΔECD*, *UAS-V5-Ten-m-ΔICD*, *APEX2-V5-Ten-m, UAS-CD4-APEX2*, *UAS-Gek-FLAG, UAS-Gek-K129A-FLAG*, *UAS-V5-Ten-m (RNAi-resistant)*, *UAS-V5-Ten-m-ΔICD (RNAi-resistant)*, and *VT028327-FRT10-STOP-FRT10-p65AD* were generated in this study. The other RNAi or overexpression lines were generated previously^112,114,116^ and acquired from the Bloomington *Drosophila* Stock Center and the Vienna *Drosophila* Resource Center (stock numbers listed in **Table S3**).

## METHOD DETAILS

### Generation of *APEX2-V5-Ten-m* flies

The *APEX2-V5-Ten-m* fly line was generated by CRISPR-mediated knock-in to the *Ten-m* genomic locus. Briefly, to build the homology-directed repair (HDR) vector, a ∼1500bp genomic sequence flanking the *Ten-m* start codon (∼750bp each side) was amplified using the Q5 hot-start high-fidelity DNA polymerase (New England Biolabs) and inserted into the *pCR-Blunt-TOPO* vector (Thermo Fisher). The codon-optimized APEX2-V5 sequence was synthesized as a gBlock (Integrated DNA Technologies) and inserted into the *TOPO genomic sequence plasmid* using the NEBuilder HiFi DNA assembly master mix (New England Biolabs). CRISPR guide RNA (gRNA) targeting a locus near the start codon was designed using the flyCRISPR Target Finder web tool^118,120,121^ and cloned into the *pU6-BbsI-chiRNA* vector^123^ (Addgene #45946) by NEBuilder HiFi DNA assembly master mix. Silent mutations were introduced at the PAM site of the HDR vector by using the Q5 site-directed mutagenesis kit (New England Biolabs). The *APEX2-V5-Ten-m* HDR and the *Ten-m* gRNA vectors were co-injected into *vas-Cas9*^124^ fly embryos by BestGene. G0 flies were crossed to a third chromosome balancer line and all progenies were individually balanced and genotyped until APEX2-insertion-positive candidates were identified. APEX2-insertion-positive candidates were sequenced and then kept.

### Generation of UAS constructs and transgenic flies

To generate the *UAS-CD4-APEX2-V5* construct, the signal peptide from the Drosophila *akh* gene, the CD4 coding sequence from *UAS-CD4-GFP*^125^, and the codon-optimized APEX2-V5 sequence (see above) were amplified using the Q5 hot-start high-fidelity DNA polymerase (New England Biolabs) and inserted into the *pJFRC81-10xUAS-IVS-Syn21-GFP-p10*^126^ vector (Addgene #36432) to replace the GFP sequence using NEBuilder HiFi DNA assembly master mix (New England Biolabs).

To generate the *UAS-Gek-FLAG* and *UAS-Syd1-FLAG* constructs, we extracted the total RNA of *w1118* pupal fly heads using an RNA mini-prep kit (Zymo Research), synthesized the complementary DNA using the SuperScript III First-Strand Synthesis SuperMix (Thermo Fisher), and amplified the *Gek* or *Syd1* coding sequences using the Q5 hot-start high-fidelity DNA polymerase (New England Biolabs). The verified coding sequences were then assembled into a modified *pUAST-attB* vector, in which a FLAG tag was added at the 3’ end.

To generate the *UAS-Gek-K129A-FLAG* construct, the *K129A* mutation was introduced using the Q5 site-directed mutagenesis kit (New England Biolabs).

To generate the *UAS-V5-Ten-m* construct, a V5 tag was inserted after the start codon of Ten-m cDNA (isoform B) in the plasmid *pUAST-attB-Ten-m*^5^ using the Q5 site-directed mutagenesis kit (New England Biolabs). To generate the *UAS-V5-Ten-m-ΔICD* and *UAS-V5-Ten-m-ΔECD* constructs, N2-A225 and I256-A2731 were deleted using the NEBuilder HiFi DNA assembly master mix (New England Biolabs), respectively.

To generate the RNAi^VDRC330540^-resistant *UAS-V5-Ten-m* and *UAS-V5-Ten-m-ΔICD* constructs, mutations (**Figure 5I**) were introduced using the Q5 site-directed mutagenesis kit (New England Biolabs).

To generate the *UAS-V5-Ten-m-FLAG*, *UAS-V5-Ten-m-ΔICD-FLAG*, and *UAS-V5-Ten-m-ΔECD-FLAG* constructs, a FLAG tag was inserted before the stop codon of the corresponding V5-tagged constructs described above.

To generate the *VT02832-p65AD* construct, VT027328 primers^127^ were used to amplify the sequence from the genomic DNA of *VT027328-p65AD* fly line (BDRC #73064) The verified sequence was then assembled into the pENTR/D-TOPO vector (Thermo Fisher) and integrated into the pBPp65ADZpUw vector using the Gateway LR Clonase II Enzyme mix (Thermo Fisher).

To generate the *VT028327-FRT10-STOP-FRT10-p65AD* construct, the *FRT10-STOP-FRT10* sequence^64^ and the T2A element were inserted after the p65AD start codon of the *VT02832-p65AD* construct. Each plasmid was verified by full-length DNA sequencing.

To generate *UAS-Halo-Moesin*, *UAS-Halo-EB1*, and *UAS-Halo-alphaTub84B*, the moesin actin binding domain, EB1, and alphaTub84B coding sequences were PCR amplified from the genomic DNA of transgenic flies *UAS-GMA* (BDRC #31775), *UAS-EB1-GFP* (BDRC #35512), and *UAS-GFP-alphaTub84B* (BDRC #7373), respectively, and subcloned into *UAS-Halo-CAAX* (Addgene #87645) using XhoI and XbaI. The *pUAST-attB* constructs were inserted into the *attP24* (for *Gek* constructs and *UAS-CD4-APEX2*), *VK00027* (for *VT028327-FRT10-STOP-FRT10-p65AD*), *VK00019* (for cytoskeleton marker constructs), or *attP86Fb* (for *Ten-m* constructs) landing sites.

Transgenic flies were generated in house by standard methods involving microinjection of DNA into early *Drosophila* embryos prior to cellularization. G0 flies were crossed to a *white^–^* balancer, and all *white^+^* progenies were individually balanced and verified.

### APEX2-mediated proximity biotinylation in fly brains

The proximity labeling reaction was performed following the previously published method^46^. Briefly, we dissected APEX2-Ten-m group, spatial reference group, and negative control group in pre-chilled Schneider’s medium (Thermo Fisher) and transferred them into 500 μL of the Schneider’s medium in 1.5 mL protein low-binding tubes (Eppendorf) on ice. Brains were washed with the Schneider’s medium to remove fat bodies and debris and were incubated in 100 μM of biotin-phenol (BP; APExBIO) in the Schneider’s medium on ice for one hour, with occasional pipetting for mixing. Brains were then labeled with 1 mM (0.003%) H_2_O_2_ (Thermo Fisher) for 1 minute, and immediately quenched by five thorough washes using the quenching buffer that contains 10 mM sodium ascorbate (Spectrum Chemicals), 5 mM Trolox (Sigma-Aldrich), and 10 mM sodium azide (Sigma-Aldrich) in phosphate buffered saline (PBS; Thermo Fisher). After the washes, the quenching solution was removed, and brains were either fixed for immunostaining (see below for details) or were frozen in liquid nitrogen and stored at –80°C for proteomic analysis. For proteomic sample collection, 900 dissected and biotinylated brains were collected for each experimental group (5400 brains in total).

### Enrichment of biotinylated proteins

Brains were processed in the original collection tube, to avoid loss during transferring. We added 40 μL of high-SDS RIPA (50mM Tris-HCl [pH 8.0], 150 mM NaCl, 1% sodium dodecyl sulfate [SDS], 0.5% sodium deoxycholate, 1% Triton X-100, 1x protease inhibitor cocktail [Sigma-Aldrich], and 1 mM phenylmethylsulfonyl fluoride [PMSF; Sigma-Aldrich]) to each tube of frozen brains, and grinded the samples on ice using disposable pestles with an electric pellet pestle driver. Tubes containing brain lysates of the same group were spun down, merged, and rinsed with an additional 100 μL of high-SDS RIPA to collect remaining proteins. Samples were then vortexed briefly, sonicated twice for ten seconds each, and incubated at 95°C for five minutes to denature proteins. 1.2 mL of SDS-free RIPA buffer (50 mM Tris-HCl [pH 8.0], 150 mM NaCl, 0.5% sodium deoxycholate, 1% Triton X-100, 1x protease inhibitor cocktail, and 1 mM PMSF) were added to each sample, and the mixture was rotated for two hours at 4°C. Lysates were then diluted with 200 μL of normal RIPA buffer (50 mM Tris-HCl [pH 8.0], 150 mM NaCl, 0.2% SDS, 0.5% sodium deoxycholate, 1% Triton X-100, 1x protease inhibitor cocktail, and 1 mM PMSF), transferred to 3.5 mL ultracentrifuge tubes (Beckman Coulter), and centrifuged at 100,000 g for 30 minutes at 4°C. 1.5 mL of the supernatant was carefully collected for each sample. 400 μL of streptavidin magnetic beads (Pierce) washed twice using 1 ml RIPA buffer were added to each of the post-ultracentrifugation brain lysates. The lysate and the streptavidin bead mixture were left to rotate at 4°C overnight. On the following day, beads were washed twice with 1 mL RIPA buffer, once with 1 mL of 1 M KCl, once with 1 mL of 0.1 M Na_2_CO_3_, once with 1 mL of 2 M urea in 10 mM Tris-HCl (pH 8.0), and again twice with 1 mL RIPA buffer. The beads were resuspended in 1 mL fresh RIPA buffer. 35 μL of the bead suspension was taken out for western blot, and the rest proceeded to on-bead digestion.

### Western blotting of biotinylated proteins

Biotinylated proteins were eluted from streptavidin beads by the addition of 20 μL of elution buffer (2X Laemmli sample buffer [Bio-Rad], 20 mM dithiothreitol [Sigma-Aldrich], and 2 mM biotin [Sigma-Aldrich]) followed by a 10 min incubation at 95°C. Proteins were resolved by 4%–12% Bis-Tris PAGE gels (Thermo Fisher) and transferred to nitrocellulose membranes (Thermo Fisher). After blocking with 3% bovine serum albumin (BSA) in Tris-buffered saline with 0.1% Tween 20 (TBST; Thermo Fisher) for 1 hour, membrane was incubated with 0.3 mg/mL HRP-conjugated streptavidin for one hour. The Clarity Western ECL blotting substrate (Bio-Rad) and ChemiDoc imaging system (Bio-Rad) were used to develop and detect chemiluminescence.

### On-bead trypsin digestion of biotinylated proteins

The streptavidin-enriched sample (400 μL of streptavidin beads per condition) was processed for on-bead digestion and TMT labeling and used for mass spectrometry analysis as previously described^46^. Proteins bound to streptavidin beads were washed twice with 200 μL of 50 mM Tris-HCl buffer (pH 7.5), followed by two washes with 2 M urea/50 mM Tris (pH 7.5) buffer in fresh tubes. The final volume of 2 M urea/50 mM Tris (pH 7.5) buffer was removed, and beads were incubated with 80 μL of 2 M urea/50 mM Tris buffer containing 1 mM dithiothreitol (DTT) and 0.4 μg trypsin. Beads were incubated in the urea/trypsin buffer for 1 hour at 25°C while shaking at 1000 revolutions per minute (rpm). After 1 hour, the supernatant was removed and transferred to a fresh tube. The streptavidin beads were washed twice with 60 μL of 2 M urea/50 mM Tris (pH 7.5) buffer and the washes were combined with the on-bead digest supernatant. The eluate was reduced with 4 mM DTT for 30 min at 25°C with shaking at 1000 rpm. The samples were alkylated with 10 mM iodoacetamide and incubated for 45 min in the dark at 25°C while shaking at 1000 rpm. An additional 0.5 μg of trypsin was added to the sample and the digestion was completed overnight at 25°C with shaking at 700 rpm. After overnight digestion, the sample was acidified (pH < 3) by adding formic acid (FA) such that the sample contained 1% FA. Samples were desalted on C18 StageTips (3M). Briefly, C18 StageTips were conditioned with 100 μL of 100% MeOH, 100 μL of 50% MeCN/0.1% FA, and 2x with 100 μL of 0.1% FA. Acidified peptides were loaded onto the conditioned StageTips, which were subsequently washed 2 times with 100 μL of 0.1% FA. Peptides were eluted from StageTips with 50 μL of 50% MeCN/0.1% FA and dried to completion.

### TMT labeling and StageTip peptide fractionation

Desalted peptides were labeled with TMT6 reagents (Thermo Fisher Scientific) as directed by the manufacturer. Peptides were reconstituted in 100 μL of 50 mM HEPES. Each 0.8 mg vial of TMT reagent was reconstituted in 41 μL of anhydrous acetonitrile and added to the corresponding peptide sample for 1 hour at room temperature shaking at 1000 rpm. Labeling of samples with TMT reagents was completed with the design described in **Figure 2F**. TMT labeling reactions were quenched with 8 μL of 5% hydroxylamine at room temperature for 15 min with shaking. The entirety of each sample was pooled, evaporated to dryness in a vacuum concentrator, and desalted on C18 StageTips as described above. One SCX StageTip was prepared per sample using 3 plugs of SCX material (3M) topped with 2 plugs of C18 material. StageTips were sequentially conditioned with 100 μL of MeOH, 100 μL of 80% MeCN/0.5% acetic acid, 100 μL of 0.5% acetic acid, 100 μL of 0.5% acetic acid/500mM NH_4_AcO/20% MeCN, followed by another 100 μL of 0.5% acetic acid. Dried sample was re-suspended in 250 μL of 0.5% acetic acid, loaded onto the StageTips, and washed twice with 100 μL of 0.5% acetic acid. Sample was transeluted from C18 material onto the SCX with 100 μL of 80% MeCN/0.5% acetic acid, and consecutively eluted using 3 buffers with increasing pH—pH 5.15 (50mM NH_4_AcO/20% MeCN), pH 8.25 (50mM NH_4_HCO_3_/20% MeCN), and finally pH 10.3 (0.1% NH_4_OH, 20% MeCN). Three eluted fractions were re-suspended in 200 μL of 0.5% acetic acid to reduce the MeCN concentration and subsequently desalted on C18 StageTips as described above. Desalted peptides were dried to completion.

### Liquid chromatography and mass spectrometry

Desalted TMT-labeled peptides were resuspended in 9 μL of 3% MeCN, 0.1% FA and analyzed by online nanoflow liquid chromatography tandem mass spectrometry (LC-MS/MS) using a Q Exactive Plus (for fractionated samples) (Thermo Fisher Scientific) coupled on-line to a Proxeon Easy-nLC 1200 (Thermo Fisher Scientific). 4 μL of each sample were loaded at 500 nL/min onto a microcapillary column (360 μm outer diameter x 75 μm inner diameter) containing an integrated electrospray emitter tip (10 mm), packed to approximately 28 cm with ReproSil-Pur C18-AQ 1.9 mm beads (Dr. Maisch GmbH) and heated to 50°C. The HPLC solvent A was 3% MeCN, 0.1% FA, and the solvent B was 90% MeCN, 0.1% FA. Peptides were eluted into the mass spectrometer at a flow rate of 200 nL/min. The SCX fractions were run with 110-minute method, which used the following gradient profile: (min:%B) 0:2; 1:6, 85:30; 94:60; 95:90, 100:90,101:50,110:50 (the last two steps at 500 nL/min flow rate). The Q Exactive Plus was operated in the data-dependent mode acquiring HCD MS/MS scans (r = 17,500) after each MS1 scan (r = 70,000) on the top 12 most abundant ions using an MS1 target of 3E6 and an MS2 target of 5E4. The maximum ion time utilized for MS/MS scans was 105 ms; the HCD normalized collision energy was set to 31; the dynamic exclusion time was set to 30 s, and the peptide match was set to “preferred” and isotope exclusion functions were enabled. Charge exclusion was enabled for charge states that were unassigned, 1, 7, 8, >8.

### Mass spectrometry data processing

Collected data were analyzed using the Spectrum Mill software package (proteomics.broadinstitute.org). Nearby MS scans with a similar precursor m/z were merged if they were within ±60 s retention time and ±1.4 m/z tolerance. MS/MS spectra were excluded from searching if they failed the quality filter by not having a sequence tag length 0 or did not have a precursor MH+ in the range of 750–4000. All extracted spectra were searched against a UniProt database containing *Drosophila melanogaster* reference proteome sequences. Search parameters included: ESI QEXACTIVE-HCD-v2 scoring parent and fragment mass tolerance of 20 ppm, 40% minimum matched peak intensity, trypsin allow P enzyme specificity with up to two missed cleavages, and calculate reversed database scores enabled. Fixed modifications were carbamidomethylation at cysteine. TMT labeling was required at lysine, but peptide N termini were allowed to be either labeled or unlabeled. Allowed variable modifications were protein N-terminal acetylation and oxidized methionine. Individual spectra were automatically assigned a confidence score using the Spectrum Mill auto-validation module. Score at the peptide mode was based on a target-decoy false discovery rate (FDR) of 1%. Protein polishing auto-validation was then applied using an auto thresholding strategy. Relative abundances of proteins were determined using TMT reporter ion intensity ratios from each MS/MS spectrum and the median ratio was calculated from all MS/MS spectra contributing to a protein subgroup. Proteins identified by 2 or more distinct peptides and ratio counts were considered for the dataset.

### Linear model for the mass spectrometry data

Starting with the processed mass spectrometry data, we developed a linear model to identify prospective interacting partners of Ten-m. Using the log_2_ transformed TMT ratios, the linear model is as follows:

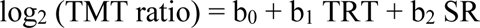

where TRT and SR are indicator variables representing APEX2-Ten-m enrichment and spatial reference, respectively. The negative control NC constitutes the baseline for the model. The [Ten-m/SR fold change] taking negative controls into account is represented by the (b_1_ – b_2_) contrast while the [Ten-m/NC fold change] is captured by the b_1_ coefficient. The model is fitted using an empirical Bayes approach and the relevant contrasts/coefficients are subject to a moderated t-test to determine nominal *p*-values for each protein in the TMT dataset. These nominal *p*-values are then corrected for multiple testing using the Benjamini-Hochberg FDR (BH-FDR) method^128^. The linear model along with the associated moderated t-test and BH-FDR correction were implemented using the limma library^129^ in R.

### Proteomic data analysis

To identify prospective interacting partners of Ten-m, we implemented three filtering steps: (1) From the total of 3454 proteins detected across 6 samples, we focused on those with at least two unique peptides, narrowing the list down to 2854 proteins. (2) We then filtered out potential contaminants, including endogenously biotinylated and endogenous peroxidase-labeled proteins, by using the [APEX2-Ten-m/NC] fold change of the Ten-m protein itself as a threshold, resulting in 781 proteins. (3) Finally, to exclude generic proteins located near the cell membrane, we employed a [APEX2-Ten-m/SR] fold change–based ratiometric approach, isolating 294 proteins specifically enriched by APEX2-Ten-m. Functional enrichment analyses, including Gene Ontology, protein domain (SMART), reactome pathway, and local network cluster, were performed on these gene sets using the STRING database.

### Immunocytochemistry

Fly brains were dissected and immunostained according to the previously published protocol^130^. Briefly, brains were dissected in pre-cooled PBS (phosphate buffered saline; Thermo Fisher) and then fixed in 4% paraformaldehyde (Electron Microscopy Sciences) in PBS with 0.015% Triton X-100 (Sigma-Aldrich) for 20 minutes (15 minutes for sparse axon experiments to prevent over-fixation background) on a nutator at room temperature. Fixed brains were washed with PBST (0.3% Triton X-100 in PBS) four times, each time nutating for 15 minutes. The brains were then blocked in 5% normal donkey serum (Jackson ImmunoResearch) in PBST for 1 hour at room temperature or overnight at 4°C on a nutator. Primary antibodies were diluted in the blocking solution and incubated with brains for 36-48 hours on a 4°C nutator. After washed with PBST four times, each time nutating for 20 minutes, brains were incubated with secondary antibodies diluted in the blocking solution and nutated in the dark for 24-48 hours at 4°C. Brains were then washed again with PBST four times, each time nutating for 20 minutes. Immunostained brains were mounted with the SlowFade antifade reagent (Thermo Fisher) and stored at 4°C before imaging.

Primary antibodies used in immunostaining include: rat anti-NCad (1:40; DN-Ex#8, Developmental Studies Hybridoma Bank), mouse anti-BRP (1:80; nc82, Developmental Studies Hybridoma Bank), chicken anti-GFP (1:1000; GFP-1020, Aves Labs), rabbit anti-DsRed (1:500; 632496, Clontech), mouse anti-rat CD2 (1:200; OX-34, Bio-Rad), mouse anti-V5 (1:200; R960-25, Thermo Fisher), rat anti-V5 (1:200; ab206571, Abcam), and mouse anti-Halo (1:100; G9211, Promega). Donkey secondary antibodies conjugated to Alexa Fluor 405/488/568/647 (Jackson ImmunoResearch or Thermo Fisher) were used at 1:250. Neutravidin pre-conjugated with Alexa Fluor 647 (1:1000; synthesized in the Ting lab) was used to detect biotin.

### HaloTag labeling

Fly brains were labelled according to the previously published protocol^131^. Janelia Fluor (JF) HaloTag dyes (stocks at 1 mM) were gifts from the Lavis lab^132,133^. Briefly, fly brains were dissected in pre-cooled PBS and then fixed in 4% paraformaldehyde in PBS for 15 minutes on a nutator at room temperature. Fixed brains were washed with PBST for 5 min, repeated 3 times, followed by incubation with JF646-HaloTag ligand (1:2500 diluted in PBS) for 1 h or overnight at room temperature in the dark. Brains were then washed with PBST for 5 min, repeated 3 times, followed by immunostaining protocol if necessary.

### Transfection and immunostaining of *Drosophila* S2 cells

S2 cells (Thermo Fisher) were cultured in the Schneider’s medium (Thermo Fisher) following the manufacturer’s protocol. S2 cells were transfected with *Actin-GAL4,* along with *UAS-V5-Ten-m-FLAG*, *UAS-V5-Ten-m-ΔICD-FLAG*, or *UAS-V5-Ten-m-ΔECD-FLAG* constructs using the FuGENE HD transfection Reagent (Promega). After 48 hours, transfected cells were transferred to coverslips pre-coated with Concanavalin A (Sigma-Aldrich). For the plasma membrane non-permeabilized condition, S2 cells were incubated with rat anti-V5 antibody (1:200; Abcam) and mouse anti-FLAG M2 antibody (1:200; Sigma-Aldrich) diluted in the Schneider’s medium (Thermo Fisher) at room temperature for 1 hour. S2 cells were rinsed with PBS, fixed with 4% PFA in PBST, washed with PBST, blocked with 5% normal donkey serum (Jackson ImmunoResearch) in PBST, incubated with secondary antibodies in the dark, washed with PBST, mounted, and imaged. For the plasma membrane permeabilized condition, S2 cells were incubated in the Schneider’s medium at room temperature for 1 hour, rinsed with PBS, fixed with 4% PFA in PBST, washed with PBST, blocked with 5% normal donkey serum in PBST, incubated with primary antibodies, washed with PBST, incubated with secondary antibodies in the dark, washed with PBST, mounted, and imaged.

### Image acquisition and processing

Images were obtained using laser scanning confocal microscopy (Zeiss LSM 780 or LSM 900). Brightness and contrast adjustments as well as image cropping were done using ImageJ.

### Co-immunoprecipitation assay

S2 cells (Thermo Fisher) were cultured in the Schneider’s medium (Thermo Fisher) following the manufacturer’s protocol. S2 cells were transfected with *UAS-Syd1-FLAG* or *UAS-Gek-FLAG,* along with a *Ten-m* expression construct and *Actin-GAL4* using the FuGENE HD transfection reagent (Promega). After 72 hours, the transfected cells were harvested, rinsed with PBS, lysed in the lysis buffer (50 mM Tris-HCl [pH 7.5], 150 mM NaCl, 0.2% TritonX-100) supplemented with protease inhibitor cocktail (Promega). The cell lysates were rotated at 4°C for 30 minutes and sonicated on ice for a total of 1 minute with a “10-second ON, 10-second OFF” cycle, using 48 W of power. The cell lysates were then centrifuged at 15,000 g for 20 minutes at 4°C. The supernatants were collected and incubated with Dynabeads Protein G beads (Thermo Fisher) pre-coated with the mouse anti-V5 antibody (1:100; R960-25, Thermo Fisher) and then left to rotate at 4°C overnight. On the following day, the samples were washed extensively in wash buffer (50 mM Tris-HCl [pH 7.5], 150 mM NaCl, 0.5% TritonX-100) for three times, 10 minutes each. The proteins were eluted from beads by adding the loading buffer (4X Laemmli sample buffer [Bio-Rad] with 20 mM dithiothreitol) followed by a 10 min incubation at 95°C. The samples were loaded in 3%–8% Tris-Acetate PAGE gels (Thermo Fisher) for protein electrophoresis and transferred to PVDF membranes (Thermo Fisher) according to the manufacturer’s protocol. The membranes were blocked with the SuperBlock blocking buffer (Thermo Fisher), incubated with mouse anti-FLAG M2 antibody (1:3000; Sigma-Aldrich), washed with TBST (Thermo Fisher), incubated with light chain specific HRP-conjugated secondary antibodies (1:5000; Jackson ImmunoResearch), washed with TBST, and developed with Clarity Western ECL blotting substrate (Bio-Rad).

### Sparse axon labeling and genetic manipulation

Each fly contains the DA1-ORN sparse driver and its reporter (*UAS-myr-mGreenLantern, UAS-mCD8-GFP*, *VT028327-FRT10-STOP-FRT10-p65AD*, *GMR22E04-GAL4^DBD^*), *hsFLP*, the DA1-PN driver and its reporter (*Mz19-QF2^G4HACK^*, *QUAS-mtdTomato-3xHA*), and other desired UAS constructs for genetic manipulation (*UAS-V5-Ten-m or UAS-dcr2*, *UAS-Rac1-RNAi^BDRC28985^*). For sparse axon experiments imaging F-actin distribution, *UAS-Halo-Moesin* is also included. Complete fly genotypes of sparse axon experiments are described in **Table S3**. Flies were raised on standard cornmeal medium in a 12h/12h light cycle at 25°C (avoiding using 29°C to prevent any leakiness of *hsFLP*). Early-stage pupae (0–6 hours APF) were collected on a single layer of water-soaked paper towel (avoiding air bubbles to prevent inefficient heat transmission), heat shocked for 30 seconds in a 37°C water bath, and then immediately cooled for 60 seconds in a room temperature water bath (**Figure S6B**). Flies were dissected at 28–34, 34–40, 40–46 hours APF for Stage 1, Stage 2, and Stage 3, respectively. For fly stocks containing the sparse driver, *VT028327-FRT10-STOP-FRT10-p65AD* (or any other sparse drivers) and *hsFLP* are kept in separate stocks to avoid stochastic FLP expression and subsequent loss of the *FRT10-STOP-FRT10* cassette. Heat shock duration was empirically determined according to the intended number of cells, developmental stage, and tissue depth. If achieving the desired sparsity proves difficult, consider replacing the *FRT10-STOP-FRT10* element with a less sensitive *FRT100-STOP-FRT100* element^64^ in the sparse driver design.

## QUANTIFICATION AND STATISTICAL ANALYSIS

### Quantification of match indices for DA1-ORNs

*Mz19-QF2^G4HACK^*-driven *QUAS-mtdTomato-3xHA* specifically labels DA1-PNs in most cases. Antennal lobes with occasional *Mz19-QF2^G4HACK^*-driven VA1d/DC3-PN labeling (cell bodies located dorsal to antennal lobe, rather than lateral for DA1-PNs) were excluded to prevent ambiguity in DA1-PN dendrite identification. DA1-ORN split GAL4–driven *UAS-mCD8-GFP* was used for DA1-ORN axon identification. “Match index” is defined as the ratio of the overlapping volume between DA1-ORN axons and DA1-PN dendrites to the total volume of DA1-PN dendrites. Data was analyzed using ImageJ (Fiji) 3D object counter and plotted using R. Data normality was assessed using the Shapiro-Wilk normality test. The Brown-Forsythe test was used to assess homoscedasticity prior to the ANOVA. For data with normal distribution and equal variance, the one-way ANOVA with Tukey’s test was used for multiple comparisons. Otherwise, the Kruskal-Wallis test with Bonferroni post-hoc correction was used for multiple comparisons.

### Quantification of V5 signal intensities of Ten-m expression in DA1-ORNs

The average V5 signal intensity of each DA1 glomerulus was measured and then normalized against the maximum and minimum signal intensities within each image. Maximum signal intensities were primarily contributed by background signals from the trachea, whose intensity is consistent across fly brains. The Kruskal-Wallis test with Bonferroni post-hoc correction was used for multiple comparisons.

### Quantification of mismatch indices in Mz19-PNs

*Mz19-GAL4*-driven *UAS-mCD8-GFP* was used for Mz19-PN dendrite identification, while *Or47b-rCD2* was used for VA1v-ORN axon identification. “Mismatch index” is defined as the ratio of the overlapping volume between VA1v-ORN axons and Mz19-PN dendrites to the total volume of VA1v-ORN axons. Data was analyzed using ImageJ (Fiji) 3D object counter and plotted using R. The Kruskal-Wallis test with Bonferroni post-hoc correction was used for multiple comparisons.

### Quantification of mistarget indices in VA1d-ORNs

*Mz19-QF2^G4HACK^*-driven *QUAS-mtdTomato-3xHA* and the NCad staining were used to identify DA1 and VA1d glomeruli. VA1d-ORN split GAL4-driven *UAS-mCD8-GFP* was used for VA1d-ORN axon identification. “Mistarget index” is defined as the ratio of the total GFP fluorescence intensity of axons in the DA1 glomerulus to that in the DA1 and VA1d glomeruli. Data was analyzed using ImageJ (Fiji) and plotted using R. The Kruskal-Wallis test with Bonferroni post-hoc correction was used for multiple comparisons.

### Image processing and quantification of sparse axon assays

Neurite tracing images were generated using Simple Neurite Tracer (SNT)^134^, processed using open-source R package natverse^135^, and analyzed and plotted in R. The stem axon was defined as the thickest segment of the axon. The antennal lobe entry point was determined by the first overlapping point of the axon (identified by GFP staining) and the antennal lobe (identified by NCad staining). The end point was defined as the farthest point of the stem axon from the antennal lobe entry point. The locations of primary branch points were normalized with the antennal lobe entry point set as 0 and the end point as 1. *Mz19-QF2^G4HACK^*-driven *QUAS-mtdTomato-3xHA* was used for DA1-PN dendrite identification. Branches extending to the DA1-PN dendrite region were categorized as “DA1-PN-contacting”. The chi-squared test with Bonferroni correction (**Figure 6L** and **6M**) and the one-way ANOVA with Tukey’s test (**Figure 6N–Q**) were used for multiple comparisons. Signal intensities of the F-actin marker (Halo-Moesin) along branches, traced using Simple Neurite Tracer, were quantified using ImageJ and normalized against the maximum and minimum signal intensities of each image. F-actin densities were then calculated by dividing the total F-actin signal intensities along each branch by its corresponding length. The Mann-Whitney *U* test was used for comparisons.

## ACKNOWLEDGMENTS

We thank members of the Luo lab, especially A. Shuster, B. Zhao, Y. Ge, D. Pederick, H. Ji, T. Hindmarsh Sten, and Y. Wu, as well as S. Yin, K. Ding, T. Südhof, and K. Shen, for their technical support, valuable insights, and feedback on this project and manuscript. We also thank T. Clandinin, T. Herman, the Bloomington *Drosophila* Stock Center, and the Vienna *Drosophila* Resource Center for providing fly lines; Addgene for supplying plasmids; L. Lavis for Janelia Fluor HaloTag dyes; K. Chen and Z. Wang for their advice on biochemical assays; Z. Song and C. Song for their guidance on statistics and programming; and M. Molacavage for administrative assistance. L.L. is an investigator of the Howard Hughes Medical Institute. J.L. is a group leader of the Janelia Research Campus of the Howard Hughes Medical Institute. This work was supported by the National Institutes of Health (R01-DC005982 to L.L. and R01-DK121409 to S.A.C. and A.Y.T.) and the Wu Tsai Neurosciences Institute of Stanford University (Neuro-omics grant to A.Y.T. and L.L.).

## AUTHOR CONTRIBUTIONS

C.X., J.L., and L.L. conceived this project. C.X., J.L., and L.L. designed the proteomic experiments with input from S.H. and A.Y.T. C.X. collected pupae, and Q.X., J.L., H.L., C.X., R.G., and D.J.L. dissected fly brains for the proteomic experiment. C.X., J.L., and S.H. processed proteomic samples. N.D.U., T.S., D.R.M., and S.A.C. performed post-enrichment sample processing, mass spectrometry, and initial data analysis. C.X. and J.L. analyzed proteomic data with input from S.H. and A.Y.T. T.L. generated cytoskeleton marker constructs. Q.X. generated the *Mz19-QF2* fly line. D.J.L. produced transgenic flies. C.X., C.L., and Z.L. screened split GAL4 lines with input from C.N.M. C.X. designed the sparse driver and developed the sparse labeling methodology. C.X. designed, performed, and analyzed all other experiments, receiving input from L.L. and assistance from J.L., Z.L., C.L., Y.H., C.N.M., K.K.L.W., and Y.L. C.X., J.L., and L.L. wrote the manuscript with input from all co-authors.

L.L. supervised all aspects of the work.

---

**Figure S1.**
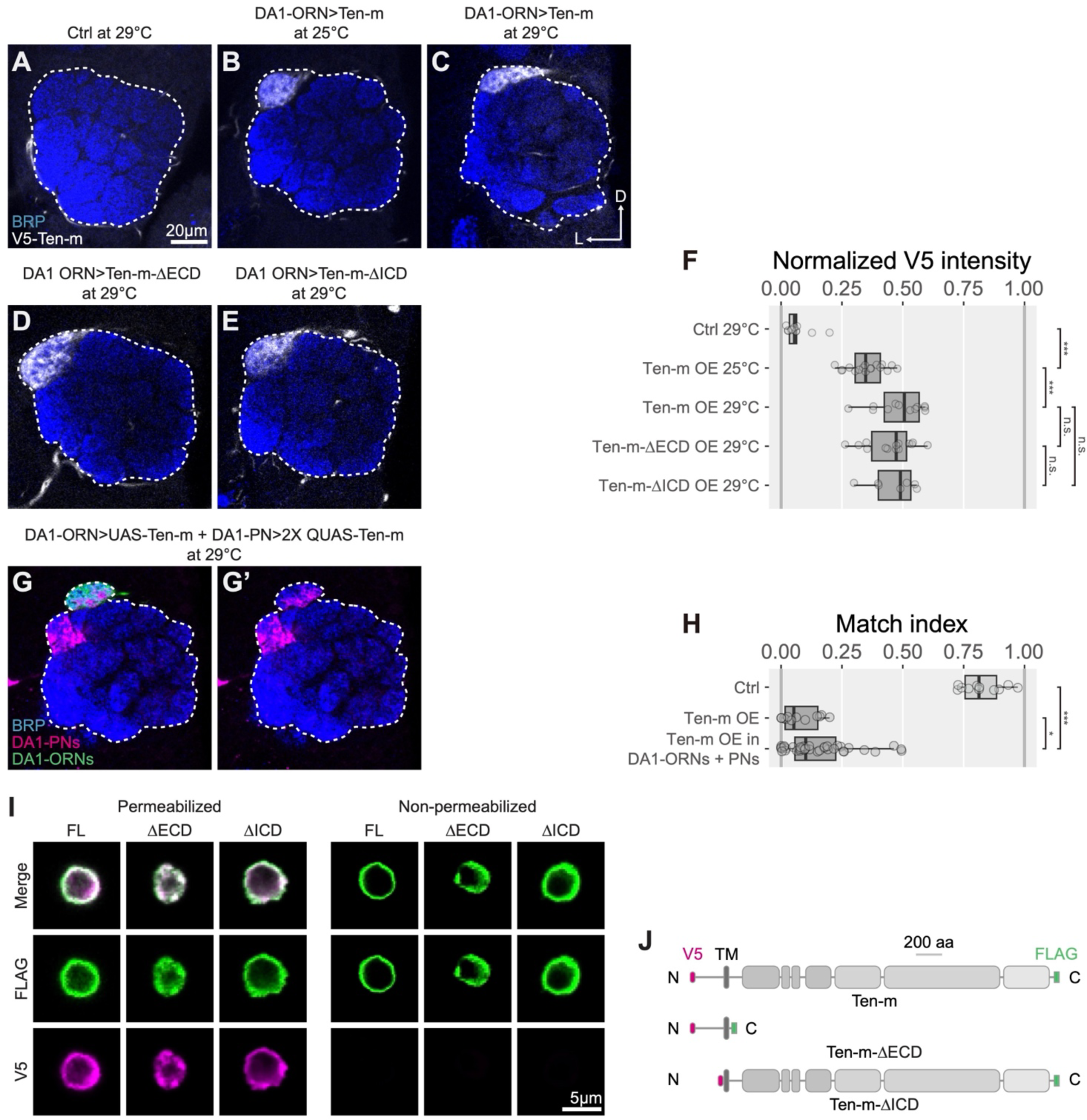
Characterization of Ten-m transgene expression, related to Figure 1. (A–E) V5 staining in representative confocal images of antennal lobes of control at 29°C (A), *UAS-V5-Ten-m* overexpression at 25°C (B), *UAS-V5-Ten-m* overexpression at 29°C (C), *UAS-V5-Ten-m-ΔECD* overexpression at 29°C (D), and *UAS-V5-Ten-m-ΔICD* overexpression at 29°C (E). These are the same brains as shown in Figure 1F–H, 1K, and 1L, respectively. (F) Quantification of normalized V5 intensities. V5 intensities were normalized to the maximum and minimum signal intensities of each image. (G, G’) Representative confocal image of Ten-m overexpression in both DA1-ORN axons (green) and DA1-PN dendrites (magenta) (G), along with the image showing only DA1-PN dendrites and neuropil marker (G’). (H) Match indices for (G) in comparison with two other experimental conditions (Figure 1I). (I) Representative confocal images of full-length Ten-m (FL), Ten-m-ΔECD (ΔECD), or Ten-m-ΔICD (ΔICD) expressing S2 cells with plasma membrane permeabilized or non-permeabilized staining, respectively. Detection of the N-terminal V5 tag (magenta) exclusively in permeabilized staining, and the C-terminal FLAG tag (green) in both permeabilized and non-permeabilized conditions, consistent with expected protein localization and orientation on the plasma membrane (intracellular V5 and extracellular FLAG). (J) Schematic of epitope tagging for constructs used in (I). All expression constructs are V5-tagged at the N-termini (magenta) and FLAG-tagged at the C-termini (green). D, dorsal; L, lateral. Dashed white outline, antennal lobe. BRP, Bruchpilot, an active zone marker used for general neuropil staining. The Kruskal-Wallis test with Bonferroni post-hoc correction for multiple comparisons was used in (F) and (H). In this and all subsequent figures, * p < 0.05; ** p < 0.01; *** p < 0.001; n.s., not significant.

**Figure S2.**
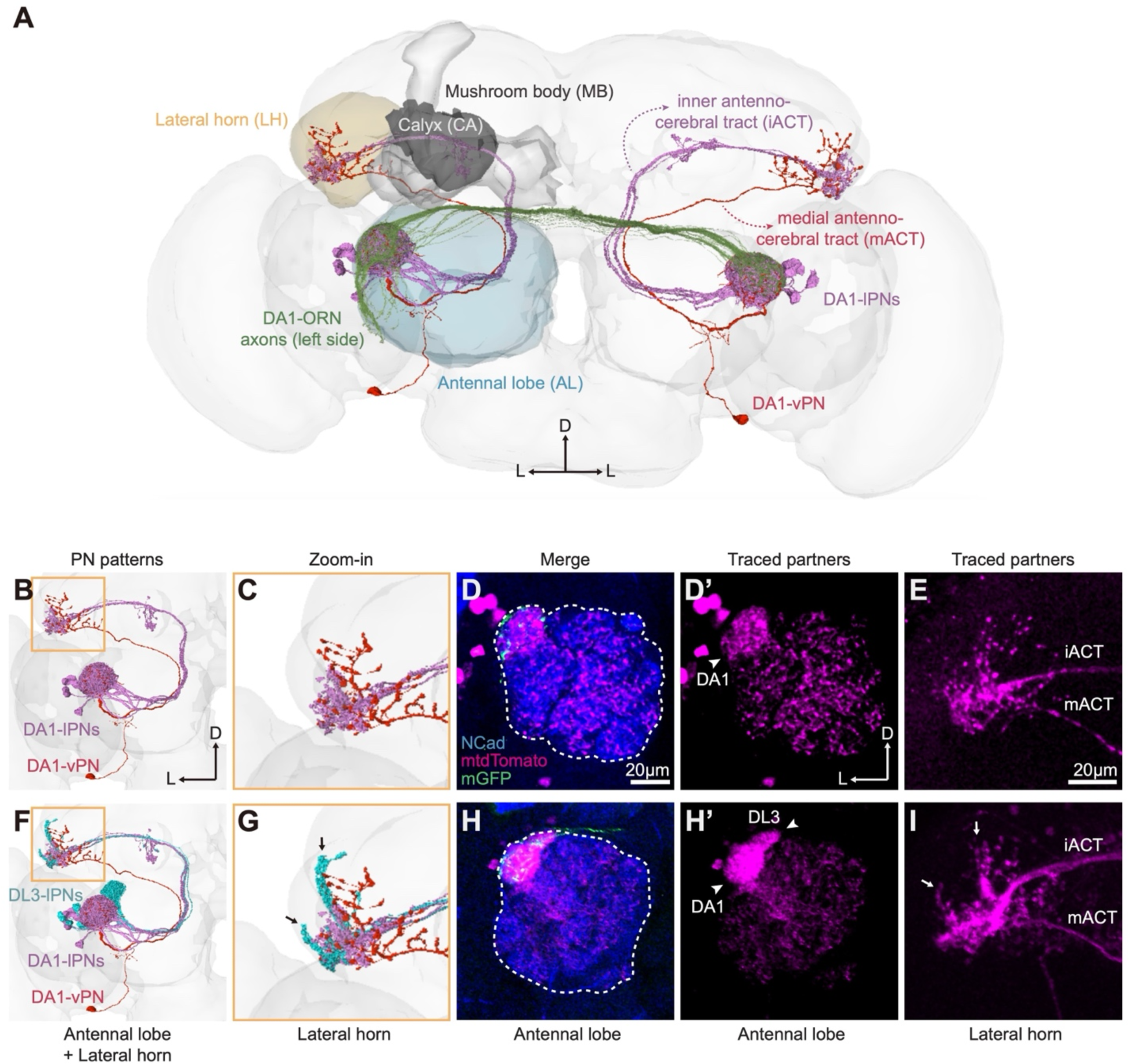
Evidence that Ten-m-overexpressing DA1-ORNs mismatch with DL3-PNs, related to Figure 1. (A) Tracings of DA1-ORNs and two types of DA1-PNs from the FlyWire dataset^80–82^ with relevant brain structures labeled. DA1-ORN axons from the left hemisphere (green) enter the antennal lobe ventrolaterally and innervate the left and right DA1 glomeruli of the antennal lobe. Excitatory DA1-PNs from the lateral neuroblast lineage (DA1-lPNs, purple) send dendrites to the DA1 glomeruli and axons through the inner antennocerebral tract (iACT) to innervate mushroom body calyx and lateral horn. A pair of GABAergic inhibitory DA1-PNs from the ventral neuroblast lineage (DA1-vPN, red) also send dendrites to the DA1 glomeruli and axons through the middle antennocerebral tract (mACT) to innervate only the lateral horn^136,137^. (B, C) FlyWire tracings of DA1-lPNs and DA1-vPN from the left hemisphere (B), with a magnified view at the lateral horn (yellow box) to visualize their stereotyped axon branching patterns (C). (D, E) Representative confocal images of *trans*-Tango-mediated trans-synaptic tracing from DA1-PNs of control. Green, ORN axons; magenta, postsynaptic neurons labeled by *trans*-Tango, which include dendrites of local interneurons and more intensely labeled DA1-PNs in the antennal lobe (D, D’) and DA1-PN axons in the lateral horn (E). Representative images from n = 6. (F, G) FlyWire tracings of DA1-PNs (same as B, C) as well as DL3-PNs (cyan) (F), with a magnified view at the lateral horn (yellow box) to visualize their stereotyped axon branching patterns (G). Arrows indicate signature axon branches of DL3-PNs. (H, I) Representative confocal images of *trans*-Tango-mediated trans-synaptic tracing from DA1-ORNs overexpressing Ten-m. Green, ORN axons; magenta, postsynaptic neurons labeled by *trans*-Tango, which includes not only local interneurons and DA1-PNs, but also notably dense labeling in the DL3 glomerulus (H, H’), as well as combined DA1-PN and DL3-PN axons in the lateral horn (I). The axon branching pattern in the lateral horn (I) is consistent with combined DA1-PN and DL3-PN axon pattern from FlyWire tracing (G). Arrows indicate signature axon branches of DL3-PNs (compared to panels E and G). Representative images from n = 8. D, dorsal; L, lateral. Dashed white circle, antennal lobe. NCad, N-cadherin, a general neuropil marker.

**Figure S3.**
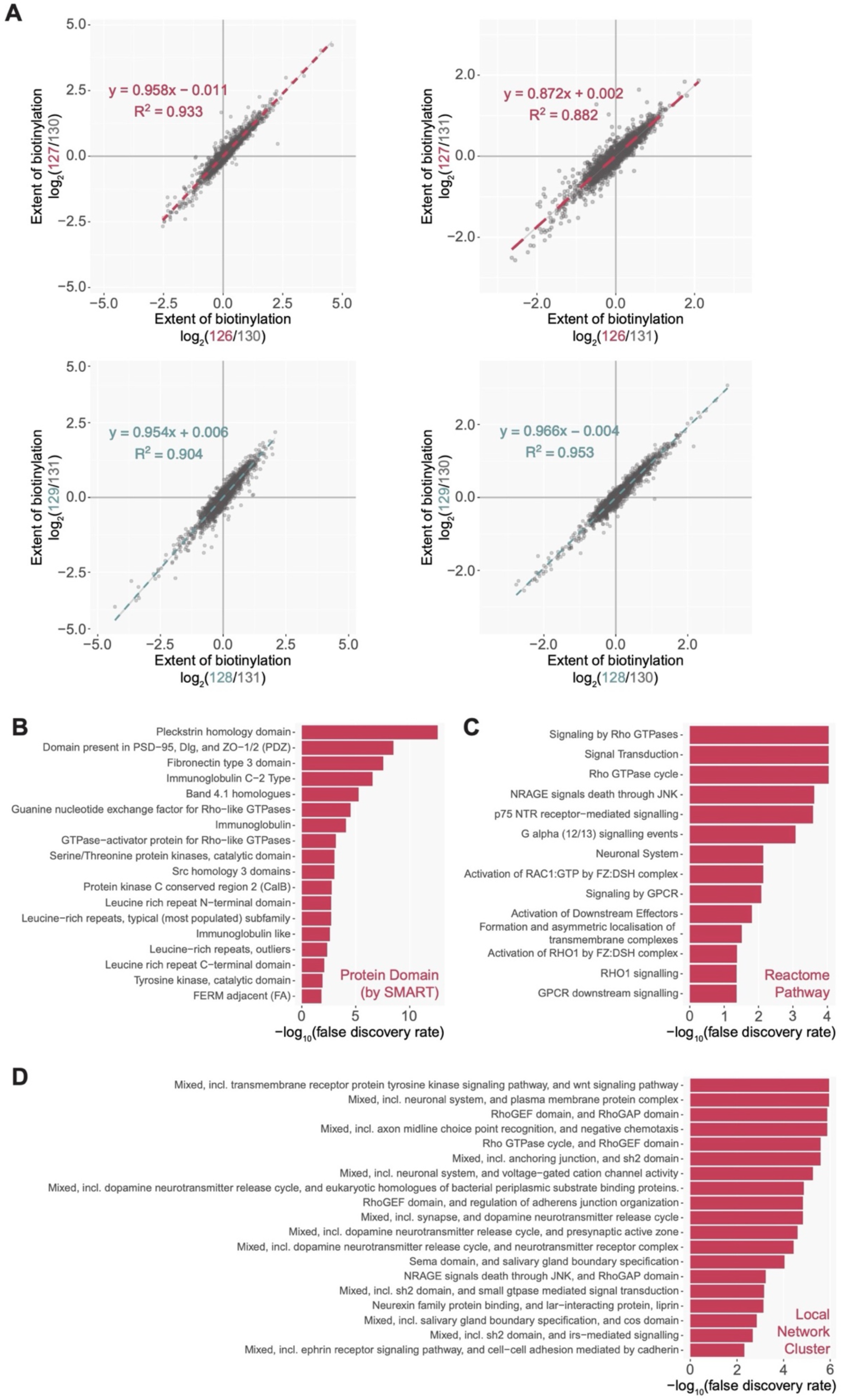
Analysis of unfiltered proteomes and the Ten-m intracellular interactome, related to Figure 2. (A) Correlation of biological replicates. See Figure 2F for the assignment of the TMT labels. (B) Top 18 protein domain terms (predicted by SMART) enriched in the Ten-m intracellular interactome. (C) Top 14 reactome pathway terms enriched in the Ten-m intracellular interactome. (D) Top 19 local network cluster terms enriched in the Ten-m intracellular interactome.

**Figure S4.**
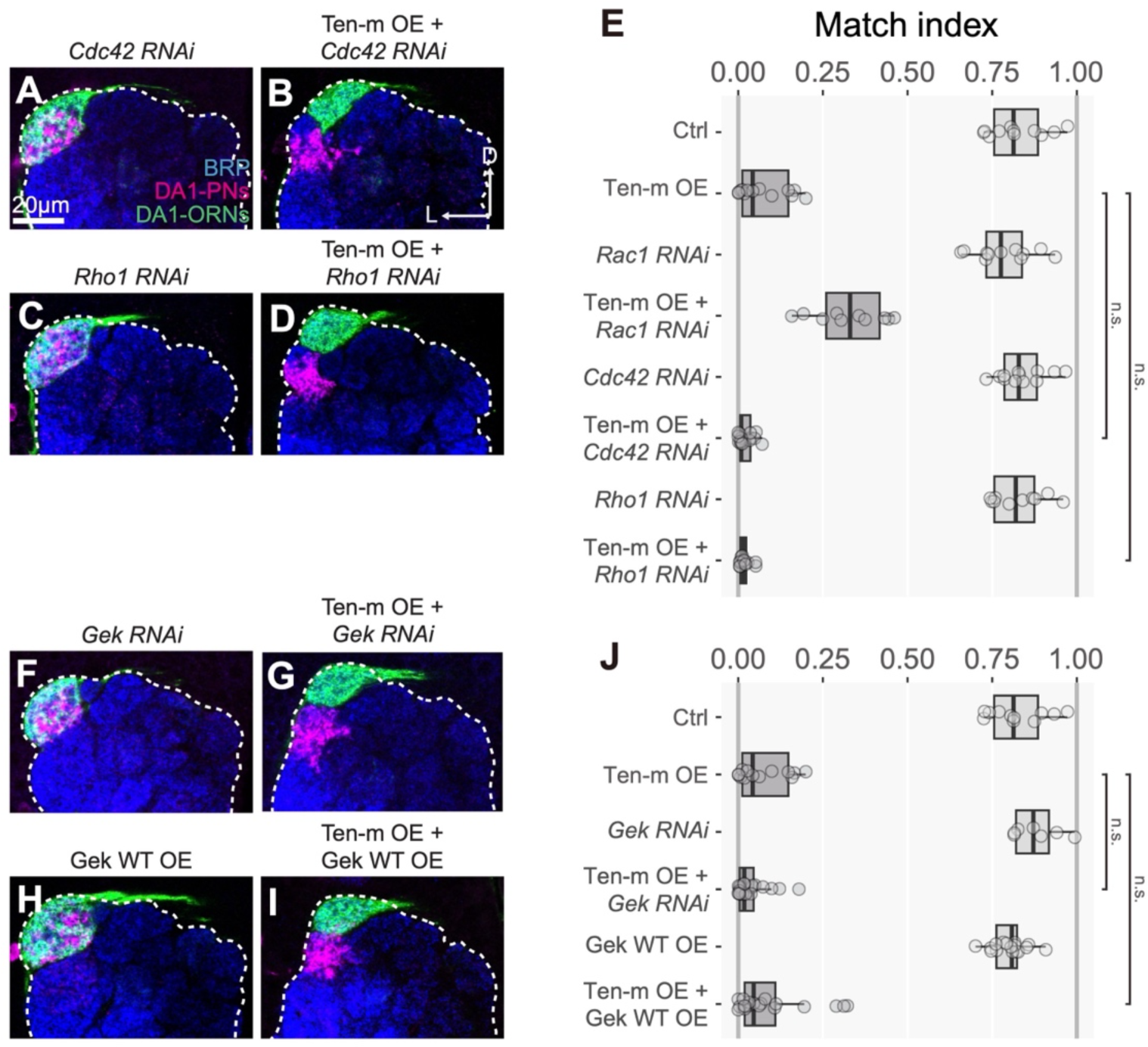
Genetic interaction tests of Ten-m with Cdc42, Rho1, and Gek in ORNs, related to Figures 3 and 4. (A–E) Representative confocal images of DA1-PN dendrites (magenta) and DA1-ORN axons (green) of *Cdc42-RNAi* (A), Ten-m overexpression with *Cdc42-RNAi* (B), *Rho1-RNAi* (C), and Ten-m overexpression with *Rho1-RNAi* (D). Match indices are quantified in (E), which also includes *Rac1-RNAi* (same data as in Figure 3S) for comparison and the control and Ten-m overexpression data from Figure 3I. (F–J) Representative confocal images of DA1-PN dendrites (magenta) and DA1-ORN axons (green) of *Gek-RNAi* (F), Ten-m overexpression with *Gek-RNAi* (G), Gek overexpression (H), and Ten-m and Gek co-overexpression (I). Match indices are quantified in (J), which also includes the control and Ten-m overexpression data from Figure 3I. D, dorsal; L, lateral. Dashed white circle, antennal lobe. BRP, Bruchpilot, an active zone marker used for general neuropil staining. The Kruskal-Wallis test with Bonferroni post-hoc correction for multiple comparisons was used in (J).

**Figure S5.**
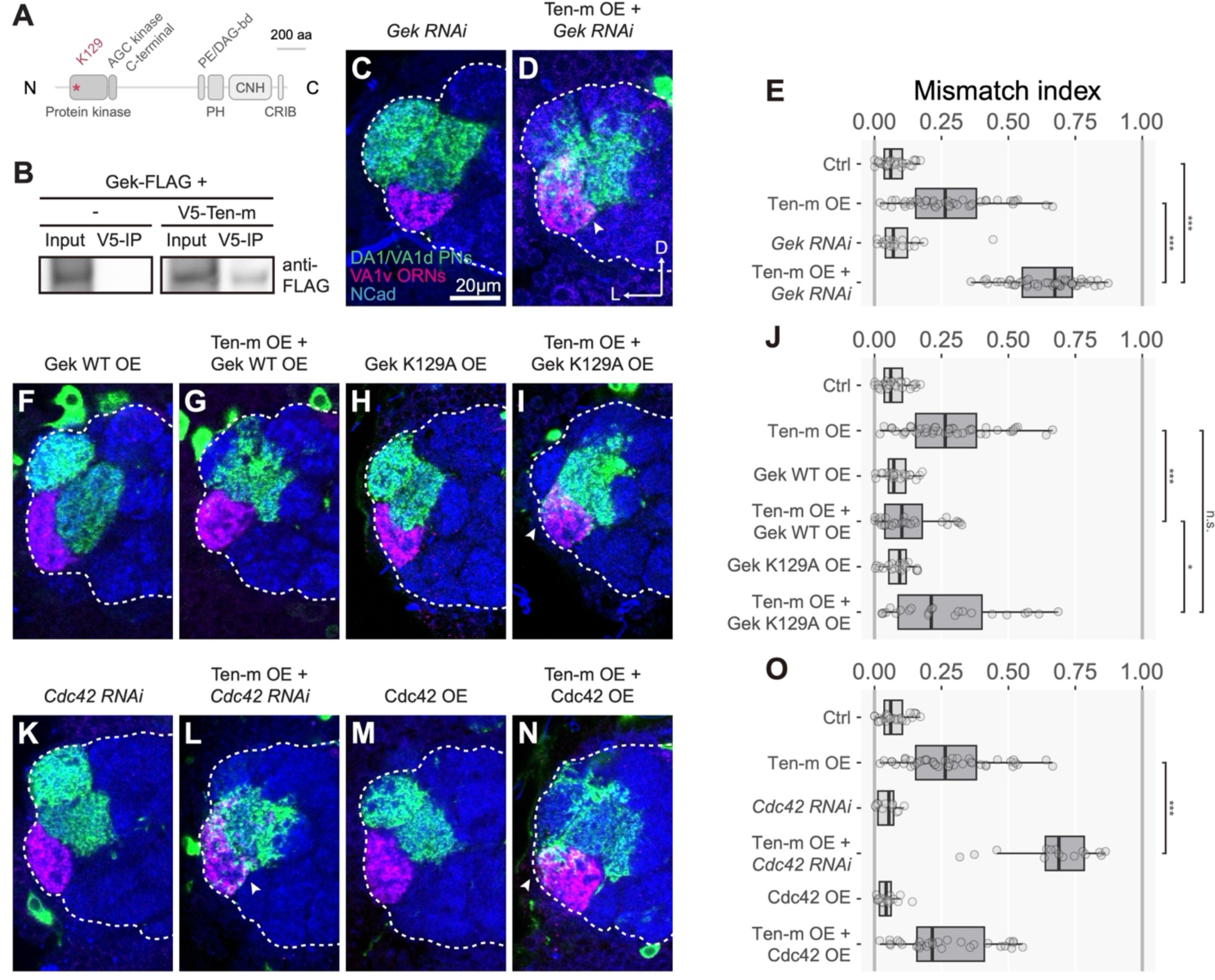
Genetic interactions of Ten-m with Gek and Cdc42 in PNs, related to Figure 4. (A) Protein domain organization of Gek. aa, amino acids. The asterisk (*) marks the lysine in the protein kinase domain essential for its catalytic activity. (B) Co-immunoprecipitation of V5-tagged Ten-m and FLAG-tagged Gek proteins from co-transfected S2 cells. (C–E) Representative confocal images of VA1v-ORN axons (magenta) and Mz19-PN dendrites (green) of *Gek-RNAi* (C) and Ten-m overexpression with *Gek-RNAi* (D). Mismatching indices are quantified in (E), which also includes the control and Ten-m overexpression data from Figure 4G. (F–J) Representative confocal images of VA1v-ORN axons (magenta) and Mz19-PN dendrites (green) of Gek overexpression (F), Gek and Ten-m co-overexpression (G), Gek-K129A protein kinase-domain mutation overexpression (H), and Gek-K129A and Ten-m co-overexpression (I). Mismatch indices are quantified in (J), which also includes the control and Ten-m overexpression data from Figure 4G. (K–O) Representative confocal images of VA1v-ORN axons (magenta) and Mz19-PN dendrites (green) of *Cdc42-RNAi* (K), Ten-m overexpression with *Cdc42-RNAi* (L), Cdc42 overexpression (M), and Cdc42 and Ten-m co-overexpression (N). Mismatching indices are quantified in (O), which also includes the control and Ten-m overexpression data from Figure 4G. D, dorsal; L, lateral. Dashed white circle, antennal lobe. NCad, N-cadherin, a general neuropil marker. Arrowheads indicate overlap regions. The Kruskal-Wallis test with Bonferroni post-hoc correction for multiple comparisons was used in (E), (J), and (O).

**Figure S6.**
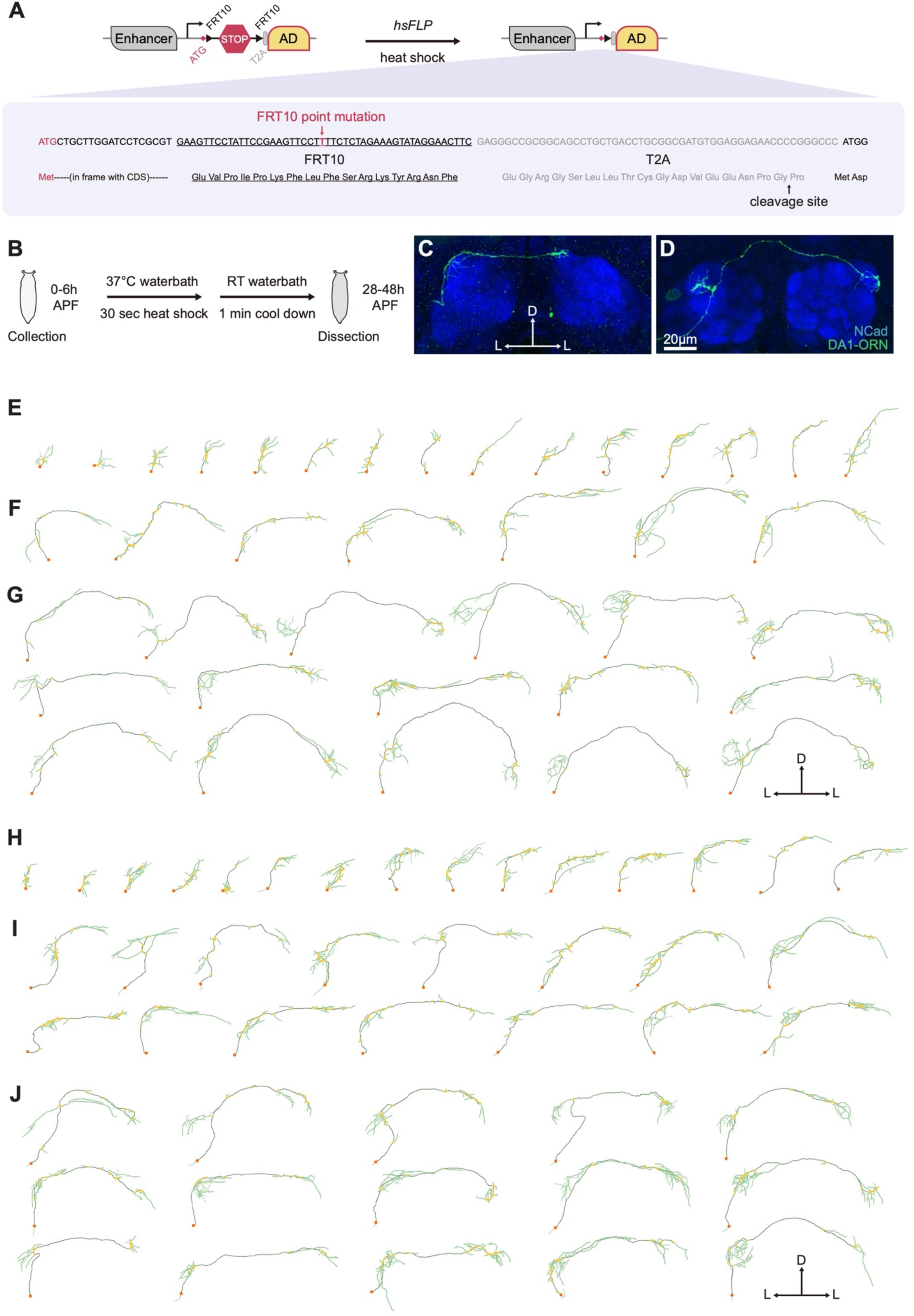
The sparse driver strategy and single-axon analyses, related to Figure 6. (A) The sparse driver strategy incorporates *hsFLP* (FLP recombinase driven by a heat shock promoter), heat shock, and mutant *FRT* (*FRT10*) sites, where the A→T mutation (red) reduces recombination efficiency by 10-fold. Following recombination, the in-frame peptide derived from *FRT10* and *T2A* sequences is excised during the translation of the activation domain (AD). (B) Protocol for activating the sparse driver in a single DA1-ORN. (C, D) Representative maximum Z-projection images of a single DA1-ORN at Stage 2 (C) and Stage 3 (D). (E–G) 3D trace Z-projections of the DA1-ORN axons of control at Stage 1 (E), Stage 2 (F), and Stage 3 (G). (H–J) 3D trace Z-projections of the DA1-ORN axons of Ten-m overexpression at Stage 1 (H), Stage 2 (I), and Stage 3 (J). (K–M) 3D trace Z-projections of the DA1-ORN axons of Ten-m overexpression with *Rac1-RNAi* at Stage 1 (K), Stage 2 (L), and Stage 3 (M). D, dorsal; L, lateral. Orange square, axon entry point. Dark green, stem axon. Light green, axon branches. Yellow dot, primary branch point.

**Figure S7.**
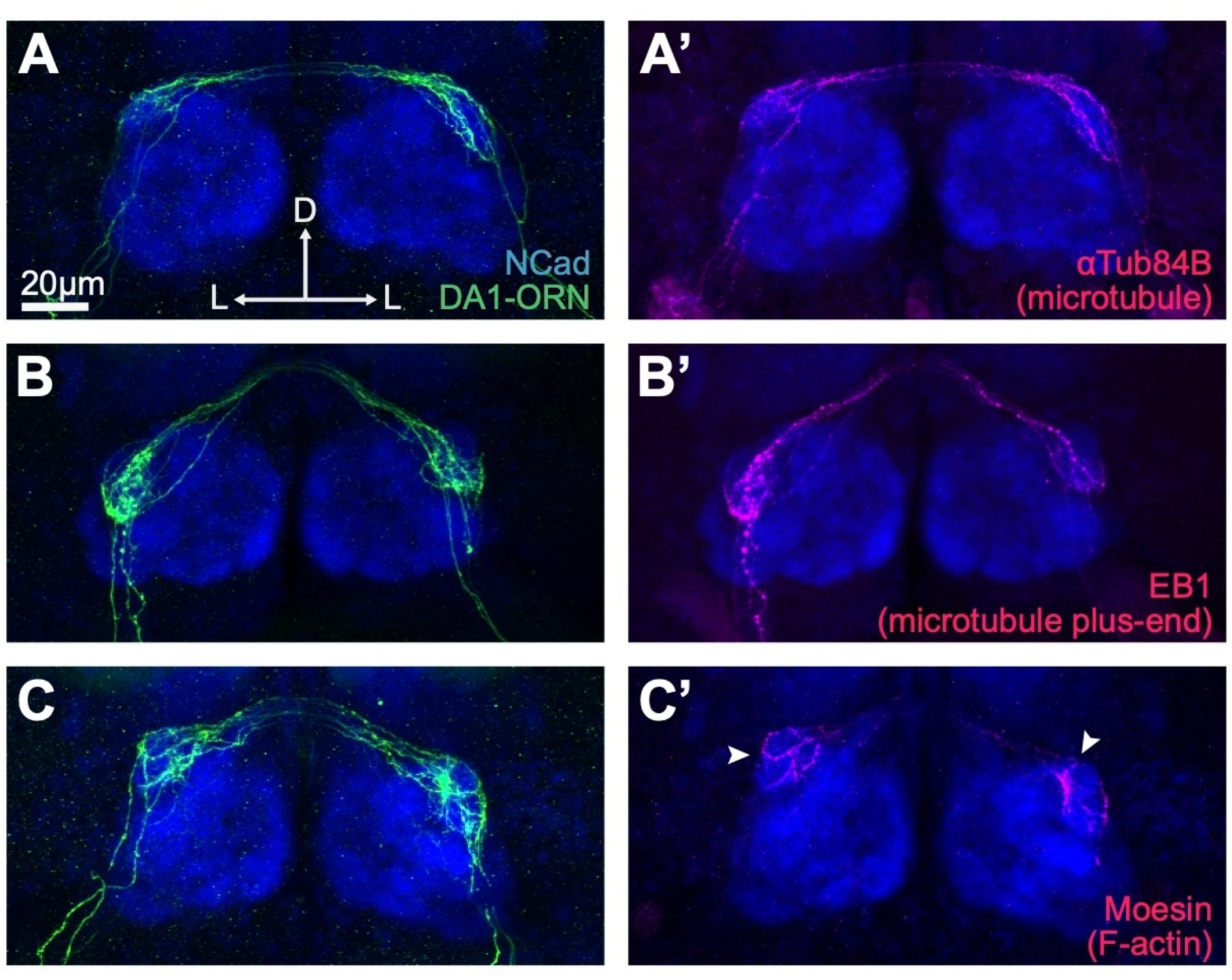
Localizations of cytoskeleton markers in sparsely labeled developing ORN axons, related to Figure 7. Representative maximum Z-projection images of sparse DA1-ORN axons with microtubule marker Halo-alphaTub84B (A and A’), microtubule plus-end marker Halo-EB1 (B and B’), or F-actin marker Halo-Moesin (C and C’). Arrowheads indicate signal peaks of differential distribution of the F-actin marker. D, dorsal; L, lateral. NCad, N-cadherin, a general neuropil marker.

**Table S1.** Processed proteomic data, related to Figure 2. (A) 3454 proteins detected in total. (B) 2854 proteins with 2 or more unique peptides detected, corresponding to **Figure 2I**, Step 1. (C) 781 proteins with [APEX2-Ten-m/NC] fold change higher than or equal to Ten-m’s [APEX2-Ten-m/NC] fold change, corresponding to **Figure 2I**, Step 2. (D) 294 proteins with [APEX2-Ten-m/SR] fold change larger than 0, corresponding to **Figure 2I**, Step 3. *See separate Excel Spreadsheets*

**Table S2.**
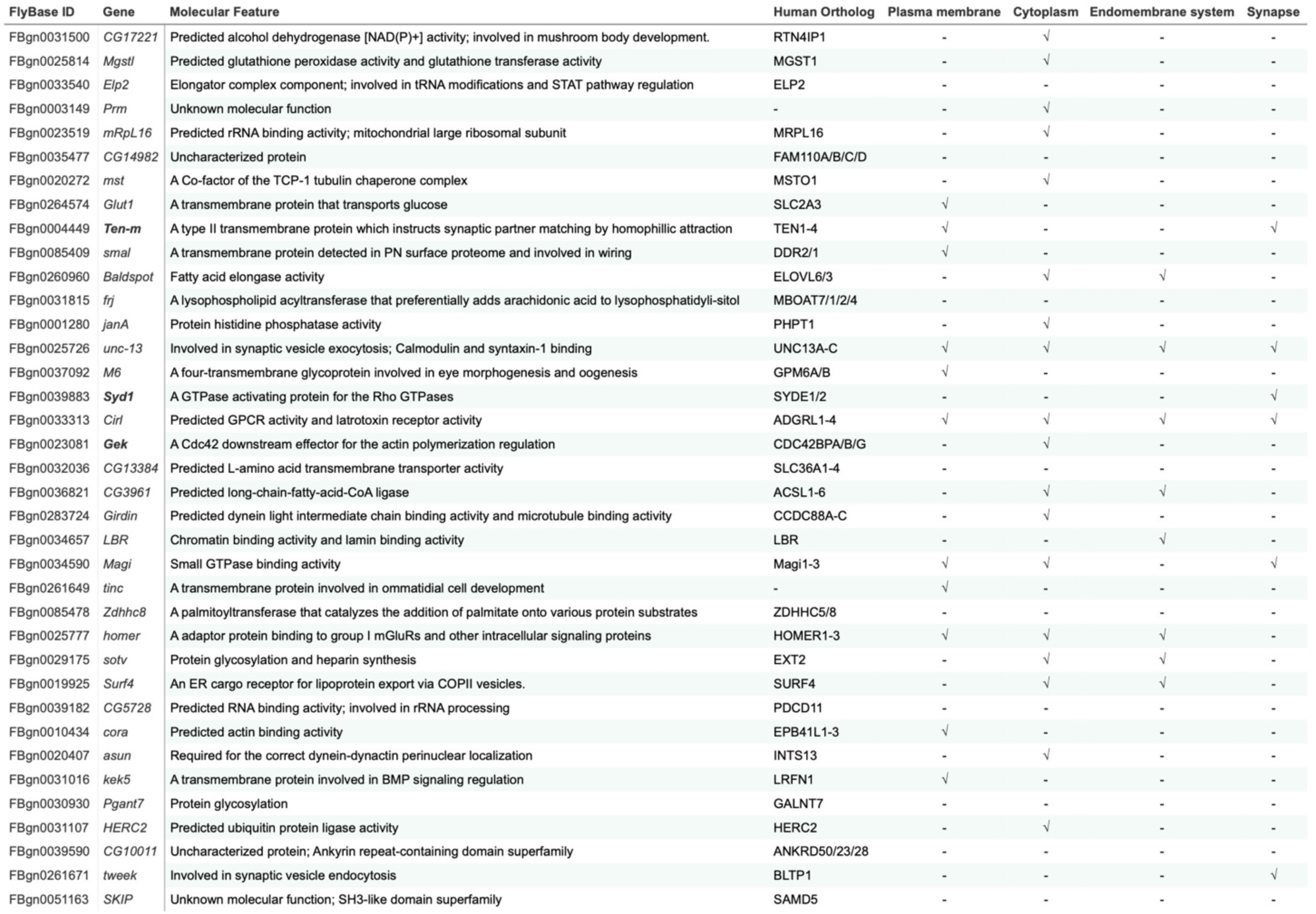
Top candidates in the Ten-m intracellular interactome and their molecular features, related to Figure 2 and Figure 3. Top 37 genes ranked by [APEX2-Ten-m/SR] fold change from the filtered Ten-m intracellular interactome (see Figure 3A red box). Human orthologs were identified using the FlyBase Homologs search tool, listing only those consistently recognized by four or more databases. Molecular features were referenced from FlyBase and UniProt.

**Table S3.**
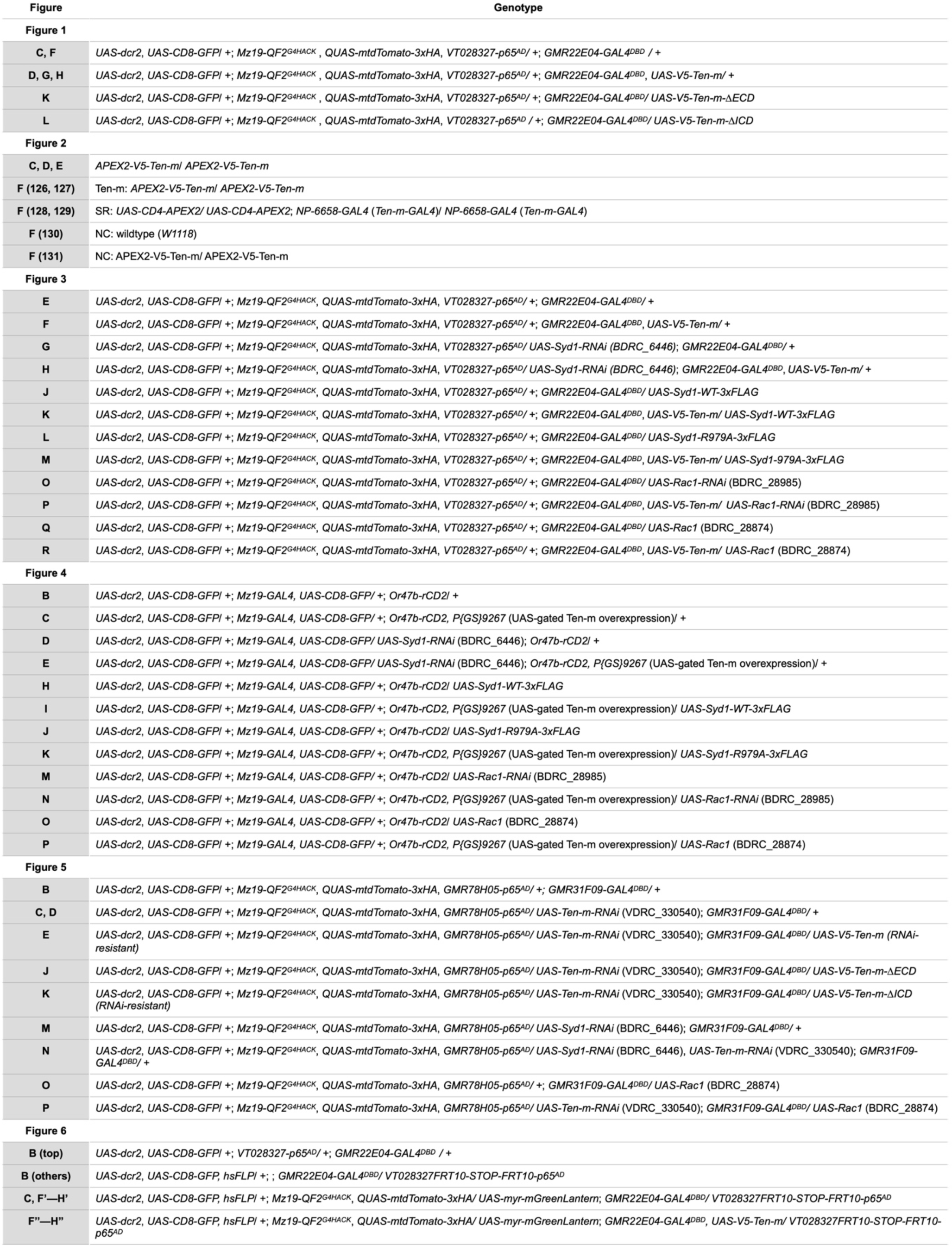
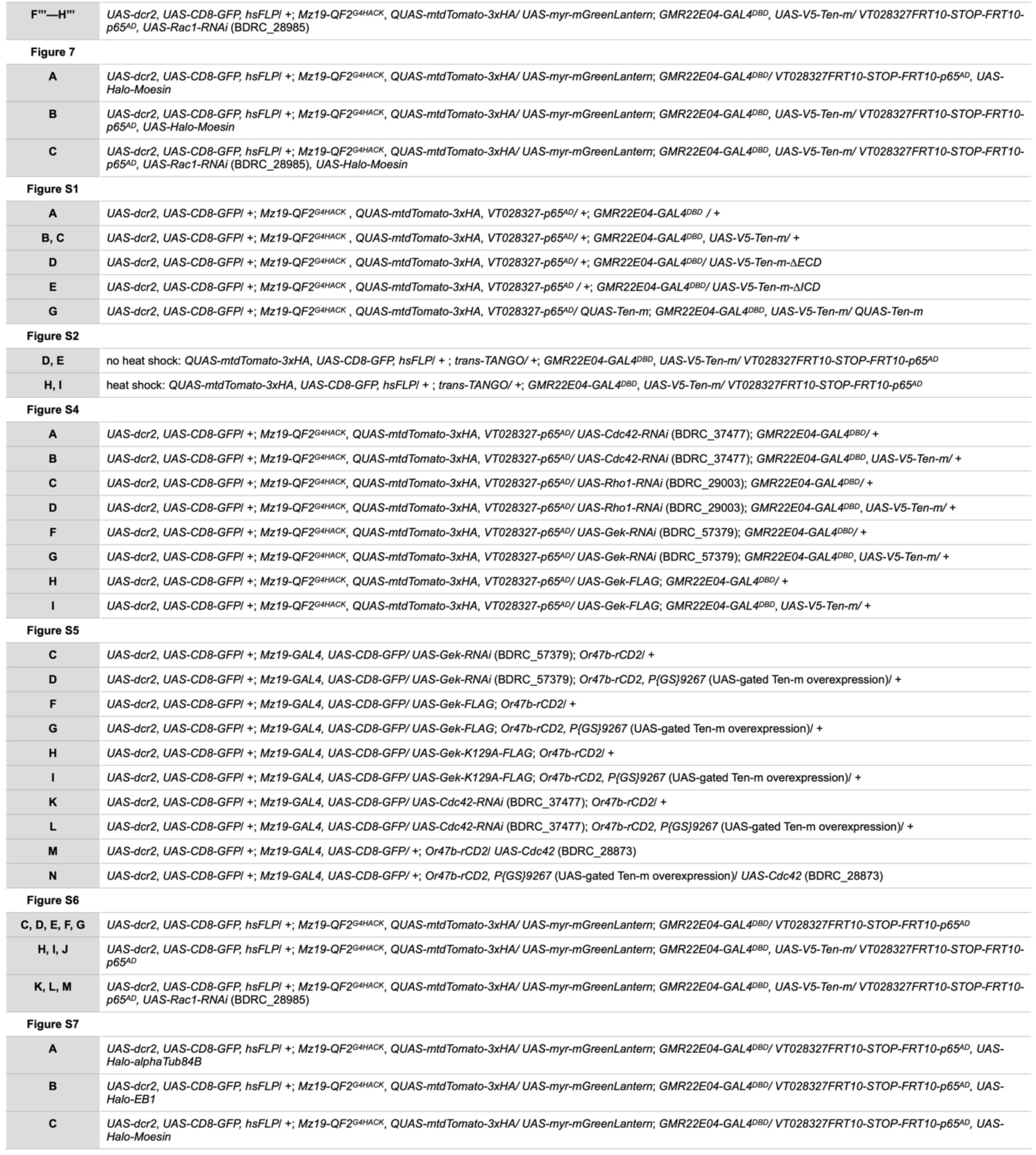
Complete genotypes of each experiment, related to STAR Methods.

## REFERENCES

1. Sanes, J.R., and Yamagata, M. (2009). Many Paths to Synaptic Specificity. Annu. Rev. Cell Dev. Biol. 25, 161–195. 10.1146/annurev.cellbio.24.110707.175402.

2. Jan, Y.-N., and Jan, L.Y. (2010). Branching out: mechanisms of dendritic arborization. Nat Rev Neurosci 11, 316–328. 10.1038/nrn2836.

3. Kolodkin, A.L., and Tessier-Lavigne, M. (2011). Mechanisms and Molecules of Neuronal Wiring: A Primer. Cold Spring Harbor Perspectives in Biology 3, a001727–a001727. 10.1101/cshperspect.a001727.

4. Sanes, J.R., and Zipursky, S.L. (2020). Synaptic Specificity, Recognition Molecules, and Assembly of Neural Circuits. Cell 181, 536–556. 10.1016/j.cell.2020.04.008.

5. Hong, W., Mosca, T.J., and Luo, L. (2012). Teneurins instruct synaptic partner matching in an olfactory map. Nature 484, 201–207. 10.1038/nature10926.

6. Mosca, T.J., Hong, W., Dani, V.S., Favaloro, V., and Luo, L. (2012). Trans-synaptic Teneurin signalling in neuromuscular synapse organization and target choice. Nature 484, 237–241. 10.1038/nature10923.

7. Paré, A.C., Naik, P., Shi, J., Mirman, Z., Palmquist, K.H., and Zallen, J.A. (2019). An LRR Receptor-Teneurin System Directs Planar Polarity at Compartment Boundaries. Developmental Cell 51, 208–221.e6. 10.1016/j.devcel.2019.08.003.

8. Trzebiatowska, A., Topf, U., Sauder, U., Drabikowski, K., and Chiquet-Ehrismann, R. (2008). *Caenorhabditis elegans* Teneurin, *ten-1*, Is Required for Gonadal and Pharyngeal Basement Membrane Integrity and Acts Redundantly with Integrin ina-1 and Dystroglycan *dgn-1*. MBoC 19, 3898–3908. 10.1091/mbc.e08-01-0028.

9. Del Toro, D., Carrasquero-Ordaz, M.A., Chu, A., Ruff, T., Shahin, M., Jackson, V.A., Chavent, M., Berbeira-Santana, M., Seyit-Bremer, G., Brignani, S., et al. (2020). Structural Basis of Teneurin-Latrophilin Interaction in Repulsive Guidance of Migrating Neurons. Cell 180, 323–339.e19. 10.1016/j.cell.2019.12.014.

10. Leamey, C.A., Merlin, S., Lattouf, P., Sawatari, A., Zhou, X., Demel, N., Glendining, K.A., Oohashi, T., Sur, M., and Fässler, R. (2007). Ten_m3 Regulates Eye-Specific Patterning in the Mammalian Visual Pathway and Is Required for Binocular Vision. PLoS Biol 5, e241. 10.1371/journal.pbio.0050241.

11. Zheng, L., Michelson, Y., Freger, V., Avraham, Z., Venken, K.J.T., Bellen, H.J., Justice, M.J., and Wides, R. (2011). Drosophila Ten-m and Filamin Affect Motor Neuron Growth Cone Guidance. PLoS ONE 6, e22956. 10.1371/journal.pone.0022956.

12. Hunyara, J.L., Daly, K.M., Torres, K., Yurgel, M.E., Komal, R., Hattar, S., and Kolodkin, A.L. (2023). Teneurin-3 regulates the generation of non-image-forming visual circuitry and responsiveness to light in the suprachiasmatic nucleus. PLoS Biol 21, e3002412. 10.1371/journal.pbio.3002412.

13. Berns, D.S., DeNardo, L.A., Pederick, D.T., and Luo, L. (2018). Teneurin-3 controls topographic circuit assembly in the hippocampus. Nature 554, 328–333. 10.1038/nature25463.

14. Pederick, D.T., Lui, J.H., Gingrich, E.C., Xu, C., Wagner, M.J., Liu, Y., He, Z., Quake, S.R., and Luo, L. (2021). Reciprocal repulsions instruct the precise assembly of parallel hippocampal networks.

15. Sando, R., Jiang, X., and Südhof, T.C. (2019). Latrophilin GPCRs direct synapse specificity by coincident binding of FLRTs and teneurins.

16. Mosca, T.J., and Luo, L. (2014). Synaptic organization of the Drosophila antennal lobe and its regulation by the Teneurins. eLife 3, e03726. 10.7554/eLife.03726.

17. Suzuki, N., Fukushi, M., Kosaki, K., Doyle, A.D., De Vega, S., Yoshizaki, K., Akazawa, C., Arikawa-Hirasawa, E., and Yamada, Y. (2012). Teneurin-4 Is a Novel Regulator of Oligodendrocyte Differentiation and Myelination of Small-Diameter Axons in the CNS. J. Neurosci. 32, 11586–11599. 10.1523/JNEUROSCI.2045-11.2012.

18. Alkelai, A., Olender, T., Haffner-Krausz, R., Tsoory, M. m., Boyko, V., Tatarskyy, P., Gross-Isseroff, R., Milgrom, R., Shushan, S., Blau, I., et al. (2016). A role for TENM1 mutations in congenital general anosmia. Clinical Genetics 90, 211–219. 10.1111/cge.12782.

19. Aldahmesh, M.A., Mohammed, J.Y., Al-Hazzaa, S., and Alkuraya, F.S. (2012). Homozygous null mutation in ODZ3 causes microphthalmia in humans. Genet Med 14, 900–904. 10.1038/gim.2012.71.

20. Singh, B., Srivastava, P., and Phadke, S.R. (2019). Sequence variations in TENM3 gene causing eye anomalies with intellectual disability: Expanding the phenotypic spectrum. Eur J Med Genet 62, 61–64. 10.1016/j.ejmg.2018.05.004.

21. Hor, H., Francescatto, L., Bartesaghi, L., Ortega-Cubero, S., Kousi, M., Lorenzo-Betancor, O., Jiménez-Jiménez, F.J., Gironell, A., Clarimón, J., Drechsel, O., et al. (2015). Missense mutations in TENM4, a regulator of axon guidance and central myelination, cause essential tremor. Hum Mol Genet 24, 5677–5686. 10.1093/hmg/ddv281.

22. Nava, C., Lamari, F., Héron, D., Mignot, C., Rastetter, A., Keren, B., Cohen, D., Faudet, A., Bouteiller, D., Gilleron, M., et al. (2012). Analysis of the chromosome X exome in patients with autism spectrum disorders identified novel candidate genes, including TMLHE. Transl Psychiatry 2, e179–e179. 10.1038/tp.2012.102.

23. Yi, X., Li, M., He, G., Du, H., Li, X., Cao, D., Wang, L., Wu, X., Yang, F., Chen, X., et al. (2021). Genetic and functional analysis reveals TENM4 contributes to schizophrenia. iScience 24, 103063. 10.1016/j.isci.2021.103063.

24. Psychiatric GWAS Consortium Bipolar Disorder Working Group, Sklar, P., Ripke, S., Scott, L.J., Andreassen, O.A., Cichon, S., Craddock, N., Edenberg, H.J., Nurnberger, J.I., Rietschel, M., et al. (2011). Large-scale genome-wide association analysis of bipolar disorder identifies a new susceptibility locus near ODZ4. Nat Genet 43, 977–983. 10.1038/ng.943.

25. Rebolledo-Jaramillo, B., and Ziegler, A. (2018). Teneurins: An Integrative Molecular, Functional, and Biomedical Overview of Their Role in Cancer. Front. Neurosci. 12, 937. 10.3389/fnins.2018.00937.

26. Peppino, G., Ruiu, R., Arigoni, M., Riccardo, F., Iacoviello, A., Barutello, G., and Quaglino, E. (2021). Teneurins: Role in Cancer and Potential Role as Diagnostic Biomarkers and Targets for Therapy. IJMS 22, 2321. 10.3390/ijms22052321.

27. Pederick, D.T., and Luo, L. (2021). Teneurins. Current Biology 31, R936–R937. 10.1016/j.cub.2021.06.035.

28. Silva, J.-P., Lelianova, V.G., Ermolyuk, Y.S., Vysokov, N., Hitchen, P.G., Berninghausen, O., Rahman, M.A., Zangrandi, A., Fidalgo, S., Tonevitsky, A.G., et al. (2011). Latrophilin 1 and its endogenous ligand Lasso/teneurin-2 form a high-affinity transsynaptic receptor pair with signaling capabilities. Proc. Natl. Acad. Sci. U.S.A. 108, 12113–12118. 10.1073/pnas.1019434108.

29. O’Sullivan, M.L., de Wit, J., Savas, J.N., Comoletti, D., Otto-Hitt, S., Yates, J.R., and Ghosh, A. (2012). FLRT Proteins Are Endogenous Latrophilin Ligands and Regulate Excitatory Synapse Development. Neuron 73, 903–910. 10.1016/j.neuron.2012.01.018.

30. Boucard, A.A., Maxeiner, S., and Südhof, T.C. (2014). Latrophilins Function as Heterophilic Cell-adhesion Molecules by Binding to Teneurins. Journal of Biological Chemistry 289, 387–402. 10.1074/jbc.M113.504779.

31. Li, J., Shalev-Benami, M., Sando, R., Jiang, X., Kibrom, A., Wang, J., Leon, K., Katanski, C., Nazarko, O., Lu, Y.C., et al. (2018). Structural Basis for Teneurin Function in Circuit-Wiring: A Toxin Motif at the Synapse. Cell 173, 735–748.e15. 10.1016/j.cell.2018.03.036.

32. Jackson, V.A., Meijer, D.H., Carrasquero, M., Van Bezouwen, L.S., Lowe, E.D., Kleanthous, C., Janssen, B.J.C., and Seiradake, E. (2018). Structures of Teneurin adhesion receptors reveal an ancient fold for cell-cell interaction. Nat Commun 9, 1079. 10.1038/s41467-018-03460-0.

33. Li, J., Xie, Y., Cornelius, S., Jiang, X., Sando, R., Kordon, S.P., Pan, M., Leon, K., Südhof, T.C., Zhao, M., et al. (2020). Alternative splicing controls teneurin-latrophilin interaction and synapse specificity by a shape-shifting mechanism. Nat Commun 11, 2140. 10.1038/s41467-020-16029-7.

34. Meijer, D.H., Frias, C.P., Beugelink, J.W., Deurloo, Y.N., and Janssen, B.J.C. (2022). Teneurin4 dimer structures reveal a calcium-stabilized compact conformation supporting homomeric trans-interactions. The EMBO Journal 41, e107505. 10.15252/embj.2020107505.

35. Li, J., Bandekar, S.J., and Araç, D. (2023). The structure of fly Teneurin-m reveals an asymmetric self-assembly that allows expansion into zippers. EMBO Reports 24, e56728. 10.15252/embr.202256728.

36. Gogou, C., Beugelink, J.W., Frias, C.P., Kresik, L., Jaroszynska, N., Drescher, U., Janssen, B.J.C., Hindges, R., and Meijer, D.H. (2023). Alternative splicing controls teneurin-3 compact dimer formation for neuronal recognition. Preprint at bioRxiv, 10.1101/2023.10.27.564434 10.1101/2023.10.27.564434.

37. Pederick, D.T., Perry-Hauser, N.A., Meng, H., He, Z., Javitch, J.A., and Luo, L. (2023). Context-dependent requirement of G protein coupling for Latrophilin-2 in target selection of hippocampal axons. eLife 12, e83529. 10.7554/eLife.83529.

38. Jefferis, G.S.X.E., Vyas, R.M., Berdnik, D., Ramaekers, A., Stocker, R.F., Tanaka, N.K., Ito, K., and Luo, L. (2004). Developmental origin of wiring specificity in the olfactory system of *Drosophila*. Development 131, 117–130. 10.1242/dev.00896.

39. Wong, K.K.L., Li, T., Fu, T.-M., Liu, G., Lyu, C., Kohani, S., Xie, Q., Luginbuhl, D.J., Upadhyayula, S., Betzig, E., et al. (2023). Origin of wiring specificity in an olfactory map revealed by neuron type-specific, time-lapse imaging of dendrite targeting. Elife 12, e85521. 10.7554/eLife.85521.

40. Brand, A.H., Manoukian, A.S., and Perrimon, N. (1994). Chapter 33 Ectopic Expression in Drosophila. In Methods in Cell Biology, L. S. B. Goldstein and E. A. Fyrberg, eds. (Academic Press), pp. 635–654. 10.1016/S0091-679X(08)60936-X.

41. Duffy, J.B. (2002). GAL4 system in Drosophila: a fly geneticist’s Swiss army knife. Genesis 34, 1–15. 10.1002/gene.10150.

42. Talay, M., Richman, E.B., Snell, N.J., Hartmann, G.G., Fisher, J.D., Sorkaç, A., Santoyo, J.F., Chou-Freed, C., Nair, N., Johnson, M., et al. (2017). Transsynaptic Mapping of Second-Order Taste Neurons in Flies by trans-Tango. Neuron 96, 783–795.e4. 10.1016/j.neuron.2017.10.011.

43. Han, S., Li, J., and Ting, A.Y. (2018). Proximity labeling: spatially resolved proteomic mapping for neurobiology. Current Opinion in Neurobiology 50, 17–23. 10.1016/j.conb.2017.10.015.

44. Rhee, H.-W., Zou, P., Udeshi, N.D., Martell, J.D., Mootha, V.K., Carr, S.A., and Ting, A.Y. (2013). Proteomic Mapping of Mitochondria in Living Cells via Spatially Restricted Enzymatic Tagging. Science 339, 1328–1331. 10.1126/science.1230593.

45. Lam, S.S., Martell, J.D., Kamer, K.J., Deerinck, T.J., Ellisman, M.H., Mootha, V.K., and Ting, A.Y. (2015). Directed evolution of APEX2 for electron microscopy and proximity labeling. Nat Methods 12, 51–54. 10.1038/nmeth.3179.

46. Li, J., Han, S., Li, H., Udeshi, N.D., Svinkina, T., Mani, D.R., Xu, C., Guajardo, R., Xie, Q., Li, T., et al. (2020). Cell-Surface Proteomic Profiling in the Fly Brain Uncovers Wiring Regulators. Cell 180, 373–386.e15. 10.1016/j.cell.2019.12.029.

47. Xie, Q., Li, J., Li, H., Udeshi, N.D., Svinkina, T., Orlin, D., Kohani, S., Guajardo, R., Mani, D.R., Xu, C., et al. (2022). Transcription factor Acj6 controls dendrite targeting via a combinatorial cell-surface code. Neuron 110, 2299–2314.e8. 10.1016/j.neuron.2022.04.026.

48. Klein, R., and Kania, A. (2014). Ephrin signalling in the developing nervous system. Current Opinion in Neurobiology 27, 16–24. 10.1016/j.conb.2014.02.006.

49. Kania, A., and Klein, R. (2016). Mechanisms of ephrin–Eph signalling in development, physiology and disease. Nat Rev Mol Cell Biol 17, 240–256. 10.1038/nrm.2015.16.

50. Koropouli, E., and Kolodkin, A.L. (2014). Semaphorins and the dynamic regulation of synapse assembly, refinement, and function. Curr Opin Neurobiol 27, 1–7. 10.1016/j.conb.2014.02.005.

51. Pascoe, H.G., Wang, Y., and Zhang, X. (2015). Structural mechanisms of plexin signaling. Progress in Biophysics and Molecular Biology 118, 161–168. 10.1016/j.pbiomolbio.2015.03.006.

52. Alto, L.T., and Terman, J.R. (2017). Semaphorins and their Signaling Mechanisms. Methods Mol Biol 1493, 1–25. 10.1007/978-1-4939-6448-2_1.

53. Patel, M.R., Lehrman, E.K., Poon, V.Y., Crump, J.G., Zhen, M., Bargmann, C.I., and Shen, K. (2006). Hierarchical assembly of presynaptic components in defined C. elegans synapses. Nat Neurosci 9, 1488–1498. 10.1038/nn1806.

54. Owald, D., Fouquet, W., Schmidt, M., Wichmann, C., Mertel, S., Depner, H., Christiansen, F., Zube, C., Quentin, C., Körner, J., et al. (2010). A Syd-1 homologue regulates pre- and postsynaptic maturation in *Drosophila*. Journal of Cell Biology 188, 565–579. 10.1083/jcb.200908055.

55. Spinner, M.A., Walla, D.A., and Herman, T.G. (2018). *Drosophila* Syd-1 Has RhoGAP Activity That Is Required for Presynaptic Clustering of Bruchpilot/ELKS but Not Neurexin-1. Genetics 208, 705–716. 10.1534/genetics.117.300538.

56. Luo, L. (2002). Actin Cytoskeleton Regulation in Neuronal Morphogenesis and Structural Plasticity. Annu. Rev. Cell Dev. Biol. 18, 601–635. 10.1146/annurev.cellbio.18.031802.150501.

57. Ng, J., Nardine, T., Harms, M., Tzu, J., Goldstein, A., Sun, Y., Dietzl, G., Dickson, B.J., and Luo, L. (2002). Rac GTPases control axon growth, guidance and branching. Nature 416, 442–447. 10.1038/416442a.

58. Hakeda-Suzuki, S., Ng, J., Tzu, J., Dietzl, G., Sun, Y., Harms, M., Nardine, T., Luo, L., and Dickson, B.J. (2002). Rac function and regulation during Drosophila development. Nature 416, 438–442. 10.1038/416438a.

59. Simon, M.A., Bowtell, D.D., Dodson, G.S., Laverty, T.R., and Rubin, G.M. (1991). Ras1 and a putative guanine nucleotide exchange factor perform crucial steps in signaling by the sevenless protein tyrosine kinase. Cell 67, 701–716. 10.1016/0092-8674(91)90065-7.

60. Bateman, J., Shu, H., and Van Vactor, D. (2000). The Guanine Nucleotide Exchange Factor Trio Mediates Axonal Development in the Drosophila Embryo. Neuron 26, 93–106. 10.1016/S0896-6273(00)81141-1.

61. Ng, J., and Luo, L. (2004). Rho GTPases Regulate Axon Growth through Convergent and Divergent Signaling Pathways. Neuron 44, 779–793. 10.1016/j.neuron.2004.11.014.

62. Luo, L., Lee, T., Tsai, L., Tang, G., Jan, L.Y., and Jan, Y.N. (1997). Genghis Khan (Gek) as a putative effector for *Drosophila* Cdc42 and regulator of actin polymerization. Proc. Natl. Acad. Sci. U.S.A. 94, 12963–12968. 10.1073/pnas.94.24.12963.

63. Gontang, A.C., Hwa, J.J., Mast, J.D., Schwabe, T., and Clandinin, T.R. (2011). The cytoskeletal regulator Genghis khan is required for columnar target specificity in the *Drosophila* visual system. Development 138, 4899–4909. 10.1242/dev.069930.

64. Li, T., Fu, T.-M., Wong, K.K.L., Li, H., Xie, Q., Luginbuhl, D.J., Wagner, M.J., Betzig, E., and Luo, L. (2021). Cellular bases of olfactory circuit assembly revealed by systematic time-lapse imaging. Cell 184, 5107–5121.e14. 10.1016/j.cell.2021.08.030.

65. Pollard, T.D., Blanchoin, L., and Mullins, R.D. (2000). Molecular Mechanisms Controlling Actin Filament Dynamics in Nonmuscle Cells. Annu. Rev. Biophys. Biomol. Struct. 29, 545–576. 10.1146/annurev.biophys.29.1.545.

66. Dillon, C., and Goda, Y. (2005). THE ACTIN CYTOSKELETON: Integrating Form and Function at the Synapse. Annu. Rev. Neurosci. 28, 25–55. 10.1146/annurev.neuro.28.061604.135757.

67. Lowery, L.A., and Van Vactor, D. (2009). The trip of the tip: understanding the growth cone machinery. Nat Rev Mol Cell Biol 10, 332–343. 10.1038/nrm2679.

68. Dent, E.W., Gupton, S.L., and Gertler, F.B. (2011). The Growth Cone Cytoskeleton in Axon Outgrowth and Guidance. Cold Spring Harbor Perspectives in Biology 3, a001800–a001800. 10.1101/cshperspect.a001800.

69. Grieder, N.C., Cuevas, M. de, and Spradling, A.C. (2000). The fusome organizes the microtubule network during oocyte differentiation in Drosophila. Development 127, 4253–4264. 10.1242/dev.127.19.4253.

70. Rusan, N.M., and Peifer, M. (2007). A role for a novel centrosome cycle in asymmetric cell division. Journal of Cell Biology 177, 13–20. 10.1083/jcb.200612140.

71. Edwards, K.A., Demsky, M., Montague, R.A., Weymouth, N., and Kiehart, D.P. (1997). GFP-Moesin Illuminates Actin Cytoskeleton Dynamics in Living Tissue and Demonstrates Cell Shape Changes during Morphogenesis inDrosophila. Developmental Biology 191, 103–117. 10.1006/dbio.1997.8707.

72. Tosney, K.W., and Landmesser, L.T. (1985). Specificity of early motoneuron growth cone outgrowth in the chick embryo. J Neurosci 5, 2336–2344. 10.1523/JNEUROSCI.05-09-02336.1985.

73. Sanes, J.R., and Lichtman, J.W. (1999). Development of the vertebrate neuromuscular junction. Annu Rev Neurosci 22, 389–442. 10.1146/annurev.neuro.22.1.389.

74. Luria, V., Krawchuk, D., Jessell, T.M., Laufer, E., and Kania, A. (2008). Specification of motor axon trajectory by ephrin-B:EphB signaling: symmetrical control of axonal patterning in the developing limb. Neuron 60, 1039–1053. 10.1016/j.neuron.2008.11.011.

75. Chou, V.T., Johnson, S.A., and Van Vactor, D. (2020). Synapse development and maturation at the drosophila neuromuscular junction. Neural Development 15, 11. 10.1186/s13064-020-00147-5.

76. Buffelli, M., Burgess, R.W., Feng, G., Lobe, C.G., Lichtman, J.W., and Sanes, J.R. (2003). Genetic evidence that relative synaptic efficacy biases the outcome of synaptic competition. Nature 424, 430–434. 10.1038/nature01844.

77. Lu, J., Tapia, J.C., White, O.L., and Lichtman, J.W. (2009). The interscutularis muscle connectome. PLoS Biol 7, e32. 10.1371/journal.pbio.1000032.

78. Tapia, J.C., Wylie, J.D., Kasthuri, N., Hayworth, K.J., Schalek, R., Berger, D.R., Guatimosim, C., Seung, H.S., and Lichtman, J.W. (2012). Pervasive synaptic branch removal in the mammalian neuromuscular system at birth. Neuron 74, 816–829. 10.1016/j.neuron.2012.04.017.

79. Kano, M., and Watanabe, T. (2019). Developmental synapse remodeling in the cerebellum and visual thalamus. F1000Res 8, F1000 Faculty Rev-1191. 10.12688/f1000research.18903.1.

80. Zheng, Z., Lauritzen, J.S., Perlman, E., Robinson, C.G., Nichols, M., Milkie, D., Torrens, O., Price, J., Fisher, C.B., Sharifi, N., et al. (2018). A Complete Electron Microscopy Volume of the Brain of Adult Drosophila melanogaster. Cell 174, 730–743.e22. 10.1016/j.cell.2018.06.019.

81. Dorkenwald, S., Matsliah, A., Sterling, A.R., Schlegel, P., Yu, S., McKellar, C.E., Lin, A., Costa, M., Eichler, K., Yin, Y., et al. (2023). Neuronal wiring diagram of an adult brain (Neuroscience) 10.1101/2023.06.27.546656.

82. Schlegel, P., Yin, Y., Bates, A.S., Dorkenwald, S., Eichler, K., Brooks, P., Han, D.S., Gkantia, M., Santos, M. dos, Munnelly, E.J., et al. (2023). Whole-brain annotation and multi-connectome cell typing quantifies circuit stereotypy in Drosophila. Preprint at bioRxiv, 10.1101/2023.06.27.546055 10.1101/2023.06.27.546055.

83. Tobin, W.F., Wilson, R.I., and Lee, W.-C.A. (2017). Wiring variations that enable and constrain neural computation in a sensory microcircuit. eLife 6, e24838. 10.7554/eLife.24838.

84. Joo, W.J., Sweeney, L.B., Liang, L., and Luo, L. (2013). Linking cell fate, trajectory choice, and target selection: genetic analysis of Sema-2b in olfactory axon targeting. Neuron 78, 673–686. 10.1016/j.neuron.2013.03.022.

85. Li, J., Guajardo, R., Xu, C., Wu, B., Li, H., Li, T., Luginbuhl, D.J., Xie, X., and Luo, L. (2018). Stepwise wiring of the Drosophila olfactory map requires specific Plexin B levels. eLife 7, e39088. 10.7554/eLife.39088.

86. Zou, D.-J., Chesler, A., and Firestein, S. (2009). How the olfactory bulb got its glomeruli: a just so story? Nat Rev Neurosci 10, 611–618. 10.1038/nrn2666.

87. Fujimoto, S., Leiwe, M.N., Aihara, S., Sakaguchi, R., Muroyama, Y., Kobayakawa, R., Kobayakawa, K., Saito, T., and Imai, T. (2023). Activity-dependent local protection and lateral inhibition control synaptic competition in developing mitral cells in mice. Developmental Cell 58, 1221–1236.e7. 10.1016/j.devcel.2023.05.004.

88. Chen, Y., Akin, O., Nern, A., Tsui, C.Y.K., Pecot, M.Y., and Zipursky, S.L. (2014). Cell-type-specific labeling of synapses in vivo through synaptic tagging with recombination. Neuron 81, 280–293. 10.1016/j.neuron.2013.12.021.

89. McLaughlin, C.N., Brbić, M., Xie, Q., Li, T., Horns, F., Kolluru, S.S., Kebschull, J.M., Vacek, D., Xie, A., Li, J., et al. (2021). Single-cell transcriptomes of developing and adult olfactory receptor neurons in Drosophila. eLife 10, e63856. 10.7554/eLife.63856.

90. Bashaw, G.J., and Klein, R. (2010). Signaling from Axon Guidance Receptors. Cold Spring Harbor Perspectives in Biology 2, a001941–a001941. 10.1101/cshperspect.a001941.

91. Rosa, M., Noel, T., Harris, M., and Ladds, G. (2021). Emerging roles of adhesion G protein-coupled receptors. Biochemical Society Transactions 49, 1695–1709. 10.1042/BST20201144.

92. Langenhan, T., Piao, X., and Monk, K.R. (2016). Adhesion G protein-coupled receptors in nervous system development and disease. Nat Rev Neurosci 17, 550–561. 10.1038/nrn.2016.86.

93. Sando, R., and Südhof, T.C. (2021). Latrophilin GPCR signaling mediates synapse formation. eLife 10, e65717. 10.7554/eLife.65717.

94. Wang, S., DeLeon, C., Sun, W., Quake, S.R., Roth, B.L., and Südhof, T.C. (2024). Alternative splicing of latrophilin-3 controls synapse formation. Nature, 1–8. 10.1038/s41586-023-06913-9.

95. Song, H., and Poo, M. (2001). The cell biology of neuronal navigation. Nat Cell Biol 3, E81–88. 10.1038/35060164.

96. O’Donnell, M., Chance, R.K., and Bashaw, G.J. (2009). Axon growth and guidance: receptor regulation and signal transduction. Annu Rev Neurosci 32, 383–412. 10.1146/annurev.neuro.051508.135614.

97. Südhof, T.C. (2012). The Presynaptic Active Zone. Neuron 75, 11–25. 10.1016/j.neuron.2012.06.012.

98. Cajal, R. (1995). Histology of the Nervous System of Man and Vertebrates (Oxford University Press).

99. Luo, L. (2007). Fly MARCM and mouse MADM: Genetic methods of labeling and manipulating single neurons. Brain Research Reviews 55, 220–227. 10.1016/j.brainresrev.2007.01.012.

100. Jefferis, G.S., and Livet, J. (2012). Sparse and combinatorial neuron labelling. Current Opinion in Neurobiology 22, 101–110. 10.1016/j.conb.2011.09.010.

101. Lin, R., Wang, R., Yuan, J., Feng, Q., Zhou, Y., Zeng, S., Ren, M., Jiang, S., Ni, H., Zhou, C., et al. (2018). Cell-type-specific and projection-specific brain-wide reconstruction of single neurons. Nat Methods 15, 1033–1036. 10.1038/s41592-018-0184-y.

102. Winnubst, J., Bas, E., Ferreira, T.A., Wu, Z., Economo, M.N., Edson, P., Arthur, B.J., Bruns, C., Rokicki, K., Schauder, D., et al. (2019). Reconstruction of 1,000 Projection Neurons Reveals New Cell Types and Organization of Long-Range Connectivity in the Mouse Brain. Cell 179, 268–281.e13. 10.1016/j.cell.2019.07.042.

103. Peng, H., Xie, P., Liu, L., Kuang, X., Wang, Y., Qu, L., Gong, H., Jiang, S., Li, A., Ruan, Z., et al. (2021). Morphological diversity of single neurons in molecularly defined cell types. Nature 598, 174–181. 10.1038/s41586-021-03941-1.

104. Nern, A., Pfeiffer, B.D., and Rubin, G.M. (2015). Optimized tools for multicolor stochastic labeling reveal diverse stereotyped cell arrangements in the fly visual system. Proceedings of the National Academy of Sciences 112, E2967–E2976. 10.1073/pnas.1506763112.

105. Isaacman-Beck, J., Paik, K.C., Wienecke, C.F.R., Yang, H.H., Fisher, Y.E., Wang, I.E., Ishida, I.G., Maimon, G., Wilson, R.I., and Clandinin, T.R. (2020). SPARC enables genetic manipulation of precise proportions of cells. Nat Neurosci 23, 1168–1175. 10.1038/s41593-020-0668-9.

106. Lee, T., and Luo, L. (1999). Mosaic Analysis with a Repressible Cell Marker for Studies of Gene Function in Neuronal Morphogenesis. Neuron 22, 451–461. 10.1016/S0896-6273(00)80701-1.

107. Wu, J.S., and Luo, L. (2006). A protocol for mosaic analysis with a repressible cell marker (MARCM) in Drosophila. Nat Protoc 1, 2583–2589. 10.1038/nprot.2006.320.

108. Luan, H., Peabody, N.C., Vinson, C.R., and White, B.H. (2006). Refined Spatial Manipulation of Neuronal Function by Combinatorial Restriction of Transgene Expression. Neuron 52, 425–436. 10.1016/j.neuron.2006.08.028.

109. Ting, C.-Y., Gu, S., Guttikonda, S., Lin, T.-Y., White, B.H., and Lee, C.-H. (2011). Focusing Transgene Expression in Drosophila by Coupling Gal4 With a Novel Split-LexA Expression System. Genetics 188, 229–233. 10.1534/genetics.110.126193.

110. Riabinina, O., Vernon, S.W., Dickson, B.J., and Baines, R.A. (2019). Split-QF System for Fine-Tuned Transgene Expression in Drosophila. Genetics 212, 53–63. 10.1534/genetics.119.302034.

111. Jenett, A., Rubin, G.M., Ngo, T.-T.B., Shepherd, D., Murphy, C., Dionne, H., Pfeiffer, B.D., Cavallaro, A., Hall, D., Jeter, J., et al. (2012). A GAL4-Driver Line Resource for Drosophila Neurobiology. Cell Reports 2, 991–1001. 10.1016/j.celrep.2012.09.011.

112. Dietzl, G., Chen, D., Schnorrer, F., Su, K.-C., Barinova, Y., Fellner, M., Gasser, B., Kinsey, K., Oppel, S., Scheiblauer, S., et al. (2007). A genome-wide transgenic RNAi library for conditional gene inactivation in Drosophila. Nature 448, 151–156. 10.1038/nature05954.

113. Dionne, H., Hibbard, K.L., Cavallaro, A., Kao, J.-C., and Rubin, G.M. (2018). Genetic Reagents for Making Split-GAL4 Lines in Drosophila. Genetics 209, 31–35. 10.1534/genetics.118.300682.

114. Ni, J.-Q., Zhou, R., Czech, B., Liu, L.-P., Holderbaum, L., Yang-Zhou, D., Shim, H.-S., Tao, R., Handler, D., Karpowicz, P., et al. (2011). A genome-scale shRNA resource for transgenic RNAi in Drosophila. Nat Methods 8, 405–407. 10.1038/nmeth.1592.

115. Ito, K., Suzuki, K., Estes, P., Ramaswami, M., Yamamoto, D., and Strausfeld, N.J. (1998). The Organization of Extrinsic Neurons and Their Implications in the Functional Roles of the Mushroom Bodies in Drosophila melanogaster Meigen. Learn. Mem. 5, 52–77. 10.1101/lm.5.1.52.

116. Perkins, L.A., Holderbaum, L., Tao, R., Hu, Y., Sopko, R., McCall, K., Yang-Zhou, D., Flockhart, I., Binari, R., Shim, H.-S., et al. (2015). The Transgenic RNAi Project at Harvard Medical School: Resources and Validation. Genetics 201, 843–852. 10.1534/genetics.115.180208.

117. Potter, C.J., Tasic, B., Russler, E.V., Liang, L., and Luo, L. (2010). The Q System: A Repressible Binary System for Transgene Expression, Lineage Tracing, and Mosaic Analysis. Cell 141, 536–548. 10.1016/j.cell.2010.02.025.

118. Gratz, S.J., Ukken, F.P., Rubinstein, C.D., Thiede, G., Donohue, L.K., Cummings, A.M., and O’Connor-Giles, K.M. (2014). Highly Specific and Efficient CRISPR/Cas9-Catalyzed Homology-Directed Repair in *Drosophila*. Genetics 196, 961–971. 10.1534/genetics.113.160713.

119. Zhu, H., and Luo, L. (2004). Diverse Functions of N-Cadherin in Dendritic and Axonal Terminal Arborization of Olfactory Projection Neurons. Neuron 42, 63–75. 10.1016/S0896-6273(04)00142-4.

120. Gratz, S.J., Rubinstein, C.D., Harrison, M.M., Wildonger, J., and O’Connor-Giles, K.M. (2015). CRISPR-Cas9 Genome Editing in *Drosophila*. CP Molecular Biology 111. 10.1002/0471142727.mb3102s111.

121. Bier, E., Harrison, M.M., O’Connor-Giles, K.M., and Wildonger, J. (2018). Advances in Engineering the Fly Genome with the CRISPR-Cas System. Genetics 208, 1–18. 10.1534/genetics.117.1113.

122. Golic, K.G., and Lindquist, S. (1989). The FLP recombinase of yeast catalyzes site-specific recombination in the Drosophila genome. Cell 59, 499–509. 10.1016/0092-8674(89)90033-0.

123. Gratz, S.J., Cummings, A.M., Nguyen, J.N., Hamm, D.C., Donohue, L.K., Harrison, M.M., Wildonger, J., and O’Connor-Giles, K.M. (2013). Genome engineering of Drosophila with the CRISPR RNA-guided Cas9 nuclease. Genetics 194, 1029–1035. 10.1534/genetics.113.152710.

124. Port, F., Chen, H.-M., Lee, T., and Bullock, S.L. (2014). Optimized CRISPR/Cas tools for efficient germline and somatic genome engineering in Drosophila. Proc Natl Acad Sci U S A 111, E2967–2976. 10.1073/pnas.1405500111.

125. Han, C., Jan, L.Y., and Jan, Y.-N. (2011). Enhancer-driven membrane markers for analysis of nonautonomous mechanisms reveal neuron–glia interactions in Drosophila. Proceedings of the National Academy of Sciences 108, 9673–9678. 10.1073/pnas.1106386108.

126. Pfeiffer, B.D., Truman, J.W., and Rubin, G.M. (2012). Using translational enhancers to increase transgene expression in *Drosophila*. Proc. Natl. Acad. Sci. U.S.A. 109, 6626–6631. 10.1073/pnas.1204520109.

127. Tirian, L., and Dickson, B.J. (2017). The VT GAL4, LexA, and split-GAL4 driver line collections for targeted expression in the Drosophila nervous system. Preprint at bioRxiv, 10.1101/198648 10.1101/198648.

128. Benjamini, Y., and Hochberg, Y. (1995). Controlling the False Discovery Rate: A Practical and Powerful Approach to Multiple Testing. Journal of the Royal Statistical Society. Series B (Methodological) 57, 289–300.

129. Ritchie, M.E., Phipson, B., Wu, D., Hu, Y., Law, C.W., Shi, W., and Smyth, G.K. (2015). limma powers differential expression analyses for RNA-sequencing and microarray studies. Nucleic Acids Res 43, e47. 10.1093/nar/gkv007.

130. Wu, J.S., and Luo, L. (2006). A protocol for dissecting Drosophila melanogaster brains for live imaging or immunostaining. Nat Protoc 1, 2110–2115. 10.1038/nprot.2006.336.

131. Kohl, J., Ng, J., Cachero, S., Ciabatti, E., Dolan, M.-J., Sutcliffe, B., Tozer, A., Ruehle, S., Krueger, D., Frechter, S., et al. (2014). Ultrafast tissue staining with chemical tags. Proc. Natl. Acad. Sci. U.S.A. 111. 10.1073/pnas.1411087111.

132. Grimm, J.B., Muthusamy, A.K., Liang, Y., Brown, T.A., Lemon, W.C., Patel, R., Lu, R., Macklin, J.J., Keller, P.J., Ji, N., et al. (2017). A general method to fine-tune fluorophores for live-cell and in vivo imaging. Nat Methods 14, 987–994. 10.1038/nmeth.4403.

133. Grimm, J.B., Xie, L., Casler, J.C., Patel, R., Tkachuk, A.N., Falco, N., Choi, H., Lippincott-Schwartz, J., Brown, T.A., Glick, B.S., et al. (2021). A General Method to Improve Fluorophores Using Deuterated Auxochromes. JACS Au 1, 690–696. 10.1021/jacsau.1c00006.

134. Arshadi, C., Günther, U., Eddison, M., Harrington, K.I.S., and Ferreira, T.A. (2021). SNT: a unifying toolbox for quantification of neuronal anatomy. Nat Methods 18, 374–377. 10.1038/s41592-021-01105-7.

135. Bates, A.S., Manton, J.D., Jagannathan, S.R., Costa, M., Schlegel, P., Rohlfing, T., and Jefferis, G.S. (2020). The natverse, a versatile toolbox for combining and analysing neuroanatomical data. eLife 9, e53350. 10.7554/eLife.53350.

136. Marin, E.C., Jefferis, G.S.X.E., Komiyama, T., Zhu, H., and Luo, L. (2002). Representation of the glomerular olfactory map in the Drosophila brain. Cell 109, 243–255. 10.1016/s0092-8674(02)00700-6.

137. Liang, L., Li, Y., Potter, C.J., Yizhar, O., Deisseroth, K., Tsien, R.W., and Luo, L. (2013). GABAergic Projection Neurons Route Selective Olfactory Inputs to Specific Higher Order Neurons. Neuron 79, 10.1016/j.neuron.2013.06.014. 10.1016/j.neuron.2013.06.014.

